# Gaining or cutting SLAC: the evolution of plant guard cell signalling pathways

**DOI:** 10.1101/2021.05.26.445736

**Authors:** Frances C. Sussmilch, Tobias Maierhofer, Johannes Herrmann, Lena J. Voss, Christof Lind, Maxim Messerer, Heike M. Müller, Maria S. Bünner, Peter Ache, Klaus F. X. Mayer, Dirk Becker, M. Rob G. Roelfsema, Dietmar Geiger, Jörg Schultz, Rainer Hedrich

## Abstract

The evolution of adjustable plant pores (stomata), enabling CO_2_ acquisition in cuticle wax-sealed tissues was one of the most significant events in the development of life on land. But how did the guard cell signalling pathways that regulate stomatal movements evolve? We investigate this through comparison of fern and angiosperm guard cell transcriptomes. We find that these divergent plant groups share expression of similar genes in guard cells including biosynthesis and signalling genes for the drought stress hormone abscisic acid (ABA). However, despite conserved expression in guard cells, S-type anion channels from the SLAC/SLAH family – known for ABA-mediated stomatal closure in angiosperms – are not activated by the same pathways in ferns, highlighting likely differences in functionality. Examination of other land plant channels revealed a complex evolutionary history, featuring multiple gains or losses of SLAC activation mechanisms, as these channels were recruited to a role in stomatal closure. Taken together, the guard cells of flowering and non-flowering plants share similar core features, but also show lineage-specific and ecological niche-related adaptations, likely underlying differences in behaviour.

## Introduction

Land plants evolved from a green algal ancestor, which conquered dry land approximately 500 million years ago [1]. A major plant innovation that helped plants to thrive on land is the adjustable stomatal pore, which enables CO_2_ acquisition in cuticle-covered tissues. The stomatal pore is usually flanked by two guard cells that regulate stomatal aperture and are found in all major land plant lineages except liverworts. In vascular plants (lycophytes, ferns, gymnosperms and angiosperms), mature stomata are able to close and reopen throughout the day in response to internal and environmental signals and have a major role in preventing excessive water loss. In mosses and hornworts, stomata appear important for drying spores, opening once only through irreversible guard cell collapse on hornwort sporangia [2], and developing mechanical restrictions that prevent closure with maturity on moss spore capsules [3]. Mosses in the genus *Sphagnum* have pseudostomata that lack a stomatal pore but possess two guard cells that collapse irreversibly and promote spore desiccation [4, 5].

In angiosperms, osmotic regulation of guard cell turgor via ion flux is central to the control of active stomatal movements [see 6]. Stomatal closure in response to dehydration stress is mediated by the hormone abscisic acid (ABA), synthesised in leaves and even guard cells themselves during drought stress [7, 8]. In diverse angiosperms, orthologs of the S-type anion efflux channel SLAC1 are activated by the serine/threonine protein kinase OPEN STOMATA 1 (OST1) in an ABA-dependent, calcium-independent manner [9-11]. In addition, SLAC1 and its homolog SLAH3 are sensitive to calcium signals via activation by calcium-dependent protein kinases (CPKs) and/or Calcineurin B-like protein (CBL)-interacting protein kinases (CIPKs) [12-15]. These calcium-sensitive kinases form a separate branch of the ABA-signalling pathway [13, 15-17], with ABA inducing calcium signals in guard cells prior to stomatal closure [18]. Other members of the SLAC/SLAH family – SLAH1, SLAH2 and SLAH4 – have apparent roles in translocation of Cl^-^ and NO ^-^ ions in roots and vascular tissues for nutrition and salinity tolerance [19-21].

ABA perception and signalling components appear largely conserved between land plants [22-25]. However, whether or not ABA has a conserved role in stomatal closure in seedless plant groups is currently debated [see 26, 27]. Notably, so far only seed plants have been shown capable of closing stomata in response to elevated endogenous ABA levels [28, 29]. Intriguingly, experiments in *Xenopus* oocytes have revealed that SLAC/SLAH channels from the alga *Klebsormidium nitens*, the liverwort *Marchantia polymorpha*, the lycophyte *Selaginella moellendorffii* and the fern *Ceratopteris richardii* (two CrSLAC homologs tested) lack the capacity for activation by OST1 kinases from the same species, but a functional OST1-SLAC pair was isolated from among the 4 SLAC/SLAH family members present in the moss *Physcomitrium/Physcomitrella patens* [30, 31]. This indicates that gains or losses of SLAC activation mechanisms appeared between divergence of mosses and their last common ancestor with flowering plants during land plant evolution.

To investigate the evolution of guard cell signalling pathways, we examined the transcriptomes of two fern species, *C. richardii* and *Polypodium vulgare*, in comparison to the angiosperms Arabidopsis and barley. We tested the activity of the guard cell-expressed fern SLAC channels and looked in depth at the evolution of SLAC activity in land plants using additional bryophyte and seed plant models. Our results indicate that fern guard cells express homologs of many important angiosperm guard cell genes, but that fern SLAC proteins are not activated by the same ABA-dependent pathways as angiosperms. We find that the evolution of SLAC activity in land plants has had a complex history featuring multiple gain or loss events. Our results reveal the diversity of guard cell regulatory mechanisms in plants alive today, while shedding light on how these have evolved over the past 500 million years of plant life on land.

## Results

### Fern and angiosperm guard cells express similar types of genes with some lineage-specific differences

For insight into how fern guard cell transcriptomes compare to those of angiosperms, we performed RNA-seq experiments using Arabidopsis, barley (*Hordeum vulgare*) and two members of the largest extant fern group – the Polypodiidae: *C. richardii* – an aquatic fern with a history as a model for genetics and molecular biology [32, 33], and *P. vulgare* – an historical stomatal research model shown capable of rapid and reversible stomatal responses to air humidity [34-36]. For each species, we enriched guard cells by mechanical isolation of epidermal fragments in a way that disrupts other cell types [7], and compared gene expression levels to whole leaf samples. For *P. vulgare*, we also compared whole leaf samples to leaf samples with abaxial epidermis removed, lacking guard cells, to confirm decreased expression of guard cell-expressed genes. We assembled sporophyte leaf transcriptomes for the two fern species and identified orthogroups (gene sets that are each descended from a single gene in a last common ancestor) containing characterised genes of interest including model species from major land plant lineages and streptophyte algae. For evolutionary reconstruction, we classified each orthogroup based on the presence and absence of genes significantly (p ≤ 0.01) enriched in guard cells into one of three groups: shared, angiosperm-specific and fern-specific. We defined ‘shared’ as groups with guard cell enriched genes in all four species, ‘angiosperm-specific’ as those with guard cell-enriched genes in both angiosperms but neither fern, and ‘fern-specific’ orthogroups as those in which the genes of both ferns but neither of the angiosperms were enriched. We found 172 shared, 214 angiosperm-specific and 275 fern-specific guard cell orthogroups, with the majority of these sharing a deep-rooted, ancient ancestor with green algae (Fig. 1A).

**Figure 1.**
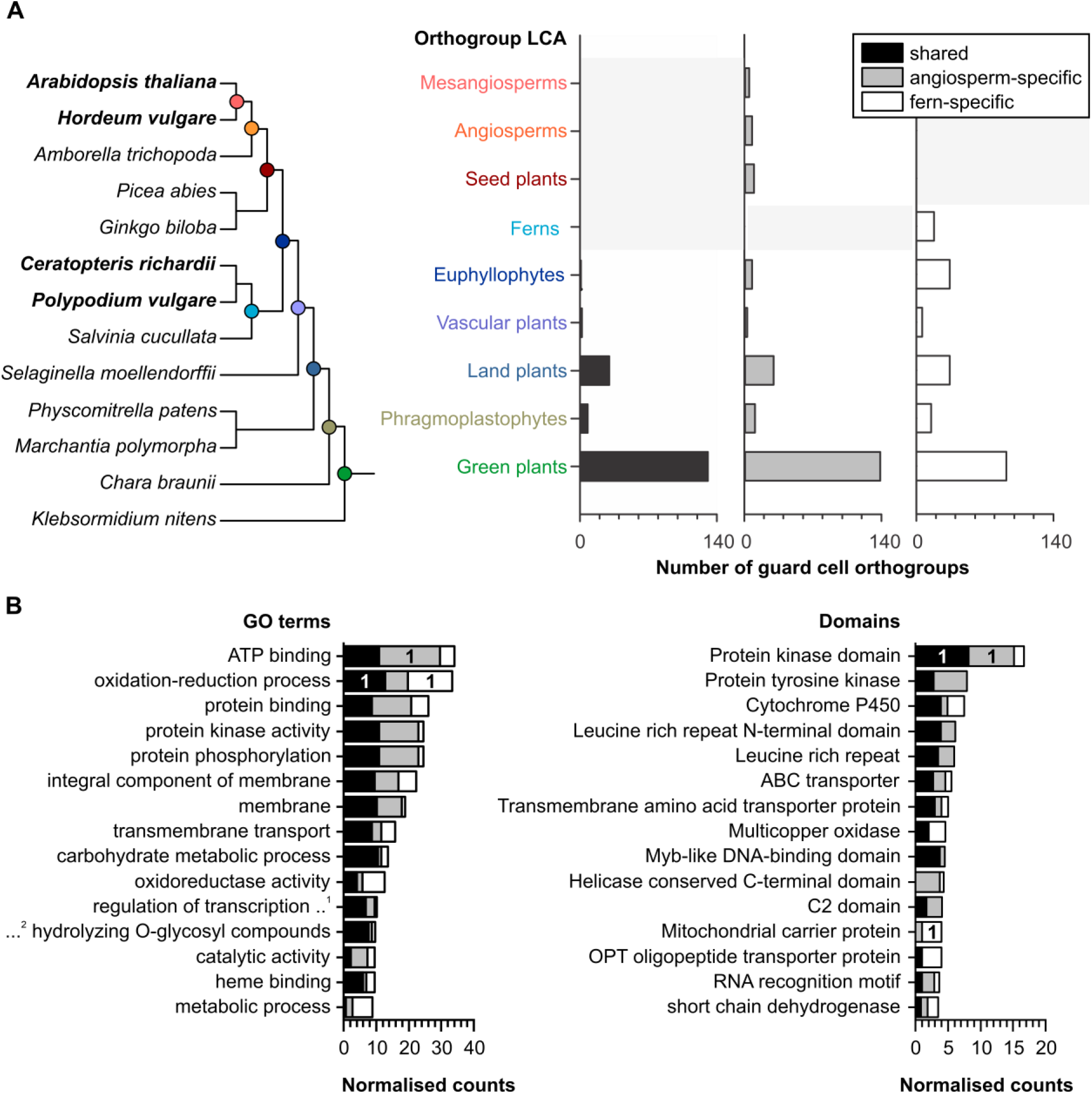
There are core sets of genes shared between angiosperm and fern guard cell transcriptomes with some lineage-specific differences. **(A)** Plant species included in RNA-seq experiments (bold) and orthology analyses with last common ancestor (LCA) nodes coloured (left) and orthogroups with significantly higher expression in guard cell than whole leaf samples (right) in all angiosperm (*A. thaliana, H. vulgare*) and fern (*C. richardii, P. vulgare*) species examined (shared; black), both angiosperm but neither fern species (angiosperm-specific; grey), and both fern but neither angiosperm (fern-specific) shown according to the LCA in each group. **(B)** The top 15 most common Gene Ontology (GO) terms and annotated domains of guard cell orthogroups, with counts normalised to the total number of genes per orthogroup. The most common GO term/domain for each expression group is indicated with “1”. Abbreviations are as follows …^1^ – “DNA templated”, …^2^ – “hydrolase activity”. See Table S1 for all GO terms/annotated domains related to guard cell orthogroups.

To gain insight into the functional commonalities and differences between these groups, we compared their domain content as well as their Gene Ontology (GO) classifications. Indeed, we identified common themes but also lineage-specific differences (Fig. 1B; Table S1). Overall, similar types of genes show preferential expression in guard cells of ferns or angiosperms specifically, particularly genes involved in metabolic processes, signalling and transport, suggesting that these core processes are critical in guard cells across vascular plants. Genes linked to oxidation-reduction processes were the most common among shared and fern-specific guard cell orthogroups, while ATP-binding was the most common GO classification in angiosperm-specific orthogroups, reflective of the importance of metabolic and energy-fuelled processes in guard cells. Genes encoding protein kinase domains were most common in shared and angiosperm-specific groups, with shared guard cell protein kinases including mitogen-activated protein kinases (MAPKs) related to AtMPK12 and AtMPK4, which have a role in stomatal CO_2_ and pathogen defence signalling in angiosperms [37, 38], and more general roles in immune signalling conserved in other land plants [39]. Angiosperm-specific guard cell protein kinases included cysteine-rich receptor-like kinases (CRKs), which have roles in biotic and abiotic stress signalling in Arabidopsis including stomatal closure [40]. Genes encoding mitochondrial carrier protein domains were the most common in fern-specific guard cell orthogroups, indicating the importance of mitochondrial transport in fern guard cells; this may reflect the unusually high number of chloroplasts present in fern guard cells [41, 42], which work together with mitochondria to supply cells with energy and metabolites.

### Fern guard cells express homologs of ABA biosynthesis and signalling genes

In angiosperms, key genes for guard cell regulation are well described. In contrast, less is known about the presence or absence of these genes in fern guard cells, hindering the reconstruction of guard cell evolution. Using our angiosperm and fern guard cell RNA-seq data, we looked specifically at orthologs of key angiosperm genes. Consistent with angiosperms, orthologs of the conserved guard cell specification gene *FAMA* [43, 44], showed significantly higher expression in fern guard cells than whole leaf samples (Fig. 2; Table S2). Similarly, both ferns showed expression of homologs of ABA biosynthesis and signalling genes in the guard cell samples, similar to the angiosperms (Figs. 2 and S1; Table S2). Although orthologs of ABA2 – a short chain dehydrogenase (SDR) dedicated to the ABA biosynthesis pathway [45, 46] – are restricted to angiosperms [47-49], other SDRs are likely able to catalyse this step, and some closely-related SDRs were expressed in the fern guard cell samples (Fig. S1; Table S2). Among the ABA-signalling pathway, some genes encoding PP2CA phosphatases and kinases from the CPK and CIPK families were expressed at higher levels in guard cell than whole leaf samples in the ferns (Fig. 2). Other ABA-signalling genes including members of the *OST1* subclade of the *sucrose non-fermenting-1-related protein kinase 2* (*SnRK2*) family were also non-specifically expressed in guard cells of the ferns. Among the ion channels that are downstream of ABA-signalling genes in angiosperms, S-type anion channel (SLAC/SLAH) genes were enriched in guard cells relative to whole leaves in all species. Overall, these results suggest that, similar to Arabidopsis and barley [7, 11], fern guard cells are equipped with the genetic toolkit required for ABA biosynthesis and signalling.

**Figure 2.**
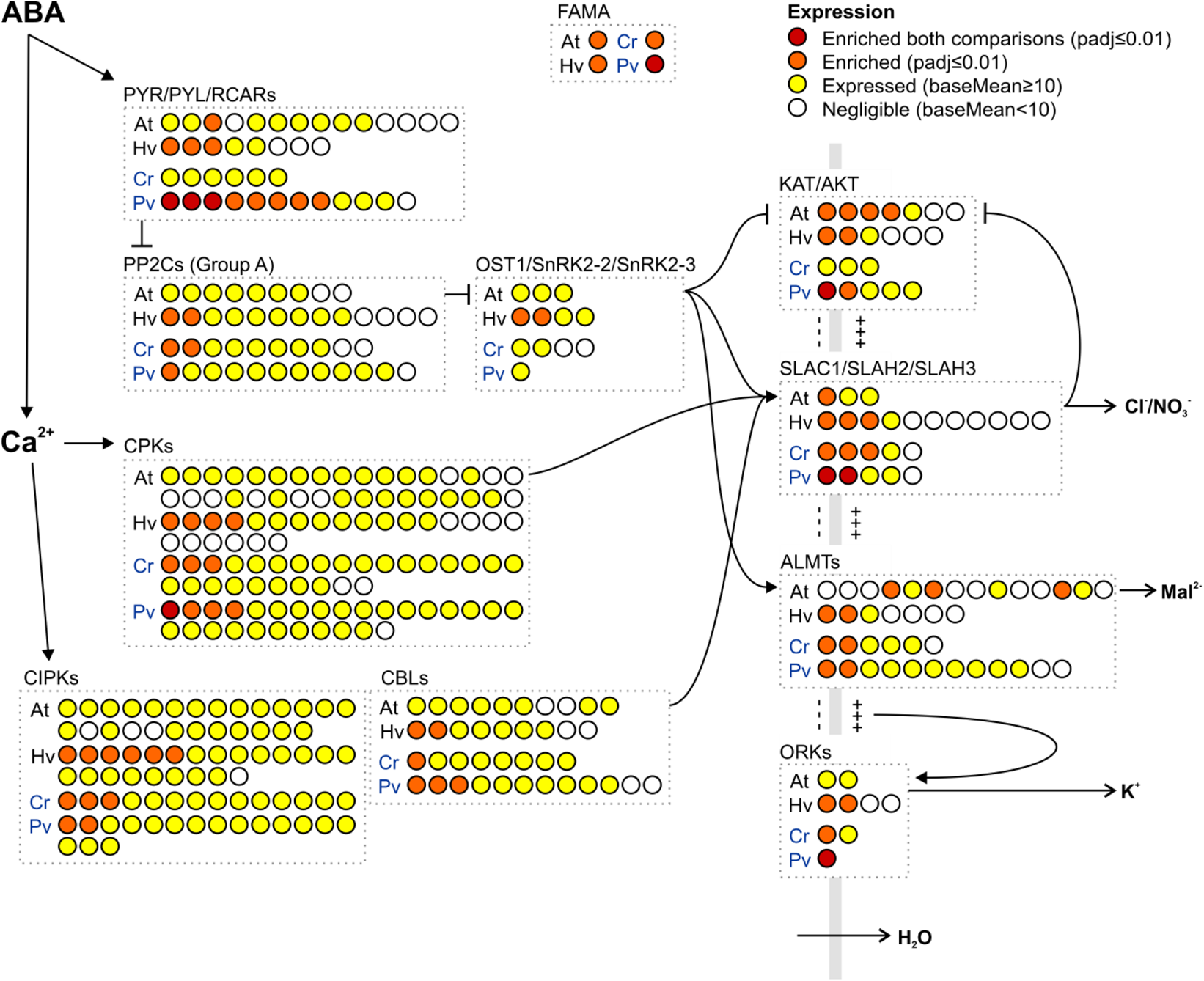
Homologs of key ABA-signalling genes and ion channels are expressed in angiosperm and fern guard cells. Groups of related genes are shown together in boxes with each circle representing a different gene and colour coding representing relative expression in guard cell-enriched samples compared to whole leaves is shown for angiosperm (At, *Arabidopsis thaliana*; Hv, *Hordeum vulgare* barley) and fern (Cr, *Ceratopteris richardii*; Pv, *Polypodium vulgare*) models, as indicated. For *P. vulgare* samples only, whole leaf vs leaf samples without abaxial epidermis (thus guard cells) removed were also included and used to separate guard cell-enriched genes with a higher level of stringency (red; “enriched both comparisons” = expression higher in guard cells than leaves and higher in whole leaves than leaves without guard cells). Orthologs of the guard cell specification gene *FAMA* are also shown. The vertical grey line represents the guard cell plasma membrane with guard cell cytoplasm on the left and apoplast on the right. Please note that for simplicity the orthogroups are displayed according to the attributes of members with key roles in guard cell signalling, but do not reflect the characteristics of all orthogroup members. See Figure S1 for ABA biosynthesis genes and Table S2 for details for all genes of interest examined.

### Fern SLAC channels are not activated by ABA-signalling kinases

To examine the stomatal responses of the fern *P. vulgare* to ABA, relative to those of the angiosperm tobacco (*Nicotiana tabacum*), we adapted a recently-developed current-ejection method [18] to apply ABA directly to guard cells in intact fern plants and monitor stomatal movements with an upright fluorescence microscope. Microelectrodes, filled with 1 mM ABA and/or 1 mM Lucifer Yellow (LY), were put in contact with the apoplast, close to an open stoma. Application of a -1 nA current for 1 min caused the appearance of fluorescence in guard cell walls of *P. vulgare* and tobacco (Fig. 3A), showing that negatively charged molecules can be applied to guard cells with this approach. The current-ejection of LY into the guard cell walls did not provoke stomatal closure, while ABA provoked rapid stomatal closure in tobacco stomata (Fig. 3B-C). To confirm that stomata prechallenged with the control LY solution had not lost ABA responsiveness, we applied the stress hormone in a second ejection. ABA-induced stomatal closure was similar between guard cells pretreated with LY and those that were not (Fig. 3C). Apoplastic application of ABA with microelectrodes thus causes rapid closure of tobacco stomata, just as previously reported for Arabidopsis [18]. In contrast, no response to ABA could be observed for *P. vulgare* (Fig. 3B-C). These results indicate that, unlike angiosperms, the fern *P. vulgare* does not close its stomata in response to ABA. These findings are consistent with those of others who have found a lack of stomatal sensitivity to ABA in ferns [28, 29].

**Figure 3.**
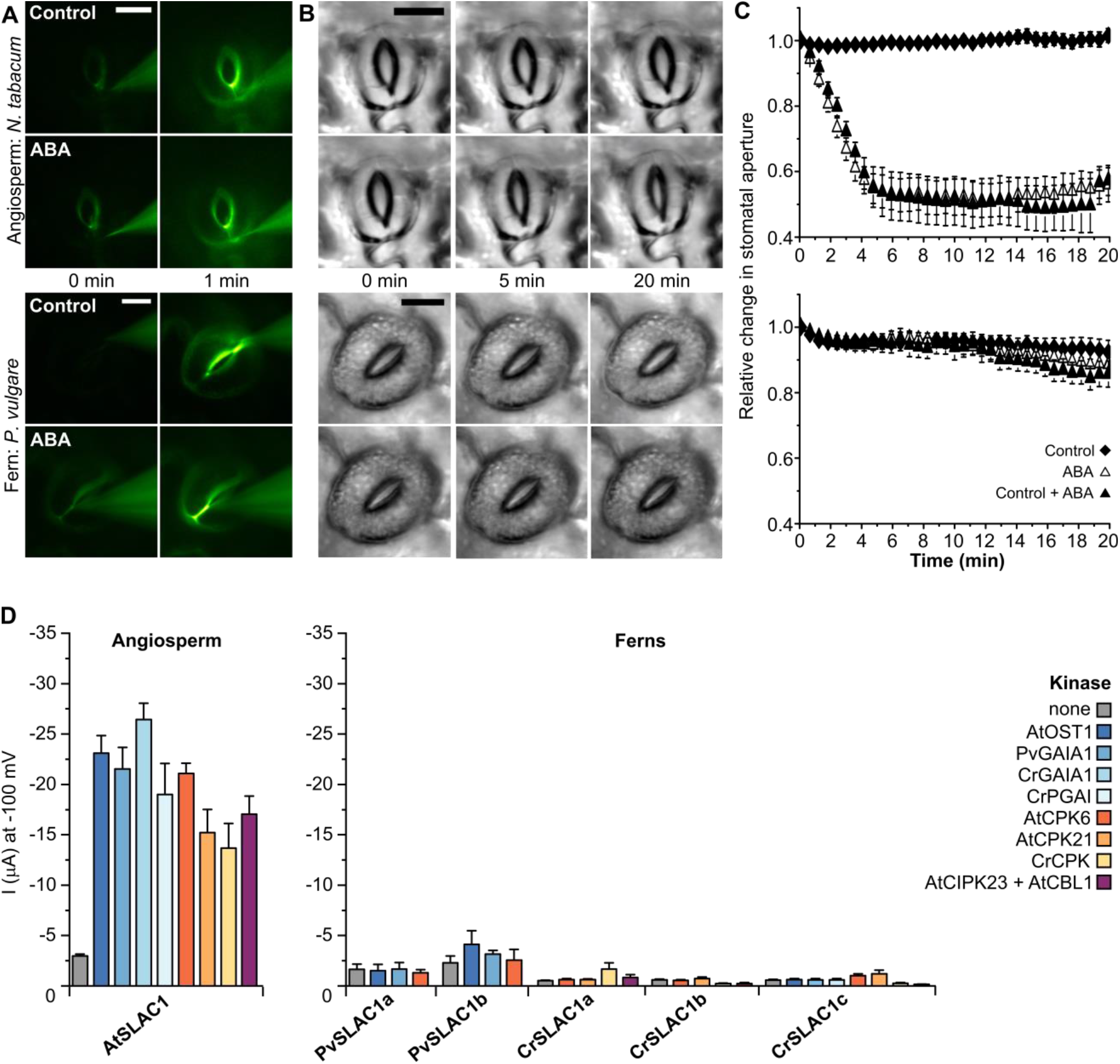
Unlike angiosperm stomata, fern stomata are insensitive to ABA, and fern guard cell SLAC homologs are not activated by ABA-signalling kinases. **(A)** Fluorescent image of guard cell walls of tobacco (upper panels) and *P. vulgare* (lower panels) at the start and 1 min after current-ejection of 1 mM Lucifer Yellow (LY; control), or 1 mM LY and 1 mM ABA (scale bar = 10 µm). **(B)** Brightfield images of the guard cells shown in **(A)** up to 20 min after current-ejection. **(C)** Normalised stomatal aperture, plotted against time for tobacco (upper graph) and *P. vulgare* (lower graph). Current-ejection was performed at t = 0. Stomata treated with LY (control, diamonds) and subsequently to ABA (control + ABA, closed triangles), or to ABA only (ABA, open triangles; mean ± SEM, n ≥ 7). **(D)** Mean whole-oocyte current measurements at −100 mV in nitrate-based standard medium with SLACs from the angiosperm *A. thaliana* and the ferns *P. vulgare* and *C. richardii*, co-expressed with or without the indicated kinases in *Xenopus* oocytes (mean ± SEM, n ≥ 4).

In angiosperms, orthologs of the anion channels SLAC1 and SLAH3 play a key role in stomatal closure responses to ABA, after activation by OST1-, CPK-or CIPK-type kinases [see 50]. We previously found that two *C. richardii* SLAC/SLAH family members, CrSLAC1a and CrSLAC1b, were not activated by Arabidopsis kinase AtOST1 or the fern OST1-subclade kinases GAMETOPHYTES ABA INSENSITIVE ON A_CE_1 (CrGAIA1) and PARALOG OF GAIA1 (CrPGAI) in the *Xenopus* oocyte expression system [31]. In this study, we identified four additional SLAC/SLAH family members in *C. richardii* between our sporophyte transcriptome (*CrSLAC1c-e*; Fig. 2) and an additional *C. richardii* gametophyte transcriptome (*CrSLAC1f*; [51]). *CrSLAC1a*-*c* are expressed at higher levels in guard cells than whole leaf samples, as are two SLAC/SLAH family members in *P. vulgare* (*PvSLAC1a* and *PvSLAC1b*; Figs. 2, S2 and S3).

Using the *Xenopus* oocyte expression system together with the double-electrode voltage-clamp technique, we tested if any of the newly-isolated *C. richardii* and *P. vulgare* SLAC homologs are activated in response to kinases that activate AtSLAC1. Specifically, we tested Arabidopsis ABA-signalling kinases, all guard cell-expressed kinases from the SnRK2 OST1-subclade in the two fern species, and a guard cell-enriched CPK from *C. richardii* (Figs. 2 and S4). Each of these kinases resulted in strong anion channel currents when co-expressed with AtSLAC1, but did not trigger increased currents with any of the *P. vulgare* guard cell-enriched SLAC homologs, or any of the *C. richardii* SLAC homologs expressed in currently-available transcriptomes (Figs. 3D and S2). The fern SLAC homologs all contain differences in the C-terminal domain, which includes AtSLAC1 T513, and was previously found to be important for PpSLAC1 activation [30]. We created a mutant in which the C-terminal domain was changed to be the same as AtSLAC1 (PvSLAC1a V663L), but this was not sufficient to impart kinase-induced SLAC activity (Fig. S2). Overall, these results indicate that the fern SLAC homologs are not activated by the same kinases as angiosperm SLAC1 and SLAH3 orthologs, consistent with the lack of stomatal response to ABA we measured in *P. vulgare*.

To confirm anion channel activity, we constructed CrSLAC1a F592A channel mutants for the SLAC1 gate residue corresponding to AtSLAC1 F450A [52]. Consistent with previous measurements for AtSLAC1 F450A [53], we measured strong currents for the CrSLAC1a F592A mutant when co-expressed with AtOST1, but only very small background currents with CrSLAC1a F592A expressed alone (Fig. S2). This confirms that CrSLAC1a encodes an anion channel that can be activated in the absence of gating restrictions.

### Moss and hornwort SLAC homologs can be activated by OST1

Given the previous finding of an active SLAC1-OST1 pair from the moss *P. patens* (PpSLAC1-PpOST1.2) [30], we further examined the evolution of SLAC activity using the distantly-related moss *Sphagnum fallax* and the hornwort *Anthoceros agrestis* to determine if other bryophytes share kinase-activated SLAC homologs. There has long been doubt over the phylogenetic relationship between hornworts and other land plants, but studies using nuclear and chloroplast data have supported a monophyletic relationship between bryophytes [54, 55]. We searched available genome sequences and identified the presence of a single SLAC/SLAH family member in *A. agrestis*, and two orthologs of *PpSLAC1* and three members of the OST1 subclade of SnRK2s in *S. fallax* (Table S3; Figs. S3 and S4), and were able to isolate all genes from gametophyte tissues. We found that both *S. fallax* and *A. agrestis* possess a SLAC that can be activated by ABA-signalling kinases in the *Xenopus* system (Fig. 4), similar to *P. patens* [30]. This indicates that among bryophytes, moss and hornwort species possess kinase-activated SLAC homologs.

**Figure 4.**
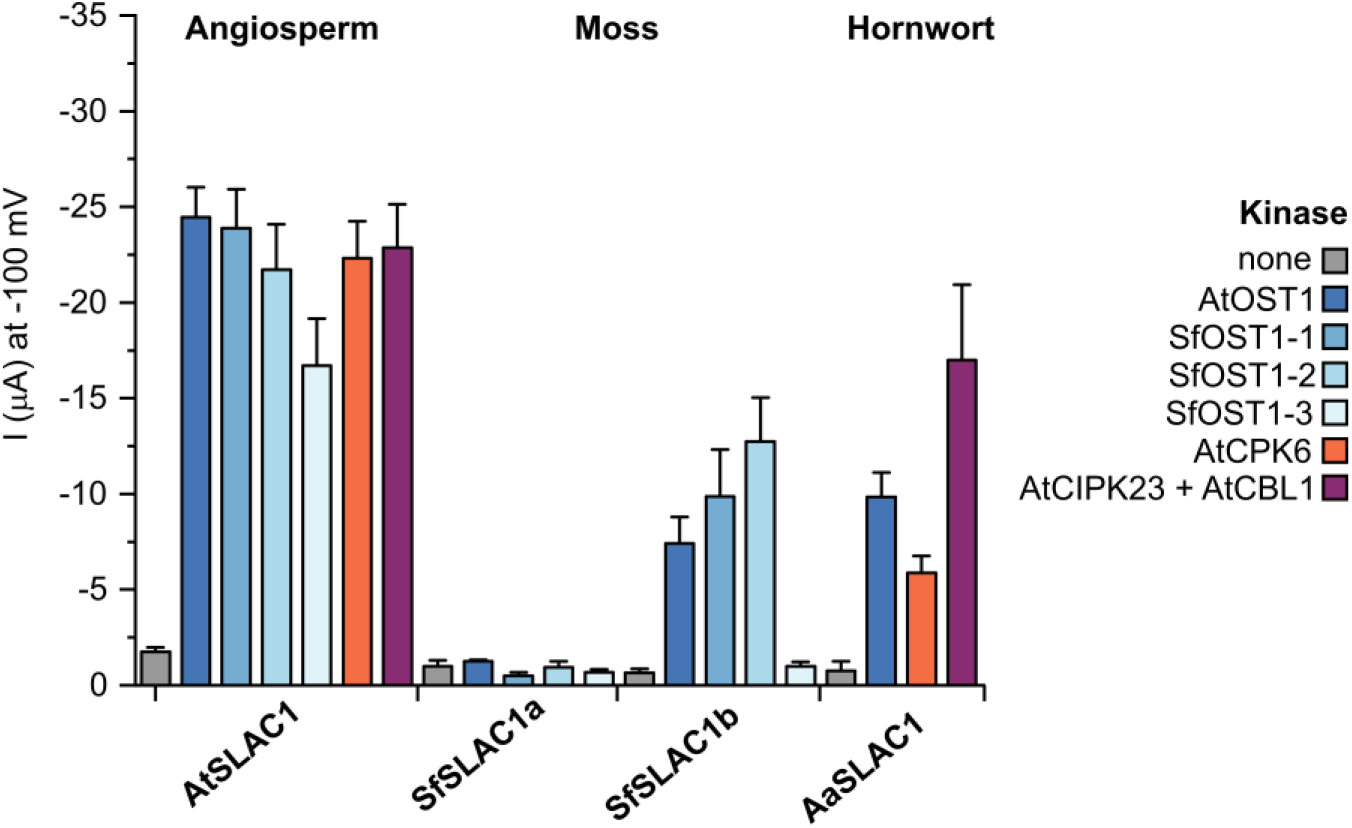
Moss and hornwort SLAC homologs can be activated by ABA-signalling kinases. Mean whole-oocyte current measurements at −100 mV in nitrate-based standard medium with SLAC homologs from the angiosperm *A. thaliana*, the moss *S. fallax* and the hornwort *A. agrestis*, co-expressed with or without the indicated kinases in *Xenopus* oocytes (mean ± SEM, n ≥ 4).

### Activation in gymnosperm and ‘basal angiosperm’ lineages requires auxiliary factors

Seed plants (gymnosperms and angiosperms) are the only plants to possess clear orthologs of Arabidopsis SLAC1, SLAH2/3, and SLAH1/4 that are strongly supported by phylogenetic analyses (Fig. S3; [56]). So far, the only seed plant SLAC1 orthologs to be examined have been from the core angiosperm lineages – dicots and monocots; SLAC1 activation has not previously been studied in gymnosperms and ‘basal angiosperm’ lineages. To further examine the evolution of SLAC1 activity in seed plants we made use of the model gymnosperm *Picea abies*, which shows stomatal sensitivity to ABA, similar to angiosperms [57]. The *P. abies* genome encodes four SLAC/SLAH proteins – two orthologs of AtSLAC1 (PaSLAC1a, PaSLAC1b), an ortholog of both AtSLAH1 and AtSLAH4 (PaSLAH1), and an ortholog of both AtSLAH2 and AtSLAH3 (PaSLAH2) (Fig. S3; Table S3; [6]). Investigation of the protein sequences revealed the absence of serine or threonine residues in PaSLAC1a in the region corresponding to AtSLAC1 S120, and in PaSLAC1b in the region of AtSLAC1 S59 (Fig. 5B) – both residues are required (although not sufficient) for phosphorylation of SLAC1 proteins by OST1 and CPKs [9, 16, 58]. When co-expressed with OST1 in *Xenopus* oocytes, the *P. abies* SLAC1 orthologs did not show S-type anion channel currents (Figs. 5A and S5). Further examination of other gymnosperm SLAC1 ortholog sequences revealed that those of other species from the Pinaceae family similarly lack S120 (Fig. 5B). However, a serine is present at these positions in the SLAC1 orthologs of species from other gymnosperm families, including *Ginkgo biloba*. We tested the activity of this *Ginkgo biloba* ortholog (GbSLAC1a) and found that it could be activated by OST1 and CPKs (Fig. 5A and C). Replacement of the S120 region in PaSLAC1a with the six amino acids from GbSLAC1a that differ in this region (Fig. 5B), failed to impart sensitivity to ABA-signalling kinases (Fig. S5). However, introduction of a mutation to the SLAC1 gate residue PaSLAC1a F512A enabled substantial currents when co-expressed with a kinase (OST1 or CPK), indicating that PaSLAC1a can carry anion currents in nitrate-based solutions when gated open (Fig. S5). In contrast to the *P. abies* SLAC1 orthologs, wild type PaSLAH2 (an ortholog of both AtSLAH2 and AtSLAH3) yielded substantial currents in nitrate-based media in the presence of a CPK (Fig. 5A), similar to AtSLAH3 [12]. This indicates that, in addition to OST1s or CPKs, auxiliary factors are needed for kinase activation in some lineages.

**Figure 5.**
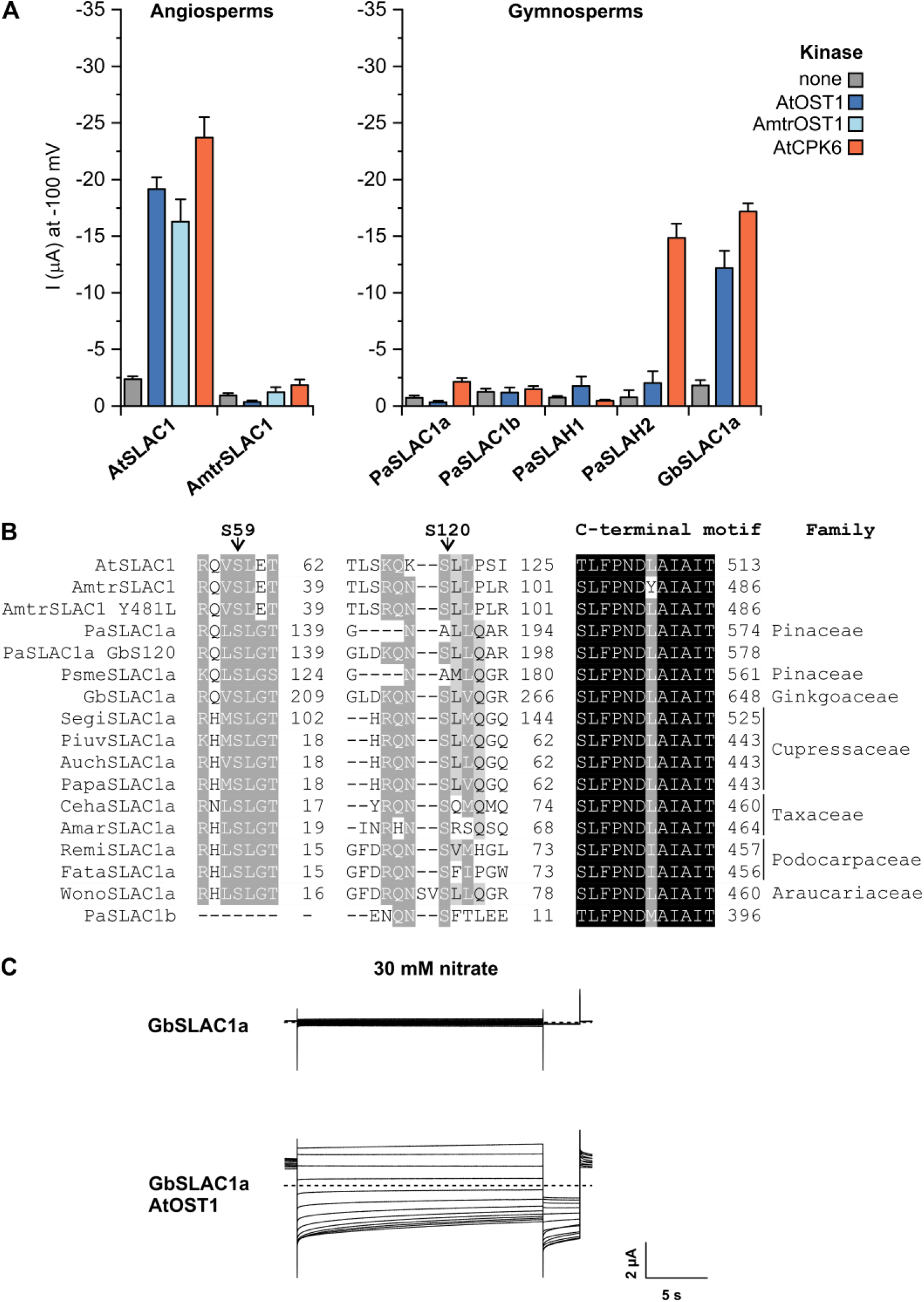
Seed plants differ in terms of SLAC1 activity, but SLAH2/3 ortholog activity may be sufficient for sensitivity to ABA-signalling kinases. **(A)** Mean whole-oocyte current measurements at −100 mV in nitrate-based standard medium with SLAC/SLAH channels from the angiosperms *A. thaliana* and *A. trichopoda*, and the gymnosperms *P. abies* and the *G. biloba*, co-expressed with or without kinases in *Xenopus* oocytes (mean ± SEM, n ≥ 4). **(B)** Alignment of protein regions surrounding AtSLAC1 residues S59, S120 and the C-terminal motif important for activation by ABA-signalling kinases, showing S59 is not conserved in PaSLAC1b, S120 is conserved in seed plants except members of the gymnosperm Pinaceae family, and the C-terminal motif is not conserved in AmtrSLAC1. Shading levels indicate degree of conservation (black = 100%, dark grey = 80%, and light grey = 60%). Numbers indicate position in each protein sequence. Family is indicated for gymnosperm species, and sequences for mutants Y481L and GbS120 are indicated (see Fig. S5). Full sequence details are given in Table S3. **(C)** Example whole-oocyte currents of GbSLAC1a, either expressed alone (upper panel) or with AtOST1 (lower panel), recorded in nitrate-based bath solution. Voltage pulses lasting 20 s ranging from +40 to −180 mV in 20 mV decrements were applied (holding potential V_H_ was 0 mV).

To further examine the evolution of SLAC1 activity in seed plants we made use of the single living representative of the Amborellales – the sister lineage to all other extant flowering plants – *Amborella trichopoda* [59]. The *A. trichopoda* genome encodes a single SLAC1 ortholog (AmtrSLAC1), two orthologs of SLAH2/3 and a single ortholog of AtOST1/AtSnRK2.2/AtSnRK2.3 (AmtrOST1; Figs. S3 and S4; Table S3; [6, 27]). We examined the activity of AmtrSLAC1 when co-expressed with AtOST1, AmtrOST1 or AtCPK6, and found that it was insensitive to activation from any of these ABA-signalling kinases (Fig. 5A). As AmtrSLAC1 contains a difference in its C-terminal motif relative to AtSLAC1, we created a mutant where this motif is changed to resemble AtSLAC1 (AmtrSLAC1 Y481L), however this was not sufficient to impart sensitivity to OST1 kinases (Fig. S5).

## Discussion

We find that guard cells of the fern models *C. richardii* and *P. vulgare* are equipped with ABA biosynthesis and signalling pathways (Figs. 2 and S1). Previous studies have suggested that expression of ABA-signalling and ion channel homologs in stomata-bearing tissues indicates a potential conserved role in ABA-mediated stomata closure [60, 61]. However, we found *P. vulgare* to be insensitive to ABA, and guard cell SLAC homologs from the ferns *P. vulgare* and *C. richardii* to be insensitive to ABA-signalling kinases (Fig. 3). These results are consistent with previous findings of a lack of stomatal closure response to endogenous ABA levels in ferns [28, 29], and the lack of stomatal phenotype for ABA-insensitive mutants of the *C. richardii OST1* ortholog *GAIA1* [31] in sharp contrast to the wilty phenotypes of angiosperm ABA biosynthesis and signalling mutants [49, 62-64]. In addition, a recent study showed a lack of ABA-mediated K^+^ efflux from the guard cells of fern and lycophyte species, further emphasising differences between the stomatal responses of seed plants and other vascular plant groups [65]. Together, these results support a passive model for stomatal closure in ferns [29], whereby guard cell turgor is dependent on leaf turgor rather than ABA-dependent activation of guard cell-expressed ion channels.

Ferns synthesise ABA in response to dehydration stress, similar to all other land plant groups [23, 28, 66-69], consistent with an ancient role for ABA in desiccation tolerance [25, 70-73]. In response to slow dehydration, ABA triggers the upregulation of proteins that function in osmoregulation to protect fragile cellular components during desiccation, including metabolic enzymes, sugar transporters and aquaporins, in diverse plant species including bryophytes, lycophytes, ferns and angiosperms [25, 69, 72, 74-77]. In line with this function, ABA does induce changes in gene expression in ferns including in guard cells [61]. Thus, although our results indicate that ABA signalling is not involved in rapid stomatal closure via ion channel activation as occurs in seed plants, it is likely that ABA signalling is involved in slow dehydration tolerance mechanisms in ferns, similar to other land plants.

We do find kinase-sensitive SLAC homologs in the mosses *P. patens* and *S. fallax* and the hornwort *A. agrestis* (Figs. 3-5; [30]), where they might serve plant anion transport functions. However, it is not physically possible for SLAC homologs to play any role in stomatal closure in *S. fallax* and *A. agrestis*, as stomata ‘open’ by irreversible guard cell collapse in these species [2, 4, 5]. In other mosses, developmental thickening of the guard cell walls prevents stomatal movement at maturity [3]. These features are in line with a predominant role for moss and hornwort stomata in promoting desiccation for spore drying and dispersal [2, 43, 78]. Although a previous study identified small differences in stomatal pore aperture in response to applied ABA in *P. patens ost1-1* mutants [79], PpOST1-1 kinase did not activate PpSLAC1 or PpSLAC2 anion currents, when tested [30], and the mechanism for any role in stomatal movement has not yet been shown. In accordance with an ancient role for ABA in desiccation tolerance [see 80], quadruple knockout mutants for all SnRK2 genes in *P. patens* – including the *P. patens* SnRK2 shown capable of inducing PpSLAC1 activity are unable to survive freezing and sustained dehydration of protonemata – the earliest developmental stage of the moss life cycle [25]. Gametophyte tissues (including protenemata) lack stomata, but do show strong expression of SLAC and OST1 homologs in *P. patens* [see 6, 27]. It is tempting to speculate that SLAC homologs may play a role in nutrient movement in non-flowering plants, similar to that seen for SLAH channels in angiosperms [19-21], and that this ancient role for SLAC homologs may have been co-opted for ABA-dependent ion movement associated with desiccation tolerance in gametophyte tissues in mosses and hornworts. Other potential roles for SLAC homologs in osmoregulation include saving thin-walled cells (such as tip growing cells) from bursting during rainfall. Future mutant studies are required to test this and determine the precise roles of SLAC homologs in bryophytes.

We have found that SLAC homologs from the alga *K. nitens*, liverwort *M. polymorpha* and lycophyte *S. moellendorffii*, in addition to the ferns *C. richardii* and *P. vulgare* and SLAC1 orthologs of seed plants *P. abies* and *A. trichopoda* lack sensitivity to ABA-signalling kinases (Figs. 3-5; [30, 31]). The lack of activation of the seed plant SLAC1 orthologs was particularly unexpected, given the shared ABA-mediated stomatal closure response in these species [57, 81], but it is possible that activation of SLAH2/3 orthologs by ABA-signalling kinases is sufficient for this response in these species. Other ABA-independent mechanisms for activation of SLAC homologs including acetate and other SLAC homologs – which occur in angiosperms [19-21] – remain to be tested in other plant groups.

Overall, we find evidence of a complex evolutionary history for SLAC sensitivity to ABA-signalling kinases with either i) multiple gains or ii) an early gain for a non-guard cell-specific functional origin and subsequent losses (Fig. 6). Given the absence of this trait in lycophytes and ferns, we propose that bryophytes and seed plants may have separately co-opted SLAC/SLAH channels for different roles downstream of the ABA-signalling pathway, with seed plants using ABA to trigger rapid stomatal closure and bryophytes using ABA for osmoregulation more generally in other cell types including vegetative tissues that lack stomata. We suggest the possibility that in some S-type anion channels, distant site mutations or factor binding may cause SLAC structural rearrangements that expose kinase-sensitive sites. Our results explain differences in ABA sensitivity between ferns and seed plants and highlight the importance of functional studies in addition to genomic and transcriptomic studies for understanding the evolution of plant signalling processes.

**Figure 6.**
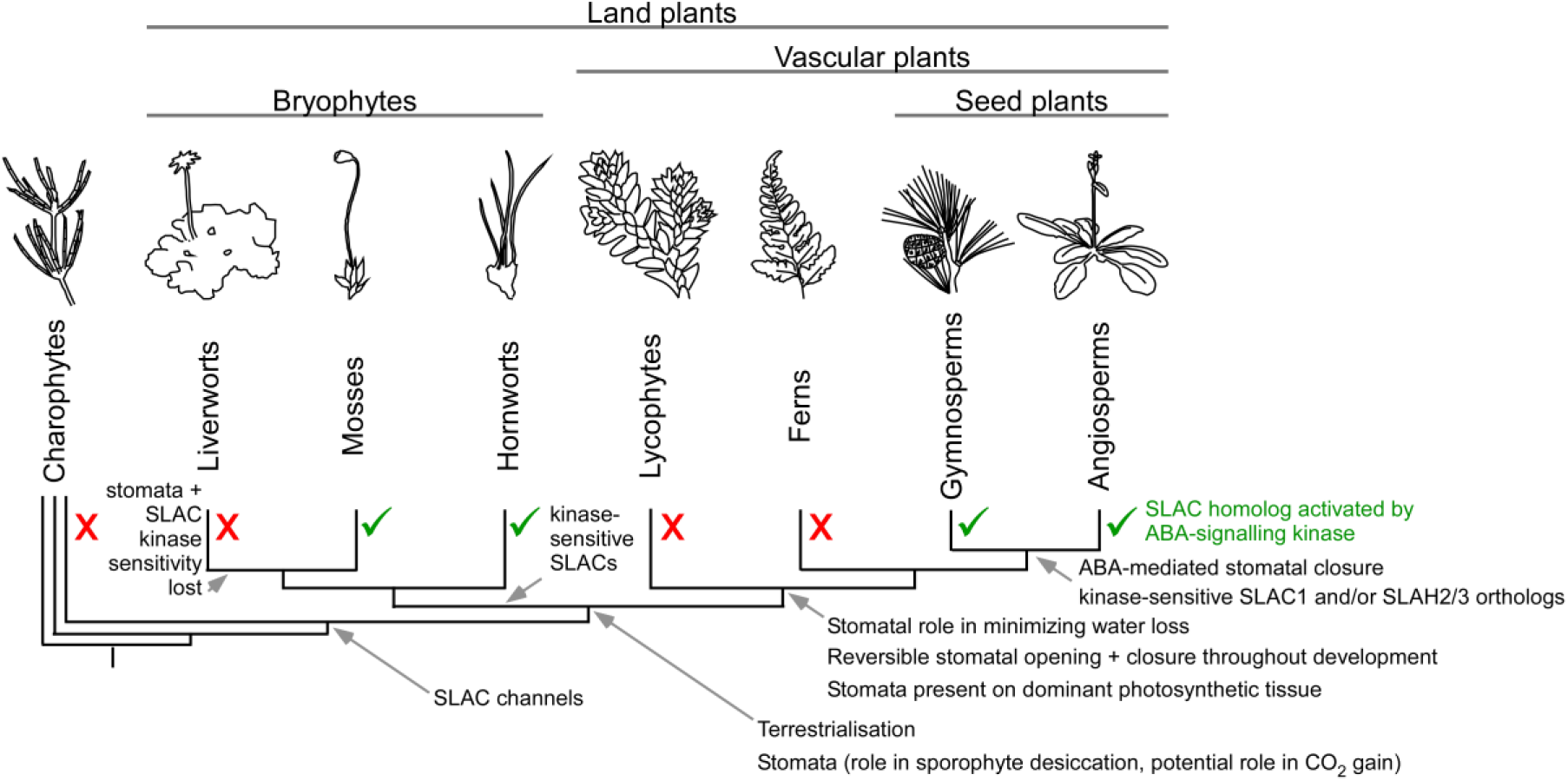
A model for the timing of key events during stomatal evolution. The hypothesised timing of key traits is indicated on the current model of land plant phylogeny [54, 55]. Branch lengths are not to scale.

## Material and Methods

### ABA electro-infusion

*Polypodium vulgare* plants were obtained from the Würzburg Botanical Gardens grown in a greenhouse with *Nicotiana tabacum* L. cv SR1, under natural light extended with additional light supplied with HQL-pressure lamps (Philips, http://www.lighting.philips.com, Powerstar HQI-E, 400 W) at a day/night cycle of 12/12 h. Extracellular application experiments (current ejection) with microelectrodes were conducted with intact plants. The adaxial side of fronds/leaves was attached with double-sided adhesive tape to a Plexiglas holder in the focal plane of an upright microscope (Axioskop 2FS, Zeiss, http://www.zeiss.com). Stomata were visualized with a water immersion objective (W Plan-Apochromat, 40x/0.8, or 63x/1.0, Zeiss) dipped into a drop of bath solution (5 mM KCl, 0.1 mM CaCl_2_, and 5 mM K-citrate, pH 5.0) placed on the abaxial surface of the frond or leaf. Microelectrodes were put in contact with the cell wall of stomata with a piezo-driven micro-manipulator (MM3A, Kleindiek Nanotechnik, http://www.nanotechnik.com), as previously described [18]. The microelectrodes were prepared from borosilicate glass capillaries (inner diameter, 0.58 mm; outer diameter, 1.0 mm; Hilgenberg, http://www.hilgenberg-gmbh.com) and filled with a solution containing 1 mM Lucifer Yellow (LY) and 1 mm ABA, or only 1 mM LY (control). The stomatal movements and ejection of LY were monitored with a charge-multiplying CCD camera (QuantEM, Photometrics, http://www.photometrics.com/). The fluorescent probe was excited with light from a Hg metal-halide lamp (HXP120, Leistungselektronic JENA, http://www.lej.de), which was filtered through a band pass filter of 430/24 nm (ET 430/24, Chroma Technology Corp., https://www.chroma.com). A dichroic mirror (495 nm LP) guided the excitation light through the objective, while the fluorescent light was filtered with an emission band pass filter (520/30 nm, BrightLine, Semrock). The filters could be rapidly moved in and out of the light-path, with filter wheels of a spinning disc confocal unit (CARV II, Crest Optics, http://www.crestopt.com) that was mounted to the camera port of the upright microscope. Image acquisition was conducted with VisiView software (Visitron, Puchheim, Germany) and analyzed the Image-J/Fiji software package [82].

### Arabidopsis RNAseq

Arabidopsis plants were grown in semi-sterilized soil (for 20 min at 100 °C), cultivated in climate chambers (Binder KBWF 720; www.binder-world.com) in a 12-h day-night rhythm (22/16 °C, 60% RH) and were illuminated with 125 μmol m^−2^ s^−1^ white light. RNA was isolated from whole leaf samples and guard cell-enriched samples isolated by mechanical disruption as previously described [7], from 6-7 week old plants. Library preparation and RNAseq were carried out as described in the Illumina TruSeq Stranded mRNA Sample Preparation Guide, the Illumina NextSeq 500 System Guide (Illumina, Inc., San Diego, CA, USA), and the KAPA Library Quantification Kit -Illumina/ABI Prism User Guide (Kapa Biosystems, Inc., Woburn, MA, USA).

Briefly, 250 ng of total RNA was used for purifying the poly-A containing mRNA molecules using poly-T oligo-attached magnetic beads. Following purification, the mRNA was fragmented to an average insert size of 200-400 bases using divalent cations under elevated temperature (94°C for 4 minutes). Next, the cleaved RNA fragments were reverse transcribed into first strand cDNA using reverse transcriptase and random hexamer primers. Actinomycin D was added to improve strand specificity by preventing spurious DNA-dependent synthesis. Blunt-ended second strand cDNA was synthesized using DNA Polymerase I, RNase H and dUTP nucleotides. The incorporation of dUTP, in place of dTTP, quenched the second strand during the later PCR amplification, because the polymerase does not incorporate past this nucleotide. The resulting cDNA fragments were adenylated at the 3’ ends, the indexing adapters were ligated and subsequently specific cDNA libraries were created by PCR enrichment. The libraries were quantified using the KAPA SYBR FAST ABI Prism Library Quantification Kit. Equimolar amounts of each library were sequenced on a NextSeq 500 instrument using two 150 Cycles High Output and one 150 Cycles Mid Output Kits with the single index, paired-end (PE) run parameters. Image analysis and base calling resulted in .bcl files, which were converted into .fastq files with the bcl2fastq v2.18 software. Library preparation and RNAseq were performed at the service facility “KFB -Center of Excellence for Fluorescent Bioanalytics” (Regensburg, Germany; www.kfb-regensburg.de). All sequences are accessible under the GenBank project accession PRJNA731641.

### Barley RNA-seq

Barley (Hordeum vulgare cv. Barke) seeds were provided by a commercial supplier (Saatzucht J. Breun GmbH & Co. KG) and cultivated at 22/16°C and 50 ± 5% RH at a 12/12h day/night cycle and a photon flux density of 500 mmol m^-2^ s^-1^ white light (Philips Master T Green Powers, 400 W). For guard cell-enriched samples, epidermal peels were first isolated from the abaxial side of 8 to 12-day-old leaves and then subsidiary cells were disrupted with successive blender cycles in ice-cold water as previously described [11]. RNA was extracted using the NucleoSpin^®^ RNA Plant Kit (Macherey-Nagel, Drueren, Germany). RNA isolation from whole leaves was performed similarly.

The extracted RNA was treated with RNase-free DNase (New England Biolabs, Ipswich, MA, USA). Quality control measurements were performed on a 2100 Bioanalyzer (Agilent, Santa Clara, CA, USA) and the concentration was determined using a Nanodrop ND-1000 spectrophotometer (Thermo Fisher Scientific, Wilmington, DE, USA). Libraries were prepared with the TruSeq RNA Sample Prep Kit v2 (Illumina, San Diego, CA, USA) using 1 mg of RNA and sequenced on a HiSeq 3000 (Illumina) resulting in a sequence depth of 35 million paired-end reads (2x 150bp). Sequences are accessible at EMBL-EBI ArrayExpress under E-MTAB-10534.

### Fern RNA-seq

*C. richardii* wild-type strain Hn-n [33] was grown in controlled-condition glasshouse facilities under a 16h photoperiod with supplemented natural light. *P. vulgare* plants (Common Polypody-winter hardy; Westdijk’s, Kwekerijen) were grown in a growth chamber under a 12h photoperiod, with 22°C day/16°C night temperatures.

Four biological replicates, each comprising up to 100 mg tissues were collected from both species for whole leaf samples comprising fully expanded sporophyte fronds lacking sporangia, and guard cell-enriched samples. Guard cell-enriched samples were obtained from fully-expanded fronds lacking sporangia, after removal of the main veins, by mechanical disruption of other cell types using a successive 1-2 min blending cycles in deionised ice water with epidermal fragments collected by filtration through a 210 µm nylon mesh, according to a previously-published method [7] optimised for *C. richardii* (3 cycles) and *P. vulgare* (5 cycles). Fluorescein diacetate staining [83] was used to confirm guard cell viability and purity. In *P. vulgare*, ‘leaf without epidermis’ samples lacking guard cells were also obtained by using fine forceps to fully remove the abaxial epidermis from leaves.

Total RNA was extracted, treated with RNase-free DNase and quality control measurements were performed as described above. Library preparation and mRNA-seq after polyA-enrichment was performed by the Core Unit Systems Medicine (University of Würzburg). Both species were sequenced on the Illumina NextSeq500 platform using a single high-output flow cell for the 8 *C. richardii* samples, and duplicate high-output flow cells for the 12 *P. vulgare* samples for 150 nt paired-end reads (300 cycles). All sequences are accessible in the GenBank Short Read Archive under the project accession PRJEB45027.

### Transcriptome assembly and annotation

Transcriptomes for *P. vulgare* and *C. richardii* were assembled using Trinity [84] with the trimmomatic option (Table S4). Coding regions were predicted with TransDecoder (https://github.com/TransDecoder/TransDecoder). Domains and GeneOntology annotations were predicted using InterPro [85]. To generate a level of annotation comparable to the fern data, all proteomes (see Table S5) were annotated de-novo using InterPro.

### Differential expression

For the ferns, reads were mapped onto the reference transcriptomes by Salmon [86]. For *A. thaliana* and *H. vulgare*, the reads were mapped onto the reference genomes (Table S5) by RSEM [87]. Differentially expressed genes were identified with DESeq2 [88].

### Evolutionary reconstruction

Orthology relationships over all species (Table S5) were predicted using Orthofinder [89]. The last common ancestor for each orthogroup was calculated using the Bio::TreeIO Perl package.

### Phylogenetic analyses for streptophyte SLAC and SnRK2 families

Genes were identified in the literature or by performing BLASTp searches using Arabidopsis protein sequences against the relevant genome or transcriptome assembly as indicated in Table S3, and confirmed with reciprocal BLASTp searches back against Arabidopsis and preliminary phylogenetic analyses. The maximum likelihood phylogenetic tree for the SLAC/SLAH family (Fig. S3) was calculated from a MAFFT alignment of full length predicted protein sequences using PhyML 3.0 at http://www.trex.uqam.ca/ with the JTT substitution model and 1000 bootstrap replicates [90]. The maximum likelihood phylogenetic tree for the SnRK2 family was calculated using PhyML 3.0 at http://www.atgc-montpellier.fr/phyml/ with SmartModel Selection [90, 91], and 1000 bootstrap replicates, from a MAFFT alignment of predicted protein sequences trimmed using Gblocks via the online server at http://molevol.cmima.csic.es/castresana/Gblocks_server.html [S3], with all options for reduced stringency selected. Full sequence and species details are given in Table S3.

### Cloning and complementary RNA generation

Full-length coding sequence from OST1, SLAC and/or CPK homologs of *C. richardii* (Hn-n, obtained from J. Banks; leaf), *P. vulgare* (Common Polypody-winter hardy; Westdijk’s, Kwekerijen; leaf), *P. abies* (plants maintained at the University of Tasmania; needles), *G. biloba* (plants maintained at the Würzburg Botanical Gardens; leaves), *A. trichopoda* (plants maintained at the University of Tasmania; leaves), *S. fallax* (‘MN’, obtained from S. Rensing, from an original sample from Dave Weston, Oak Ridge National Laboratory; gametophyte), *A. agrestis* (‘Bonn’, obtained from P. Szoevenyi; gametophyte) were isolated from cDNA generated from RNA from the tissues indicated using primers outlined in Table S6 and cloned into pNB1uYN and pNB1uYC expression vectors by uracil excision-based cloning [92]. The details for all *A. thaliana* constructs, and *C. richardii* CrSLAC1a, CrSLAC1b, CrGAIA1 and CrPGAI constructs have been published previously [9, 12-15, 31]. Where indicated, site-directed mutations were introduced using a modified USER fusion method, as previously described [93], using the primers outlined in Table S6. For functional analysis, complementary RNA (cRNA) was prepared with the AmpliCap-Max™ T7 High Yield Message Maker Kit (Epicentre Biotechnologies). Oocyte preparation and cRNA injection were performed as previously described [94]. For oocyte BiFC and electrophysiological experiments, 10 ng of each SLAC:YFP^CT^ (vector pNB1uYC) and 10 ng of each OST1:YFPNT or 5 ng of CPK:YFP^NT^ or AtCIPK23ΔEF:YFP^NT^ + AtCBL1:YFP^NT^ (vector pNB1uYN) cRNA, or cRNA of the same genes cloned into the pNB1u vector without YFP fragments, were injected into *Xenopus* oocytes.

### Double-electrode voltage-clamp studies

Oocytes were perfused with MES/Tris-based buffers, with standard solutions containing 10 mM MES/Tris (pH 5.6), 1 mM Ca(gluconate)_2_, 1 mM Mg(gluconate)_2_, 1 mM LaCl_3_ and 100 mM NaCl, NaNO_3_ or Na(gluconate). To balance the ionic strength, we compensated for changes in the chloride or nitrate concentration with Na(gluconate). For recording representative current traces standard voltage protocol was as follows: starting from a holding potential (VH) of 0 mV, single 20 s-voltage pulses were applied in 20 mV decrements from +40 to −180 mV. Instantaneous currents were extracted right after the voltage jump from the holding potential of 0 mV to 50 ms test pulses ranging from +70 to −150 mV.

## Acknowledgments

We thank Asst. Prof. Scott McAdam and Prof. Tim Brodribb for providing clones of *C. richardii* and *P. abies* SLAC homologs and fruitful discussion, Dr. Péter Szövényi and Prof. Jody Banks for early access to sequence data and provision of plant material, Dr. Andrew Plackett for early sharing of sequence information, and Prof. Stefan Rensing and Marco Göttig for provision of plant material. The support of the German Science Foundation (DFG) is acknowledged by M.R.G.R, J.S. and R.H. for the project ‘Evolution of molecular mechanisms that control stomatal closure’ (RO2381/8-1, SCHU2352/7-1 and HE1640/40-1), by D.B. and R.H. for the DFG priority programme 2237: ‘MAdLand -Molecular Adaptation to Land: plant evolution to change’, and by D.G. and R.H. for the CRC/TR166 ‘ReceptorLight’ project B8. P.A. and R.H. acknowledge funding by the Bavarian State Ministry of the Environment and Consumer Protection. K.M. acknowledges support from King Saud University’s (KSU) Distinguished Scientist Fellowship Research Program (DSFP). F.S. acknowledges support from the German Academic Exchange Service (DAAD) and an Australian Research Council Discovery Early Career Award (DE200101133) funded by the Australian Government.

## Author Contributions

F.S., T.M., J.H, L.V., C.L., M.M., H.M., M.B., and J.S. performed the research and analysed the data. F.S., P.A., K.M., D.B., M.R.G.R, D.G., J.S. and R.H. designed the study. F.S., M.R.G.R., D.G., J.S. and R.H. wrote the manuscript.

