## Supplemental Information for "Gaining or cutting SLAC: the evolution of plant guard cell signalling pathways"

**Table S1. Complete list of GO terms and annotated domains for orthogroups from Figure 1B.** Normalised counts are given for 'shared' orthogroups with significantly higher expression in guard cell than whole leaf samples in all angiosperm (*A. thaliana*, *H. vulgare*) and fern (*C. richardii*, *P. vulgare*) species examined ('shared'), both angiosperm but neither fern species (angiosperm-specific; 'angio'), and both fern but neither angiosperm (fern-specific; 'fern'). Counts were normalised to the total number of genes per orthogroup.

| GO | Total | Shared | Angio | Fern | | Domain | Total | Shared | Angio | Fern |
|---|---|---|---|---|---|---|---|---|---|---|
| ATP binding | 33.99 | 10.95 | 18.75 | 4.29 | | Protein kinase domain | 16.66 | 8.14 | 6.99 | 1.54 |
| oxidation-reduction process | 33.33 | 12.81 | 6.95 | 13.57 | | Protein tyrosine kinase | 7.95 | 2.82 | 5.13 | 0.00 |
| protein binding | 26.08 | 8.67 | 12.07 | 5.34 | | Cytochrome P450 | 7.46 | 3.90 | 1.00 | 2.55 |
| protein kinase activity | 24.46 | 10.92 | 12.00 | 1.54 | | Leucine rich repeat N-terminal domain | 6.12 | 3.95 | 2.17 | 0.00 |
| protein phosphorylation | 24.46 | 10.92 | 12.00 | 1.54 | | Leucine rich repeat | 5.98 | 3.49 | 2.48 | 0.00 |
| integral component of membrane | 22.29 | 9.53 | 7.30 | 5.47 | | ABC transporter | 5.57 | 2.74 | 1.83 | 1.00 |
| membrane | 18.91 | 10.19 | 7.71 | 1.01 | | Transmembrane amino acid transporter protein | 4.99 | 2.99 | 1.00 | 1.00 |
| transmembrane transport | 15.89 | 8.73 | 2.87 | 4.29 | | Multicopper oxidase | 4.63 | 2.00 | 0.00 | 2.63 |
| carbohydrate metabolic process | 13.57 | 10.58 | 1.02 | 1.98 | | Myb-like DNA-binding domain | 4.54 | 3.78 | 0.76 | 0.00 |
| oxidoreductase activity | 12.64 | 4.11 | 1.66 | 6.87 | | Helicase conserved C-terminal domain | 4.35 | 0.00 | 3.74 | 0.61 |
| regulation of transcription, DNA-templated | 10.13 | 6.86 | 2.60 | 0.67 | | C2 domain | 4.10 | 1.71 | 2.40 | 0.00 |
| hydrolase activity, hydrolyzing O-glycosyl compounds | 9.64 | 7.62 | 1.02 | 1.00 | | Mitochondrial carrier protein | 4.00 | 0.00 | 1.00 | 3.00 |
| catalytic activity | 9.56 | 2.18 | 5.14 | 2.25 | | OPT oligopeptide transporter protein | 3.97 | 1.00 | 0.00 | 2.97 |
| heme binding | 9.45 | 5.90 | 1.00 | 2.55 | | RNA recognition motif. (a.k.a. RRM, RBD, or RNP domain) | 3.63 | 0.98 | 1.89 | 0.75 |
| metabolic process | 8.89 | 0.72 | 2.05 | 6.12 | | short chain dehydrogenase | 3.49 | 0.87 | 0.97 | 1.66 |
| hydrolase activity | 8.40 | 2.76 | 2.85 | 2.79 | | haloacid dehalogenase-like hydrolase | 3.38 | 1.39 | 0.00 | 1.99 |
| DNA binding | 7.99 | 3.39 | 4.13 | 0.48 | | Helix-loop-helix DNA-binding domain | 3.36 | 2.53 | 0.00 | 0.83 |
| iron ion binding | 7.46 | 3.90 | 1.00 | 2.55 | | F-box domain | 3.35 | 0.00 | 1.62 | 1.74 |
| oxidoreductase activity, acting on paired donors, with incorporation or reduction of molecular oxygen | 7.46 | 3.90 | 1.00 | 2.55 | | Remorin, C-terminal region | 3.33 | 1.60 | 1.73 | 0.00 |
| metal ion binding | 7.10 | 3.61 | 2.20 | 1.29 | | Kinesin motor domain | 3.29 | 0.01 | 3.28 | 0.00 |
| nucleic acid binding | 7.01 | 1.00 | 3.24 | 2.77 | | AMP-binding enzyme | 2.99 | 0.00 | 2.02 | 0.97 |
| DNA-binding transcription factor activity | 6.48 | 4.90 | 0.93 | 0.65 | | Response regulator receiver domain | 2.95 | 1.76 | 0.00 | 1.19 |
| protein dimerization activity | 6.27 | 3.38 | 1.16 | 1.73 | | Leucine Rich Repeat | 2.72 | 1.46 | 1.26 | 0.00 |
| zinc ion binding | 5.92 | 2.45 | 2.54 | 0.93 | | Xylanase inhibitor N-terminal | 2.72 | 0.91 | 0.95 | 0.85 |
| ATPase activity | 5.57 | 2.74 | 1.83 | 1.00 | | Sodium/hydrogen exchanger family | 2.60 | 1.74 | 0.86 | 0.00 |
| nucleus | 5.00 | 2.14 | 2.86 | 0.00 | | Ankyrin repeats (3 copies) | 2.57 | 0.00 | 2.25 | 0.33 |
| proteolysis | 4.70 | 1.79 | 0.01 | 2.90 | | ABC-2 type transporter | 2.57 | 1.62 | 0.95 | 0.00 |
| cytoplasm | 4.61 | 1.00 | 2.79 | 0.82 | | Myb/SANT-like DNA-binding domain | 2.55 | 0.00 | 0.84 | 1.71 |
| copper ion binding | 4.32 | 1.99 | 0.00 | 2.34 | | DHHC palmitoyltransferase | 2.54 | 0.00 | 2.54 | 0.00 |
| sequence-specific DNA binding | 4.30 | 2.72 | 0.00 | 1.57 | | Xylanase inhibitor C-terminal | 2.54 | 0.90 | 0.71 | 0.93 |
| microtubule binding | 3.62 | 0.01 | 3.61 | 0.00 | | SNF2 family N-terminal domain | 2.49 | 0.00 | 1.75 | 0.74 |
| flavin adenine dinucleotide binding | 3.42 | 0.51 | 2.76 | 0.15 | | 2OG-Fe(II) oxygenase superfamily | 2.48 | 1.52 | 0.96 | 0.00 |
| microtubule-based movement | 3.29 | 0.01 | 3.28 | 0.00 | | Domain of unknown function (DUF588) | 2.47 | 0.00 | 0.84 | 1.64 |
| microtubule motor activity | 3.29 | 0.01 | 3.28 | 0.00 | | Histidine kinase-, DNA gyrase B-, and HSP90-like ATPase | 2.32 | 0.00 | 0.51 | 1.80 |
| transferase activity, transferring acyl groups other than amino-acyl groups | 3.23 | 1.93 | 0.65 | 0.64 | | X8 domain | 2.27 | 1.31 | 0.96 | 0.00 |

| GO | Total | Shared | Angio | Fern | | Domain | Total | Shared | Angio | Fern |
|---|---|---|---|---|---|---|---|---|---|---|
| transmembrane transporter activity | 3.23 | 1.91 | 1.01 | 0.31 | | Glycosyl hydrolases family 17 | 2.05 | 1.05 | 1.00 | 0.00 |
| transferase activity, transferring glycosyl groups | 2.96 | 2.02 | 0.00 | 0.94 | | Leucine Rich repeat | 2.02 | 0.83 | 0.65 | 0.53 |
| phosphorelay signal transduction system | 2.95 | 1.76 | 0.00 | 1.19 | | Glycosyl hydrolase family 9 | 2.00 | 1.00 | 0.00 | 1.00 |
| protein glycosylation | 2.76 | 1.99 | 0.77 | 0.00 | | MatE | 2.00 | 1.00 | 1.00 | 0.00 |
| solute:proton antiporter activity | 2.60 | 1.74 | 0.86 | 0.00 | | Major intrinsic protein | 2.00 | 1.00 | 0.00 | 1.00 |
| cation transport | 2.60 | 1.74 | 0.86 | 0.00 | | EF-hand domain pair | 2.00 | 1.00 | 0.00 | 1.00 |
| transcription, DNA-templated | 2.57 | 0.00 | 1.00 | 1.57 | | GDSL/SGNH-like Acyl-Esterase family found in Pmr5 and Cas1p | 2.00 | 0.99 | 1.00 | 0.00 |
| peroxidase activity | 2.51 | 2.51 | 0.00 | 0.00 | | Peroxidase | 2.00 | 2.00 | 0.00 | 0.00 |
| RNA binding | 2.36 | 0.00 | 1.51 | 0.85 | | GDSL-like Lipase/Acylhydrolase | 1.98 | 1.98 | 0.00 | 0.00 |
| response to stress | 2.18 | 1.05 | 1.13 | 0.00 | | Chalcone-flavanone isomerase | 1.96 | 0.00 | 1.96 | 0.00 |
| calcium ion binding | 2.17 | 1.03 | 0.00 | 1.15 | | Glycosyl hydrolases family 28 | 1.96 | 1.96 | 0.00 | 0.00 |
| intracellular | 2.16 | 0.40 | 0.93 | 0.83 | | Thioredoxin | 1.96 | 0.95 | 0.00 | 1.00 |
| translation | 2.09 | 0.17 | 0.93 | 1.00 | | No apical meristem (NAM) protein | 1.95 | 0.98 | 0.95 | 0.02 |
| ribosome | 2.09 | 0.17 | 0.93 | 1.00 | | Metalloenzyme superfamily | 1.94 | 0.00 | 1.00 | 0.94 |
| structural constituent of ribosome | 2.09 | 0.17 | 0.93 | 1.00 | | Sugar efflux transporter for intercellular exchange | 1.93 | 0.99 | 0.00 | 0.94 |
| magnesium ion binding | 2.03 | 1.00 | 0.00 | 1.03 | | Kelch motif | 1.92 | 0.00 | 0.67 | 1.25 |
| GTP binding | 2.01 | 0.01 | 1.00 | 1.00 | | Calcineurin-like phosphoesterase | 1.90 | 0.95 | 0.00 | 0.94 |
| glycolytic process | 2.00 | 0.00 | 0.00 | 2.00 | | Probable lipid transfer | 1.88 | 0.00 | 1.88 | 0.00 |
| drug transmembrane transporter activity | 2.00 | 1.00 | 1.00 | 0.00 | | Pollen allergen | 1.86 | 0.86 | 1.00 | 0.00 |
| antiporter activity | 2.00 | 1.00 | 1.00 | 0.00 | | Peptidase family M20/M25/M40 | 1.84 | 0.00 | 0.00 | 1.84 |
| drug transmembrane transport | 2.00 | 1.00 | 1.00 | 0.00 | | Formin Homology 2 Domain | 1.79 | 0.00 | 1.79 | 0.00 |
| channel activity | 2.00 | 1.00 | 0.00 | 1.00 | | Glutathione S-transferase, N-terminal domain | 1.77 | 0.81 | 0.96 | 0.00 |
| response to oxidative stress | 2.00 | 2.00 | 0.00 | 0.00 | | Galactosyltransferase | 1.76 | 0.99 | 0.77 | 0.00 |
| trehalose biosynthetic process | 1.99 | 0.99 | 1.00 | 0.00 | | WD domain, G-beta repeat | 1.75 | 0.90 | 0.85 | 0.00 |
| isomerase activity | 1.98 | 1.00 | 0.00 | 0.98 | | Protein of unknown function (DUF620) | 1.73 | 1.00 | 0.00 | 0.73 |
| hydrolase activity, acting on ester bonds | 1.98 | 1.98 | 0.00 | 0.00 | | Cytochrome b5-like Heme/Steroid binding domain | 1.73 | 0.00 | 0.94 | 0.79 |
| intramolecular lyase activity | 1.96 | 0.00 | 1.96 | 0.00 | | Zinc finger, C3HC4 type (RING finger) | 1.72 | 0.60 | 0.94 | 0.18 |
| proton transmembrane transporter activity | 1.96 | 0.96 | 1.00 | 0.00 | | Trehalose-phosphatase | 1.71 | 0.99 | 0.71 | 0.00 |
| polygalacturonase activity | 1.96 | 1.96 | 0.00 | 0.00 | | NUDIX domain | 1.70 | 0.82 | 0.88 | 0.00 |
| cell redox homeostasis | 1.96 | 0.95 | 0.00 | 1.00 | | Lytic transglycolase | 1.69 | 0.87 | 0.82 | 0.00 |
| GTPase activity | 1.89 | 0.01 | 1.00 | 0.88 | | GRAM domain | 1.69 | 0.92 | 0.78 | 0.00 |
| cysteine-type peptidase activity | 1.86 | 0.99 | 0.00 | 0.86 | | Ring finger domain | 1.68 | 0.89 | 0.32 | 0.46 |
| calmodulin binding | 1.85 | 0.00 | 1.85 | 0.00 | | AP2 domain | 1.67 | 1.67 | 0.00 | 0.00 |
| galactosyltransferase activity | 1.76 | 0.99 | 0.77 | 0.00 | | GMC oxidoreductase | 1.64 | 0.00 | 0.92 | 0.72 |
| cell wall | 1.74 | 1.74 | 0.00 | 0.00 | | Ribonuclease III domain | 1.63 | 0.00 | 0.69 | 0.94 |
| RNA processing | 1.65 | 0.00 | 0.70 | 0.95 | | PUB domain | 1.62 | 0.00 | 1.62 | 0.00 |
| oxidoreductase activity, acting on CH-OH group of donors | 1.64 | 0.00 | 0.92 | 0.72 | | Late embryogenesis abundant protein | 1.61 | 1.10 | 0.00 | 0.51 |
| ribonuclease III activity | 1.63 | 0.00 | 0.70 | 0.94 | | E1-E2 ATPase | 1.60 | 0.77 | 0.00 | 0.82 |
| defense response | 1.57 | 0.00 | 1.00 | 0.57 | | Transferase family | 1.57 | 0.93 | 0.00 | 0.64 |
| N,N-dimethylaniline monooxygenase activity | 1.51 | 0.51 | 1.00 | 0.00 | | Seed dormancy control | 1.57 | 0.00 | 0.00 | 1.57 |
| NADP binding | 1.51 | 0.51 | 1.00 | 0.00 | | GDP-mannose 4,6 dehydratase | 1.54 | 0.66 | 0.89 | 0.00 |

| GO | Total | Shared | Angio | Fern | | Domain | Total | Shared | Angio | Fern |
|---|---|---|---|---|---|---|---|---|---|---|
| serine-type endopeptidase activity | 1.51 | 0.80 | 0.72 | 0.00 | | Glycosyl hydrolase family 3 N terminal domain | 1.52 | 1.52 | 0.00 | 0.00 |
| lipid metabolic process | 1.49 | 0.57 | 0.91 | 0.01 | | IQ calmodulin-binding motif | 1.52 | 0.68 | 0.84 | 0.00 |
| coenzyme binding | 1.42 | 0.30 | 0.11 | 1.00 | | Flavin-binding monooxygenase-like | 1.51 | 0.51 | 1.00 | 0.00 |
| helicase activity | 1.35 | 0.00 | 1.35 | 0.00 | | CBS domain | 1.51 | 0.00 | 0.00 | 1.51 |
| mitochondrion | 1.25 | 0.00 | 1.00 | 0.25 | | Protein of unknown function (DUF740) | 1.50 | 0.00 | 1.50 | 0.00 |
| lipid binding | 1.24 | 0.67 | 0.55 | 0.01 | | Glycosyl hydrolase family 3 C-terminal domain | 1.48 | 1.48 | 0.00 | 0.00 |
| hydrolase activity, acting on glycosyl bonds | 1.20 | 1.20 | 0.00 | 0.00 | | non-haem dioxygenase in morphine synthesis N-terminal | 1.45 | 0.51 | 0.93 | 0.00 |
| metal ion transport | 1.05 | 0.89 | 0.00 | 0.17 | | Tetratricopeptide repeat | 1.40 | 0.00 | 0.59 | 0.81 |
| serine-type carboxypeptidase activity | 1.04 | 0.00 | 0.00 | 1.04 | | UBA/TS-N domain | 1.40 | 0.00 | 1.40 | 0.00 |
| NAD binding | 1.04 | 0.01 | 0.03 | 1.00 | | Fasciclin domain | 1.38 | 0.90 | 0.00 | 0.48 |
| carbohydrate binding | 1.02 | 1.02 | 0.00 | 0.00 | | Anaphase-promoting complex subunit 4 WD40 domain | 1.35 | 0.93 | 0.42 | 0.00 |
| translational initiation | 1.01 | 0.00 | 0.01 | 1.00 | | Glutathione S-transferase, C-terminal domain | 1.35 | 0.92 | 0.43 | 0.00 |
| translation initiation factor activity | 1.01 | 0.00 | 0.01 | 1.00 | | Serine aminopeptidase, S33 | 1.33 | 1.00 | 0.33 | 0.00 |
| serine-type peptidase activity | 1.01 | 0.00 | 0.01 | 1.00 | | Protein phosphatase 2C | 1.32 | 0.00 | 0.99 | 0.33 |
| ATP hydrolysis coupled proton transport | 1.01 | 0.01 | 1.00 | 0.00 | | DEAD/DEAH box helicase | 1.32 | 0.00 | 1.32 | 0.00 |
| fatty acid biosynthetic process | 1.00 | 1.00 | 0.00 | 0.00 | | Universal stress protein family | 1.26 | 1.05 | 0.20 | 0.00 |
| carboxy-lyase activity | 1.00 | 0.00 | 0.00 | 1.00 | | START domain | 1.24 | 0.67 | 0.55 | 0.01 |
| hydroxymethylglutaryl-CoA reductase (NADPH) activity | 1.00 | 0.00 | 0.00 | 1.00 | | B-box zinc finger | 1.23 | 0.40 | 0.00 | 0.83 |
| coenzyme A metabolic process | 1.00 | 0.00 | 0.00 | 1.00 | | AMP-binding enzyme C-terminal domain | 1.16 | 0.00 | 1.07 | 0.09 |
| DNA topological change | 1.00 | 0.01 | 0.99 | 0.00 | | S1 RNA binding domain | 1.06 | 0.00 | 0.06 | 1.00 |
| alternative oxidase activity | 1.00 | 0.00 | 0.00 | 1.00 | | Heavy-metal-associated domain | 1.05 | 0.89 | 0.00 | 0.17 |
| protein peptidyl-prolyl isomerization | 1.00 | 0.00 | 0.00 | 1.00 | | Serine carboxypeptidase | 1.04 | 0.00 | 0.00 | 1.04 |
| phosphoenolpyruvate carboxykinase (ATP) activity | 1.00 | 0.00 | 0.00 | 1.00 | | Salt stress response/antifungal | 1.03 | 0.07 | 0.97 | 0.00 |
| cellular amino acid metabolic process | 1.00 | 0.00 | 0.00 | 1.00 | | C2H2-type zinc finger | 1.03 | 0.93 | 0.00 | 0.09 |
| pyruvate kinase activity | 1.00 | 0.00 | 0.00 | 1.00 | | Peptidase dimerisation domain | 1.02 | 0.00 | 0.00 | 1.02 |
| fatty acid metabolic process | 1.00 | 0.00 | 0.00 | 1.00 | | Glycosyl hydrolase family 1 | 1.02 | 1.00 | 0.02 | 0.00 |
| peptidyl-prolyl cis-trans isomerase activity | 1.00 | 0.00 | 0.00 | 1.00 | | Dirigent-like protein | 1.02 | 0.95 | 0.06 | 0.00 |
| phosphoglycerate kinase activity | 1.00 | 0.00 | 0.00 | 1.00 | | Aldehyde dehydrogenase family | 1.01 | 0.01 | 0.00 | 1.00 |
| acyl-[acyl-carrier-protein] desaturase activity | 1.00 | 0.00 | 0.00 | 1.00 | | Enoyl-(Acyl carrier protein) reductase | 1.01 | 1.01 | 0.00 | 0.00 |
| intramolecular transferase activity, phosphotransferases | 1.00 | 0.00 | 0.00 | 1.00 | | D-isomer specific 2-hydroxyacid dehydrogenase, NAD binding domain | 1.01 | 0.01 | 0.00 | 1.00 |
| gluconeogenesis | 1.00 | 0.00 | 0.00 | 1.00 | | Fatty acid desaturase | 1.01 | 0.01 | 0.00 | 1.00 |
| protein retention in ER lumen | 1.00 | 0.00 | 0.00 | 1.00 | | PAZ domain | 1.01 | 0.00 | 0.59 | 0.42 |
| ER retention sequence binding | 1.00 | 0.00 | 0.00 | 1.00 | | Jacalin-like lectin domain | 1.01 | 0.02 | 0.99 | 0.00 |
| potassium ion binding | 1.00 | 0.00 | 0.00 | 1.00 | | Ras family | 1.01 | 0.01 | 1.00 | 0.00 |
| ATPase activator activity | 1.00 | 0.00 | 0.00 | 1.00 | | Raffinose synthase or seed imbibition protein Sip1 | 1.01 | 0.00 | 1.00 | 0.01 |
| chaperone binding | 1.00 | 0.00 | 0.00 | 1.00 | | Pectinacetylesterase | 1.01 | 1.00 | 0.01 | 0.00 |
| ammonium transport | 1.00 | 0.00 | 1.00 | 0.00 | | GRAS domain family | 1.00 | 0.00 | 1.00 | 0.00 |
| alpha-mannosidase activity | 1.00 | 0.00 | 1.00 | 0.00 | | Glycosyl transferase family 90 | 1.00 | 1.00 | 0.00 | 0.00 |
| ammonium transmembrane transporter activity | 1.00 | 0.00 | 1.00 | 0.00 | | Thaumatin family | 1.00 | 1.00 | 0.00 | 0.00 |
| intermembrane lipid transfer | 1.00 | 0.00 | 1.00 | 0.00 | | Ubiquitin-conjugating enzyme | 1.00 | 1.00 | 0.00 | 0.00 |

| GO | Total | Shared | Angio | Fern | | Domain | Total | Shared | Angio | Fern |
|---|---|---|---|---|---|---|---|---|---|---|
| intermembrane lipid transfer activity | 1.00 | 0.00 | 1.00 | 0.00 | | Hydroxymethylglutaryl-coenzyme A reductase | 1.00 | 0.00 | 0.00 | 1.00 |
| proton-transporting V-type ATPase, V0 domain | 1.00 | 0.00 | 1.00 | 0.00 | | Serine carboxypeptidase S28 | 1.00 | 0.00 | 0.00 | 1.00 |
| superoxide metabolic process | 1.00 | 0.00 | 1.00 | 0.00 | | Ribosomal S3Ae family | 1.00 | 0.00 | 0.00 | 1.00 |
| superoxide dismutase activity | 1.00 | 0.00 | 1.00 | 0.00 | | Phenolic acid decarboxylase (PAD) | 1.00 | 0.00 | 0.00 | 1.00 |
| coproporphyrinogen oxidase activity | 1.00 | 0.00 | 1.00 | 0.00 | | CVNH domain | 1.00 | 0.00 | 0.00 | 1.00 |
| NADH dehydrogenase activity | 1.00 | 0.00 | 1.00 | 0.00 | | Alternative oxidase | 1.00 | 0.00 | 0.00 | 1.00 |
| NADH dehydrogenase (ubiquinone) activity | 1.00 | 0.00 | 1.00 | 0.00 | | Ran-interacting Mog1 protein | 1.00 | 0.00 | 0.00 | 1.00 |
| mannose metabolic process | 1.00 | 0.00 | 1.00 | 0.00 | | Pyruvate kinase, barrel domain | 1.00 | 0.00 | 0.00 | 1.00 |
| aspartic-type endopeptidase activity | 1.00 | 0.00 | 1.00 | 0.00 | | Phosphoenolpyruvate carboxykinase | 1.00 | 0.00 | 0.00 | 1.00 |
| GTPase activator activity | 1.00 | 0.00 | 1.00 | 0.00 | | Protein of unknown function, DUF599 | 1.00 | 0.00 | 0.00 | 1.00 |
| porphyrin-containing compound biosynthetic process | 1.00 | 0.00 | 1.00 | 0.00 | | ER lumen protein retaining receptor | 1.00 | 0.00 | 0.00 | 1.00 |
| molybdate ion transport | 1.00 | 0.00 | 1.00 | 0.00 | | Activator of Hsp90 ATPase, N-terminal | 1.00 | 0.00 | 0.00 | 1.00 |
| molybdate ion transmembrane transporter activity | 1.00 | 0.00 | 1.00 | 0.00 | | Protein of unknown function (DUF1336) | 1.00 | 0.00 | 0.00 | 1.00 |
| galactoside 2-alpha-L-fucosyltransferase activity | 1.00 | 1.00 | 0.00 | 0.00 | | Pyridoxamine 5'-phosphate oxidase | 1.00 | 0.00 | 0.00 | 1.00 |
| inorganic diphosphatase activity | 1.00 | 1.00 | 0.00 | 0.00 | | Phosphoglycerate kinase | 1.00 | 0.00 | 0.00 | 1.00 |
| positive regulation of circadian rhythm | 1.00 | 1.00 | 0.00 | 0.00 | | Cyclophilin type peptidyl-prolyl cis-trans isomerase/CLD | 1.00 | 0.00 | 0.00 | 1.00 |
| cell wall biogenesis | 1.00 | 1.00 | 0.00 | 0.00 | | Gamma-glutamyltranspeptidase | 1.00 | 0.00 | 0.00 | 1.00 |
| phosphate-containing compound metabolic process | 1.00 | 1.00 | 0.00 | 0.00 | | Translation initiation factor SUI1 | 1.00 | 0.00 | 0.00 | 1.00 |
| cellulose biosynthetic process | 1.00 | 0.00 | 1.00 | 0.00 | | Embryo-specific protein 3, (ATS3) | 1.00 | 0.00 | 1.00 | 0.00 |
| cellulose synthase (UDP-forming) activity | 1.00 | 0.00 | 1.00 | 0.00 | | HORMA domain | 1.00 | 0.00 | 1.00 | 0.00 |
| DNA topoisomerase type II (ATP-hydrolyzing) activity | 0.99 | 0.00 | 0.99 | 0.00 | | Permease family | 1.00 | 0.00 | 1.00 | 0.00 |
| malate transport | 0.99 | 0.99 | 0.00 | 0.00 | | TLC domain | 1.00 | 0.00 | 1.00 | 0.00 |
| regulation of cyclin-dependent protein serine/threonine kinase activity | 0.99 | 0.99 | 0.00 | 0.00 | | Cytochrome c oxidase biogenesis protein Cmc1 like | 1.00 | 0.00 | 1.00 | 0.00 |
| protein kinase binding | 0.99 | 0.99 | 0.00 | 0.00 | | Protein of unknown function (DUF616) | 1.00 | 0.00 | 1.00 | 0.00 |
| peptidyl-pyroglutamic acid biosynthetic process, using glutaminyl-peptide cyclotransferase | 0.99 | 0.00 | 0.00 | 0.99 | | Ammonium Transporter Family | 1.00 | 0.00 | 1.00 | 0.00 |
| glutaminyl-peptide cyclotransferase activity | 0.99 | 0.00 | 0.00 | 0.99 | | RNA polymerase Rpb3/Rpb11 dimerisation domain | 1.00 | 0.00 | 1.00 | 0.00 |
| acetylglucosaminyltransferase activity | 0.99 | 0.00 | 0.99 | 0.00 | | Glycolipid transfer protein (GLTP) | 1.00 | 0.00 | 1.00 | 0.00 |
| pectinesterase activity | 0.99 | 0.99 | 0.00 | 0.00 | | Sugar-tranasporters, 12 TM | 1.00 | 0.00 | 1.00 | 0.00 |
| cell wall modification | 0.99 | 0.99 | 0.00 | 0.00 | | Coproporphyrinogen III oxidase | 1.00 | 0.00 | 1.00 | 0.00 |
| anion transport | 0.98 | 0.98 | 0.00 | 0.00 | | Protein of unknown function (DUF3339) | 1.00 | 0.00 | 1.00 | 0.00 |
| ribonuclease T2 activity | 0.98 | 0.00 | 0.98 | 0.00 | | GDA1/CD39 (nucleoside phosphatase) family | 1.00 | 0.00 | 1.00 | 0.00 |
| chloroplast | 0.98 | 0.00 | 0.00 | 0.98 | | Predicted membrane protein (DUF2053) | 1.00 | 0.00 | 1.00 | 0.00 |
| response to auxin | 0.98 | 0.98 | 0.00 | 0.00 | | Glucose / Sorbosone dehydrogenase | 1.00 | 0.00 | 1.00 | 0.00 |
| Rho guanyl-nucleotide exchange factor activity | 0.98 | 0.98 | 0.00 | 0.00 | | Signal peptide peptidase | 1.00 | 0.00 | 1.00 | 0.00 |
| red chlorophyll catabolite reductase activity | 0.98 | 0.00 | 0.00 | 0.98 | | Ndr family | 1.00 | 0.00 | 1.00 | 0.00 |
| polysaccharide catabolic process | 0.98 | 0.98 | 0.00 | 0.00 | | Protein of unknown function (DUF1666) | 1.00 | 0.00 | 1.00 | 0.00 |

| GO | Total | Shared | Angio | Fern | | Domain | Total | Shared | Angio | Fern |
|---|---|---|---|---|---|---|---|---|---|---|
| beta-amylase activity | 0.98 | 0.98 | 0.00 | 0.00 | | NADH-ubiquinone oxidoreductase B18 subunit (NDUFB7) | 1.00 | 0.00 | 1.00 | 0.00 |
| ATPase activity, coupled to transmembrane movement of substances | 0.97 | 0.97 | 0.00 | 0.00 | | Serine incorporator (Serinc) | 1.00 | 0.00 | 1.00 | 0.00 |
| nutrient reservoir activity | 0.97 | 0.97 | 0.00 | 0.00 | | ATP synthase subunit H | 1.00 | 0.00 | 1.00 | 0.00 |
| protein transport | 0.97 | 0.97 | 0.00 | 0.00 | | Rapid ALkalinization Factor (RALF) | 1.00 | 0.00 | 1.00 | 0.00 |
| motor activity | 0.97 | 0.00 | 0.97 | 0.00 | | Putative GTPase activating protein for Arf | 1.00 | 0.00 | 1.00 | 0.00 |
| myosin complex | 0.97 | 0.00 | 0.97 | 0.00 | | Caleosin related protein | 1.00 | 1.00 | 0.00 | 0.00 |
| electron transfer activity | 0.96 | 0.93 | 0.03 | 0.00 | | Xyloglucan fucosyltransferase | 1.00 | 1.00 | 0.00 | 0.00 |
| mitochondrial proton-transporting ATP synthase complex, coupling factor F(o) | 0.95 | 0.95 | 0.00 | 0.00 | | Inorganic pyrophosphatase | 1.00 | 1.00 | 0.00 | 0.00 |
| ATP synthesis coupled proton transport | 0.95 | 0.95 | 0.00 | 0.00 | | FAE1/Type III polyketide synthase-like protein | 1.00 | 1.00 | 0.00 | 0.00 |
| metallopeptidase activity | 0.94 | 0.00 | 0.94 | 0.00 | | Glycosyl transferase family 8 | 1.00 | 1.00 | 0.00 | 0.00 |
| oxidoreductase activity, acting on single donors with incorporation of molecular oxygen, incorporation of two atoms of oxygen | 0.94 | 0.94 | 0.00 | 0.00 | | Domain of unknown function (DUF3511) | 1.00 | 1.00 | 0.00 | 0.00 |
| oxidoreductase activity, acting on the CH-CH group of donors | 0.91 | 0.00 | 0.91 | 0.00 | | Exostosin family | 1.00 | 1.00 | 0.00 | 0.00 |
| diacylglycerol O-acyltransferase activity | 0.91 | 0.00 | 0.91 | 0.00 | | Purine nucleobase transmembrane transport | 1.00 | 1.00 | 0.00 | 0.00 |
| lipid storage | 0.90 | 0.00 | 0.00 | 0.90 | | Reticulon | 1.00 | 1.00 | 0.00 | 0.00 |
| biosynthetic process | 0.89 | 0.81 | 0.00 | 0.08 | | Protein of unknown function (DUF1313) | 1.00 | 1.00 | 0.00 | 0.00 |
| histone-lysine N-methyltransferase activity | 0.89 | 0.00 | 0.89 | 0.00 | | Cellulose synthase | 1.00 | 0.00 | 1.00 | 0.00 |
| peptide biosynthetic process | 0.87 | 0.00 | 0.87 | 0.00 | | Mlo family | 1.00 | 0.00 | 1.00 | 0.00 |
| oxidoreductase activity, acting on the CH-OH group of donors, NAD or NADP as acceptor | 0.86 | 0.00 | 0.00 | 0.86 | | Aldose 1-epimerase | 1.00 | 1.00 | 0.00 | 0.00 |
| 2,3-bisphosphoglycerate-independent phosphoglycerate mutase activity | 0.86 | 0.00 | 0.86 | 0.00 | | Papain family cysteine protease | 0.99 | 0.99 | 0.00 | 0.00 |
| cytokinin metabolic process | 0.84 | 0.00 | 0.84 | 0.00 | | AWPM-19-like family | 0.99 | 0.99 | 0.00 | 0.00 |
| cytokinin dehydrogenase activity | 0.84 | 0.00 | 0.84 | 0.00 | | Aluminium activated malate transporter | 0.99 | 0.99 | 0.00 | 0.00 |
| translational elongation | 0.84 | 0.00 | 0.84 | 0.00 | | Histone-like transcription factor (CBF/NF-Y) and archaeal histone | 0.99 | 0.99 | 0.00 | 0.00 |
| translation elongation factor activity | 0.84 | 0.00 | 0.84 | 0.00 | | Ubiquitin family | 0.99 | 0.99 | 0.00 | 0.00 |
| phosphoglycerate mutase activity | 0.82 | 0.00 | 0.00 | 0.82 | | Regulator of chromosome condensation (RCC1) repeat | 0.99 | 0.00 | 0.99 | 0.00 |
| manganese ion binding | 0.82 | 0.00 | 0.00 | 0.82 | | Protein of unknown function (DUF1191) | 0.99 | 0.00 | 0.00 | 0.99 |
| glucose catabolic process | 0.82 | 0.00 | 0.00 | 0.82 | | Uncharacterized protein family, UPF0114 | 0.99 | 0.00 | 0.00 | 0.99 |
| nucleoside metabolic process | 0.82 | 0.00 | 0.82 | 0.00 | | Cyclin | 0.99 | 0.99 | 0.00 | 0.00 |
| acid phosphatase activity | 0.81 | 0.81 | 0.00 | 0.00 | | Glutamine cyclotransferase | 0.99 | 0.00 | 0.00 | 0.99 |
| catechol oxidase activity | 0.81 | 0.00 | 0.00 | 0.81 | | Iron/manganese superoxide dismutases, C-terminal domain | 0.99 | 0.00 | 0.99 | 0.00 |
| strictosidine synthase activity | 0.81 | 0.81 | 0.00 | 0.00 | | Core-2/I-Branching enzyme | 0.99 | 0.00 | 0.99 | 0.00 |
| histone lysine methylation | 0.80 | 0.00 | 0.80 | 0.00 | | AUX/IAA family | 0.99 | 0.99 | 0.00 | 0.00 |
| signal transduction | 0.78 | 0.01 | 0.01 | 0.76 | | Pectinesterase | 0.99 | 0.99 | 0.00 | 0.00 |
| phosphorelay sensor kinase activity | 0.76 | 0.00 | 0.00 | 0.76 | | Protein of unknown function (DUF789) | 0.98 | 0.98 | 0.00 | 0.00 |
| xyloglucan:xyloglucosyl transferase activity | 0.75 | 0.75 | 0.00 | 0.00 | | Coiled coil protein 84 | 0.98 | 0.00 | 0.98 | 0.00 |
| cellular glucan metabolic process | 0.75 | 0.75 | 0.00 | 0.00 | | HCO3- transporter family | 0.98 | 0.98 | 0.00 | 0.00 |

| GO | Total | Shared | Angio | Fern | | Domain | Total | Shared | Angio | Fern |
|---|---|---|---|---|---|---|---|---|---|---|
| apoplast | 0.75 | 0.75 | 0.00 | 0.00 | | PPPDE putative peptidase domain | 0.98 | 0.98 | 0.00 | 0.00 |
| transferase activity, transferring acyl groups | 0.70 | 0.70 | 0.00 | 0.00 | | Pollen proteins Ole e I like | 0.98 | 0.00 | 0.98 | 0.00 |
| ionotropic glutamate receptor activity | 0.69 | 0.69 | 0.00 | 0.00 | | Gamma-thionin family | 0.98 | 0.00 | 0.98 | 0.00 |
| transcription factor complex | 0.65 | 0.00 | 0.00 | 0.65 | | Zinc finger C-x8-C-x5-C-x3-H type (and similar) | 0.98 | 0.98 | 0.00 | 0.00 |
| phosphorylation | 0.61 | 0.00 | 0.61 | 0.00 | | RNA polymerase I specific transcription initiation factor RRN3 | 0.98 | 0.00 | 0.98 | 0.00 |
| kinase activity | 0.61 | 0.00 | 0.61 | 0.00 | | Calcium-activated chloride channel | 0.98 | 0.00 | 0.98 | 0.00 |
| methyltransferase activity | 0.61 | 0.00 | 0.01 | 0.60 | | Ribonuclease T2 family | 0.98 | 0.00 | 0.98 | 0.00 |
| ADP binding | 0.58 | 0.00 | 0.00 | 0.58 | | Histidine phosphatase superfamily (branch 2) | 0.98 | 0.00 | 0.98 | 0.00 |
| protein folding | 0.58 | 0.58 | 0.00 | 0.00 | | Allene oxide cyclase | 0.98 | 0.00 | 0.00 | 0.98 |
| glycerolipid biosynthetic process | 0.57 | 0.00 | 0.57 | 0.00 | | Wound-induced protein WI12 | 0.98 | 0.00 | 0.00 | 0.98 |
| response to biotic stimulus | 0.57 | 0.00 | 0.00 | 0.57 | | Histidine phosphatase superfamily (branch 1) | 0.98 | 0.00 | 0.00 | 0.98 |
| organic substance metabolic process | 0.53 | 0.00 | 0.00 | 0.53 | | Auxin responsive protein | 0.98 | 0.98 | 0.00 | 0.00 |
| endoribonuclease activity, producing 5'-phosphomonoesters | 0.52 | 0.00 | 0.52 | 0.00 | | PRONE (Plant-specific Rop nucleotide exchanger) | 0.98 | 0.98 | 0.00 | 0.00 |
| oxidoreductase activity, acting on NAD(P)H, oxygen as acceptor | 0.51 | 0.51 | 0.00 | 0.00 | | Red chlorophyll catabolite reductase (RCC reductase) | 0.98 | 0.00 | 0.00 | 0.98 |
| sulfotransferase activity | 0.43 | 0.43 | 0.00 | 0.00 | | Protein of unknown function (DUF3537) | 0.98 | 0.98 | 0.00 | 0.00 |
| cytoskeleton organization | 0.42 | 0.42 | 0.00 | 0.00 | | Protein of unknown function, DUF604 | 0.98 | 0.98 | 0.00 | 0.00 |
| actin binding | 0.42 | 0.42 | 0.00 | 0.00 | | Glycosyl hydrolase family 14 | 0.98 | 0.98 | 0.00 | 0.00 |
| DNA repair | 0.30 | 0.00 | 0.30 | 0.00 | | Neprosin | 0.98 | 0.98 | 0.00 | 0.00 |
| plasma membrane | 0.29 | 0.00 | 0.00 | 0.29 | | Haemolysin-III related | 0.98 | 0.98 | 0.00 | 0.00 |
| enzyme inhibitor activity | 0.27 | 0.27 | 0.00 | 0.00 | | CRAL/TRIO domain | 0.98 | 0.00 | 0.00 | 0.98 |
| ATPase inhibitor activity | 0.25 | 0.00 | 0.00 | 0.25 | | LSM domain | 0.97 | 0.00 | 0.00 | 0.97 |
| negative regulation of ATPase activity | 0.25 | 0.00 | 0.00 | 0.25 | | ABC transporter transmembrane region | 0.97 | 0.97 | 0.00 | 0.00 |
| damaged DNA binding | 0.24 | 0.00 | 0.24 | 0.00 | | Protein of unknown function (DUF632) | 0.97 | 0.97 | 0.00 | 0.00 |
| nuclear chromosome | 0.23 | 0.00 | 0.00 | 0.23 | | Hydrophobic seed protein | 0.97 | 0.00 | 0.97 | 0.00 |
| pyridoxal phosphate binding | 0.09 | 0.00 | 0.00 | 0.08 | | PH domain | 0.97 | 0.00 | 0.97 | 0.00 |
| vesicle-mediated transport | 0.08 | 0.00 | 0.00 | 0.08 | | Cupin | 0.97 | 0.97 | 0.00 | 0.00 |
| oxidoreductase activity, acting on the aldehyde or oxo group of donors, NAD or NADP as acceptor | 0.06 | 0.00 | 0.06 | 0.00 | | TB2/DP1, HVA22 family | 0.97 | 0.97 | 0.00 | 0.00 |
| recognition of pollen | 0.06 | 0.06 | 0.00 | 0.00 | | SCAMP family | 0.97 | 0.97 | 0.00 | 0.00 |
| identical protein binding | 0.06 | 0.00 | 0.00 | 0.06 | | N terminus of Rad21 / Rec8 like protein | 0.97 | 0.00 | 0.00 | 0.97 |
| DNA integration | 0.05 | 0.01 | 0.02 | 0.02 | | Myosin head (motor domain) | 0.97 | 0.00 | 0.97 | 0.00 |
| extracellularly glutamate-gated ion channel activity | 0.05 | 0.05 | 0.00 | 0.00 | | Glycosyl hydrolase family 79, N-terminal domain | 0.96 | 0.96 | 0.00 | 0.00 |
| chromatin binding | 0.04 | 0.00 | 0.04 | 0.00 | | Glycosyl hydrolases family 16 | 0.96 | 0.96 | 0.00 | 0.00 |
| polysaccharide binding | 0.03 | 0.03 | 0.00 | 0.00 | | PAP2 superfamily | 0.96 | 0.00 | 0.96 | 0.00 |
| iron-sulfur cluster binding | 0.03 | 0.00 | 0.03 | 0.00 | | Plant phosphoribosyltransferase C-terminal | 0.96 | 0.96 | 0.00 | 0.00 |
| lyase activity | 0.03 | 0.00 | 0.00 | 0.03 | | SET domain | 0.96 | 0.00 | 0.71 | 0.25 |
| terpene synthase activity | 0.03 | 0.00 | 0.00 | 0.03 | | TspO/MBR family | 0.95 | 0.00 | 0.00 | 0.95 |
| ubiquitin-dependent protein catabolic process | 0.03 | 0.01 | 0.02 | 0.00 | | Oxysterol-binding protein | 0.95 | 0.00 | 0.00 | 0.95 |
| ATP-binding cassette (ABC) transporter complex | 0.03 | 0.00 | 0.00 | 0.03 | | Amino acid permease | 0.95 | 0.95 | 0.00 | 0.00 |
| transcription factor binding | 0.02 | 0.02 | 0.00 | 0.00 | | Mitochondrial ATP synthase g subunit | 0.95 | 0.95 | 0.00 | 0.00 |

| GO | Total | Shared | Angio | Fern | | Domain | Total | Shared | Angio | Fern |
|---|---|---|---|---|---|---|---|---|---|---|
| nucleosome | 0.02 | 0.00 | 0.02 | 0.00 | | PQQ-like domain | 0.95 | 0.00 | 0.00 | 0.95 |
| nucleotidyltransferase activity | 0.02 | 0.00 | 0.00 | 0.02 | | WRKY DNA -binding domain | 0.95 | 0.95 | 0.00 | 0.00 |
| plant-type cell wall organization | 0.02 | 0.00 | 0.02 | 0.00 | | POT family | 0.95 | 0.95 | 0.00 | 0.00 |
| structural constituent of cell wall | 0.02 | 0.00 | 0.02 | 0.00 | | Dof domain, zinc finger | 0.95 | 0.95 | 0.00 | 0.00 |
| DNA recombination | 0.01 | 0.00 | 0.00 | 0.01 | | DNA gyrase/topoisomerase IV, subunit A | 0.95 | 0.00 | 0.95 | 0.00 |
| aromatic amino acid family biosynthetic process | 0.01 | 0.01 | 0.00 | 0.00 | | NPH3 family | 0.94 | 0.94 | 0.00 | 0.00 |
| 3-deoxy-7-phosphoheptulonate synthase activity | 0.01 | 0.01 | 0.00 | 0.00 | | Pirin C-terminal cupin domain | 0.94 | 0.00 | 0.94 | 0.00 |
| exocytosis | 0.01 | 0.00 | 0.01 | 0.00 | | Tim10/DDP family zinc finger | 0.94 | 0.00 | 0.94 | 0.00 |
| vesicle docking | 0.01 | 0.00 | 0.01 | 0.00 | | cAMP-regulated phosphoprotein/endosulfine conserved region | 0.94 | 0.94 | 0.00 | 0.00 |
| ubiquitin-like modifier activating enzyme activity | 0.01 | 0.00 | 0.01 | 0.00 | | CRT-like, chloroquine-resistance transporter-like | 0.94 | 0.00 | 0.00 | 0.94 |
| aldehyde-lyase activity | 0.01 | 0.00 | 0.00 | 0.01 | | Amidohydrolase family | 0.94 | 0.00 | 0.94 | 0.00 |
| DNA topoisomerase type I activity | 0.01 | 0.01 | 0.00 | 0.00 | | Domain of unknown function (DUF4220) | 0.94 | 0.00 | 0.94 | 0.00 |
| N-methyltransferase activity | 0.01 | 0.01 | 0.00 | 0.00 | | WRC | 0.94 | 0.00 | 0.94 | 0.00 |
| DNA methylation | 0.01 | 0.01 | 0.00 | 0.00 | | Lipoxygenase | 0.94 | 0.94 | 0.00 | 0.00 |
| proton-transporting two-sector ATPase complex, proton-transporting domain | 0.01 | 0.01 | 0.00 | 0.00 | | galactosyl transferase GMA12/MNN10 family | 0.94 | 0.00 | 0.00 | 0.94 |
| WASH complex | 0.01 | 0.01 | 0.00 | 0.00 | | Glycosyl hydrolase family 10 | 0.93 | 0.93 | 0.00 | 0.00 |
| multicellular organism development | 0.01 | 0.01 | 0.00 | 0.00 | | bZIP transcription factor | 0.93 | 0.00 | 0.93 | 0.00 |
| | | | | | | FAD dependent oxidoreductase | 0.93 | 0.00 | 0.00 | 0.93 |
| | | | | | | Lectin C-type domain | 0.93 | 0.01 | 0.92 | 0.00 |
| | | | | | | Domain of unknown function (DUF4149) | 0.93 | 0.00 | 0.00 | 0.93 |
| | | | | | | X-domain of DnaJ-containing | 0.93 | 0.00 | 0.93 | 0.00 |
| | | | | | | Ribosomal protein L33 | 0.93 | 0.00 | 0.93 | 0.00 |
| | | | | | | Plant calmodulin-binding domain | 0.93 | 0.00 | 0.93 | 0.00 |
| | | | | | | Plastocyanin-like domain | 0.93 | 0.93 | 0.00 | 0.00 |
| | | | | | | Gelsolin repeat | 0.93 | 0.93 | 0.00 | 0.00 |
| | | | | | | Lung seven transmembrane receptor | 0.93 | 0.00 | 0.00 | 0.93 |
| | | | | | | Voltage-dependent anion channel | 0.93 | 0.93 | 0.00 | 0.00 |
| | | | | | | Calmodulin binding protein-like | 0.92 | 0.00 | 0.92 | 0.00 |
| | | | | | | Pirin | 0.92 | 0.00 | 0.92 | 0.00 |
| | | | | | | Domain of unknown function (DUF296) | 0.92 | 0.92 | 0.00 | 0.00 |
| | | | | | | Ferritin-like domain | 0.92 | 0.02 | 0.00 | 0.90 |
| | | | | | | Aromatic amino acid lyase | 0.92 | 0.00 | 0.00 | 0.92 |
| | | | | | | TCP family transcription factor | 0.91 | 0.00 | 0.00 | 0.91 |
| | | | | | | Zinc-binding dehydrogenase | 0.91 | 0.91 | 0.00 | 0.00 |
| | | | | | | 3-oxo-5-alpha-steroid 4-dehydrogenase | 0.91 | 0.00 | 0.91 | 0.00 |
| | | | | | | Tetraspanin family | 0.91 | 0.91 | 0.00 | 0.00 |
| | | | | | | Elongation factor Tu domain 2 | 0.91 | 0.00 | 0.00 | 0.91 |
| | | | | | | Mitochondrial branched-chain alpha-ketoacid dehydrogenase kinase | 0.90 | 0.00 | 0.00 | 0.90 |
| | | | | | | Glycine cleavage H-protein | 0.90 | 0.00 | 0.90 | 0.00 |
| | | | | | | Putative adipose-regulatory protein (Seipin) | 0.90 | 0.00 | 0.00 | 0.90 |
| | | | | | | Alba | 0.89 | 0.00 | 0.00 | 0.89 |

| GO | Total | Shared | Angio | Fern | | Domain | Total | Shared | Angio | Fern |
|---|---|---|---|---|---|---|---|---|---|---|
| | | | | | | Phosphoglucomutase/ phosphomannomutase, alpha/beta/alpha domain I | 0.89 | 0.00 | 0.00 | 0.89 |
| | | | | | | Thioesterase superfamily | 0.89 | 0.89 | 0.00 | 0.00 |
| | | | | | | GATA zinc finger | 0.89 | 0.89 | 0.00 | 0.00 |
| | | | | | | Cyclin, N-terminal domain | 0.89 | 0.89 | 0.00 | 0.00 |
| | | | | | | ACT domain | 0.89 | 0.89 | 0.00 | 0.00 |
| | | | | | | B3 DNA binding domain | 0.89 | 0.00 | 0.89 | 0.00 |
| | | | | | | E2F transcription factor CC-MB domain | 0.89 | 0.00 | 0.00 | 0.89 |
| | | | | | | RING-variant domain | 0.88 | 0.88 | 0.00 | 0.00 |
| | | | | | | Elongation factor Tu C-terminal domain | 0.88 | 0.00 | 0.00 | 0.88 |
| | | | | | | Galactose oxidase, central domain | 0.88 | 0.00 | 0.88 | 0.00 |
| | | | | | | Cathepsin propeptide inhibitor domain (I29) | 0.88 | 0.88 | 0.00 | 0.00 |
| | | | | | | HMG (high mobility group) box | 0.88 | 0.00 | 0.88 | 0.00 |
| | | | | | | BURP domain | 0.88 | 0.88 | 0.00 | 0.00 |
| | | | | | | Protein of unknown function, DUF547 | 0.88 | 0.00 | 0.88 | 0.00 |
| | | | | | | Phospholipase D Active site motif | 0.88 | 0.88 | 0.00 | 0.00 |
| | | | | | | Glycosyltransferase family 20 | 0.88 | 0.00 | 0.88 | 0.00 |
| | | | | | | Elongation factor Tu GTP binding domain | 0.88 | 0.00 | 0.00 | 0.88 |
| | | | | | | Peptidase family M28 | 0.88 | 0.00 | 0.00 | 0.88 |
| | | | | | | Golgi-body localisation protein domain | 0.88 | 0.00 | 0.88 | 0.00 |
| | | | | | | Cytidine and deoxycytidylate deaminase zinc-binding region | 0.87 | 0.00 | 0.87 | 0.00 |
| | | | | | | HSF-type DNA-binding | 0.87 | 0.87 | 0.00 | 0.00 |
| | | | | | | GlcNAc-PI de-N-acetylase | 0.87 | 0.00 | 0.87 | 0.00 |
| | | | | | | Elongation factor P, C-terminal | 0.87 | 0.00 | 0.87 | 0.00 |
| | | | | | | bHLH-MYC and R2R3-MYB transcription factors N-terminal | 0.87 | 0.31 | 0.56 | 0.00 |
| | | | | | | Pyruvate kinase, alpha/beta domain | 0.87 | 0.00 | 0.00 | 0.87 |
| | | | | | | EXS family | 0.87 | 0.87 | 0.00 | 0.00 |
| | | | | | | Domain of unknown function | 0.86 | 0.00 | 0.86 | 0.00 |
| | | | | | | Iron/manganese superoxide dismutases, alpha-hairpin domain | 0.86 | 0.00 | 0.86 | 0.00 |
| | | | | | | Hsp20/alpha crystallin family | 0.86 | 0.00 | 0.86 | 0.00 |
| | | | | | | Ulp1 protease family, C-terminal catalytic domain | 0.86 | 0.00 | 0.00 | 0.86 |
| | | | | | | D-isomer specific 2-hydroxyacid dehydrogenase, catalytic domain | 0.86 | 0.00 | 0.00 | 0.86 |
| | | | | | | Common central domain of tyrosinase | 0.86 | 0.00 | 0.00 | 0.86 |
| | | | | | | Glu/Leu/Phe/Val dehydrogenase, dimerisation domain | 0.86 | 0.00 | 0.00 | 0.86 |
| | | | | | | Small nuclear RNA activating complex (SNAPc), subunit SNAP43 | 0.86 | 0.00 | 0.86 | 0.00 |
| | | | | | | Cadherin-like beta sandwich domain | 0.86 | 0.00 | 0.00 | 0.86 |
| | | | | | | 2,3-bisphosphoglycerate-independent phosphoglycerate mutase | 0.86 | 0.00 | 0.86 | 0.00 |
| | | | | | | SRF-type transcription factor (DNA-binding and dimerisation domain) | 0.85 | 0.85 | 0.00 | 0.00 |

| GO | Total | Shared | Angio | Fern | | Domain | Total | Shared | Angio | Fern |
|---|---|---|---|---|---|---|---|---|---|---|
| | | | | | | Phosphoglucomutase/ phosphomannomutase, alpha/beta/alpha domain II | 0.85 | 0.00 | 0.00 | 0.85 |
| | | | | | | Phloem protein 2 | 0.85 | 0.00 | 0.85 | 0.00 |
| | | | | | | Nodulin-like | 0.85 | 0.85 | 0.00 | 0.00 |
| | | | | | | Cytokinin dehydrogenase 1, FAD and cytokinin binding | 0.84 | 0.00 | 0.84 | 0.00 |
| | | | | | | Elongation factor P (EF-P) OB domain | 0.84 | 0.00 | 0.84 | 0.00 |
| | | | | | | Glycosyltransferase family 92 | 0.84 | 0.84 | 0.00 | 0.00 |
| | | | | | | NADPH-dependent FMN reductase | 0.83 | 0.00 | 0.00 | 0.83 |
| | | | | | | Peptidase family M1 domain | 0.83 | 0.00 | 0.83 | 0.00 |
| | | | | | | Leukotriene A4 hydrolase, C-terminal | 0.83 | 0.00 | 0.83 | 0.00 |
| | | | | | | Pumilio-family RNA binding repeat | 0.83 | 0.00 | 0.00 | 0.83 |
| | | | | | | Helicase associated domain (HA2) | 0.83 | 0.00 | 0.83 | 0.00 |
| | | | | | | Glutamate /Leucine/ Phenylalanine/ Valine dehydrogenase | 0.83 | 0.00 | 0.00 | 0.83 |
| | | | | | | Glycosyl hydrolases family 38 N-terminal domain | 0.83 | 0.00 | 0.83 | 0.00 |
| | | | | | | BPG-independent PGAM N-terminus (iPGM_N) | 0.82 | 0.00 | 0.00 | 0.82 |
| | | | | | | Phosphoribosyl transferase domain | 0.82 | 0.00 | 0.82 | 0.00 |
| | | | | | | Alpha mannosidase middle domain | 0.82 | 0.00 | 0.82 | 0.00 |
| | | | | | | Alcohol dehydrogenase GroES-like domain | 0.82 | 0.82 | 0.00 | 0.00 |
| | | | | | | Protein of unknown function (DUF1218) | 0.81 | 0.00 | 0.81 | 0.00 |
| | | | | | | Purple acid Phosphatase, N-terminal domain | 0.81 | 0.81 | 0.00 | 0.00 |
| | | | | | | SBP domain | 0.81 | 0.00 | 0.80 | 0.00 |
| | | | | | | Strictosidine synthase | 0.81 | 0.81 | 0.00 | 0.00 |
| | | | | | | Dpy-30 motif | 0.81 | 0.00 | 0.00 | 0.81 |
| | | | | | | Legume lectin domain | 0.80 | 0.80 | 0.00 | 0.00 |
| | | | | | | C-terminus of AA_permease | 0.80 | 0.80 | 0.00 | 0.00 |
| | | | | | | Phospholipase D C terminal | 0.80 | 0.80 | 0.00 | 0.00 |
| | | | | | | Uncharacterised protein family (UPF0121) | 0.80 | 0.00 | 0.80 | 0.00 |
| | | | | | | Pre-SET motif | 0.80 | 0.00 | 0.80 | 0.00 |
| | | | | | | Subtilase family | 0.80 | 0.80 | 0.00 | 0.00 |
| | | | | | | TPL-binding domain in jasmonate signalling | 0.79 | 0.00 | 0.79 | 0.00 |
| | | | | | | Protein of unknown function (DUF498/DUF598) | 0.79 | 0.00 | 0.00 | 0.79 |
| | | | | | | GDP-fucose protein O-fucosyltransferase | 0.79 | 0.79 | 0.00 | 0.00 |
| | | | | | | Phospholipase/ Carboxylesterase | 0.79 | 0.00 | 0.79 | 0.00 |
| | | | | | | MYB-CC type transfactor, LHEQLE motif | 0.78 | 0.78 | 0.00 | 0.00 |
| | | | | | | Rhamnogalacturonate lyase family | 0.78 | 0.00 | 0.78 | 0.00 |
| | | | | | | Leucine-zipper of ternary complex factor MIP1 | 0.78 | 0.00 | 0.78 | 0.00 |
| | | | | | | C-terminal domain of alpha-glycerophosphate oxidase | 0.77 | 0.00 | 0.00 | 0.77 |
| | | | | | | NEMP family | 0.77 | 0.00 | 0.77 | 0.00 |
| | | | | | | Protein of unknown function (DUF3675) | 0.77 | 0.77 | 0.00 | 0.00 |
| | | | | | | FAD-binding domain | 0.77 | 0.77 | 0.00 | 0.00 |
| | | | | | | His Kinase A (phospho-acceptor) domain | 0.76 | 0.00 | 0.00 | 0.76 |
| | | | | | | Polyphenol oxidase middle domain | 0.76 | 0.00 | 0.00 | 0.76 |

| GO | Total | Shared | Angio | Fern | | Domain | Total | Shared | Angio | Fern |
|---|---|---|---|---|---|---|---|---|---|---|
| | | | | | | Cellulase (glycosyl hydrolase family 5) | 0.76 | 0.76 | 0.00 | 0.00 |
| | | | | | | Glycosyl hydrolases family 38 C-terminal domain | 0.76 | 0.00 | 0.76 | 0.00 |
| | | | | | | Exonuclease | 0.76 | 0.76 | 0.00 | 0.00 |
| | | | | | | DnaJ domain | 0.75 | 0.00 | 0.75 | 0.01 |
| | | | | | | 3-Oxoacyl-[acyl-carrier-protein (ACP)] synthase III C terminal | 0.75 | 0.75 | 0.00 | 0.00 |
| | | | | | | PMR5 N terminal Domain | 0.75 | 0.75 | 0.00 | 0.00 |
| | | | | | | Xyloglucan endo-transglycosylase (XET) C-terminus | 0.75 | 0.75 | 0.00 | 0.00 |
| | | | | | | Cyclin, C-terminal domain | 0.74 | 0.74 | 0.00 | 0.00 |
| | | | | | | NADH-ubiquinone reductase complex 1 MLRQ subunit | 0.74 | 0.00 | 0.74 | 0.00 |
| | | | | | | MASE1 | 0.74 | 0.00 | 0.00 | 0.74 |
| | | | | | | Elongation factor P (EF-P) KOW-like domain | 0.74 | 0.00 | 0.74 | 0.00 |
| | | | | | | BAR domain of APPL family | 0.74 | 0.00 | 0.74 | 0.00 |
| | | | | | | Polysaccharide lyase family 4, domain II | 0.74 | 0.00 | 0.74 | 0.00 |
| | | | | | | Adenylate kinase | 0.73 | 0.00 | 0.00 | 0.73 |
| | | | | | | Ferric reductase NAD binding domain | 0.72 | 0.72 | 0.00 | 0.00 |
| | | | | | | QLQ | 0.72 | 0.00 | 0.72 | 0.00 |
| | | | | | | Neprosin activation peptide | 0.72 | 0.72 | 0.00 | 0.00 |
| | | | | | | SPX domain | 0.72 | 0.72 | 0.00 | 0.00 |
| | | | | | | Protein of unknown function (DUF1298) | 0.72 | 0.00 | 0.72 | 0.00 |
| | | | | | | Rhomboid family | 0.72 | 0.00 | 0.72 | 0.00 |
| | | | | | | Peroxisomal membrane anchor protein (Pex14p) conserved region | 0.71 | 0.00 | 0.00 | 0.71 |
| | | | | | | Plant PDR ABC transporter associated | 0.71 | 0.71 | 0.00 | 0.00 |
| | | | | | | Conserved region of unknown function on GLTSCR protein | 0.71 | 0.00 | 0.00 | 0.71 |
| | | | | | | FAD binding domain | 0.70 | 0.00 | 0.68 | 0.01 |
| | | | | | | Copine | 0.70 | 0.00 | 0.00 | 0.70 |
| | | | | | | Protein of unknown function, DUF594 | 0.70 | 0.00 | 0.70 | 0.00 |
| | | | | | | Methyltransferase domain | 0.70 | 0.00 | 0.01 | 0.68 |
| | | | | | | Acyltransferase | 0.70 | 0.70 | 0.00 | 0.00 |
| | | | | | | Polysaccharide lyase family 4, domain III | 0.70 | 0.00 | 0.70 | 0.00 |
| | | | | | | Domain of unknown function (DUF4782) | 0.70 | 0.00 | 0.70 | 0.00 |
| | | | | | | Phosphoglucomutase/ phosphomannomutase, alpha/beta/alpha domain III | 0.69 | 0.00 | 0.00 | 0.69 |
| | | | | | | Chibby family | 0.69 | 0.00 | 0.00 | 0.69 |
| | | | | | | Zinc-finger of C2H2 type | 0.69 | 0.00 | 0.00 | 0.69 |
| | | | | | | Synaptotagmin-like mitochondrial-lipid-binding domain | 0.69 | 0.00 | 0.69 | 0.00 |
| | | | | | | C-terminal associated domain of TOPRIM | 0.68 | 0.00 | 0.68 | 0.00 |
| | | | | | | Oligonucleotide/ oligosaccharide- binding (OB)-fold | 0.68 | 0.00 | 0.68 | 0.00 |
| | | | | | | PHD - plant homeodomain finger protein | 0.68 | 0.00 | 0.68 | 0.00 |
| | | | | | | Fibronectin type III-like domain | 0.68 | 0.68 | 0.00 | 0.00 |
| | | | | | | DDT domain | 0.67 | 0.00 | 0.67 | 0.00 |

| GO | Total | Shared | Angio | Fern | | Domain | Total | Shared | Angio | Fern |
|---|---|---|---|---|---|---|---|---|---|---|
| | | | | | | Ferric reductase like transmembrane component | 0.67 | 0.67 | 0.00 | 0.00 |
| | | | | | | Alternative splicing regulator | 0.66 | 0.00 | 0.66 | 0.00 |
| | | | | | | MIZ/SP-RING zinc finger | 0.66 | 0.00 | 0.66 | 0.00 |
| | | | | | | Diacylglycerol acyltransferase | 0.65 | 0.00 | 0.65 | 0.00 |
| | | | | | | Ligand-gated ion channel | 0.65 | 0.65 | 0.00 | 0.00 |
| | | | | | | E2F/DP family winged-helix DNA-binding domain | 0.65 | 0.00 | 0.00 | 0.65 |
| | | | | | | Bacterial extracellular solute-binding proteins, family 3 | 0.65 | 0.65 | 0.00 | 0.00 |
| | | | | | | Nuclease-related domain | 0.65 | 0.00 | 0.00 | 0.65 |
| | | | | | | DNA gyrase B | 0.64 | 0.00 | 0.64 | 0.00 |
| | | | | | | RNA polymerase II-binding domain. | 0.64 | 0.00 | 0.64 | 0.00 |
| | | | | | | Fusaric acid resistance protein-like | 0.64 | 0.00 | 0.00 | 0.63 |
| | | | | | | CRAL/TRIO, N-terminal domain | 0.64 | 0.00 | 0.00 | 0.64 |
| | | | | | | Probable zinc-ribbon domain | 0.63 | 0.00 | 0.63 | 0.00 |
| | | | | | | PB1 domain | 0.62 | 0.00 | 0.62 | 0.00 |
| | | | | | | Endonuclease/ Exonuclease/ phosphatase family | 0.62 | 0.00 | 0.49 | 0.12 |
| | | | | | | Regulated-SNARE-like domain | 0.62 | 0.00 | 0.00 | 0.62 |
| | | | | | | D-mannose binding lectin | 0.62 | 0.12 | 0.00 | 0.50 |
| | | | | | | Iron/zinc purple acid phosphatase-like protein C | 0.62 | 0.62 | 0.00 | 0.00 |
| | | | | | | Pyruvate phosphate dikinase, PEP/pyruvate binding domain | 0.61 | 0.00 | 0.61 | 0.00 |
| | | | | | | Ankyrin repeats (many copies) | 0.61 | 0.00 | 0.47 | 0.15 |
| | | | | | | Nucleic acid binding protein NABP | 0.61 | 0.00 | 0.00 | 0.61 |
| | | | | | | Vacuolar sorting protein 9 (VPS9) domain | 0.61 | 0.00 | 0.00 | 0.61 |
| | | | | | | Domain of unknown function (DUF4408) | 0.61 | 0.00 | 0.00 | 0.61 |
| | | | | | | Cotton fibre expressed protein | 0.61 | 0.00 | 0.00 | 0.61 |
| | | | | | | Protein of unknown function (DUF3455) | 0.60 | 0.00 | 0.00 | 0.60 |
| | | | | | | Protein of unknown function (DUF630) | 0.59 | 0.59 | 0.00 | 0.00 |
| | | | | | | Ubiquitin-binding WIYLD domain | 0.59 | 0.00 | 0.59 | 0.00 |
| | | | | | | NB-ARC domain | 0.58 | 0.00 | 0.00 | 0.58 |
| | | | | | | Patched family | 0.58 | 0.00 | 0.00 | 0.58 |
| | | | | | | Toprim domain | 0.58 | 0.00 | 0.58 | 0.00 |
| | | | | | | Chaperone for wingless signalling and trafficking of LDL receptor | 0.58 | 0.58 | 0.00 | 0.00 |
| | | | | | | Domain of unknown function (DUF4218) | 0.58 | 0.00 | 0.00 | 0.58 |
| | | | | | | Wax ester synthase-like Acyl-CoA acyltransferase domain | 0.57 | 0.00 | 0.57 | 0.00 |
| | | | | | | Transposase family tnp2 | 0.57 | 0.00 | 0.00 | 0.57 |
| | | | | | | Pathogenesis-related protein Bet v I family | 0.57 | 0.00 | 0.00 | 0.57 |
| | | | | | | Domain of unknown function (DUF4094) | 0.57 | 0.57 | 0.00 | 0.00 |
| | | | | | | Lipase (class 3) | 0.57 | 0.56 | 0.00 | 0.01 |
| | | | | | | Cation transporter/ATPase, N-terminus | 0.55 | 0.55 | 0.00 | 0.00 |
| | | | | | | Alpha/beta hydrolase family | 0.55 | 0.00 | 0.55 | 0.00 |
| | | | | | | Homeobox domain | 0.55 | 0.55 | 0.00 | 0.00 |

| GO | Total | Shared | Angio | Fern | | Domain | Total | Shared | Angio | Fern |
|---|---|---|---|---|---|---|---|---|---|---|
| | | | | | | WSTF, HB1, Itc1p, MBD9 motif 1 | 0.55 | 0.00 | 0.55 | 0.00 |
| | | | | | | Protein of unknown function (DUF_B2219) | 0.54 | 0.00 | 0.00 | 0.54 |
| | | | | | | Phosphoglucomutase/ phosphomannomutase, C-terminal domain | 0.53 | 0.00 | 0.00 | 0.53 |
| | | | | | | Zinc knuckle | 0.53 | 0.25 | 0.13 | 0.14 |
| | | | | | | Dicer dimerisation domain | 0.52 | 0.00 | 0.52 | 0.00 |
| | | | | | | GUCT (NUC152) domain | 0.52 | 0.00 | 0.52 | 0.00 |
| | | | | | | G-box binding protein MFMR | 0.52 | 0.00 | 0.52 | 0.00 |
| | | | | | | Receptor family ligand binding region | 0.52 | 0.52 | 0.00 | 0.00 |
| | | | | | | K-box region | 0.52 | 0.52 | 0.00 | 0.00 |
| | | | | | | Respiratory burst NADPH oxidase | 0.51 | 0.51 | 0.00 | 0.00 |
| | | | | | | Leucine Rich repeats (2 copies) | 0.51 | 0.35 | 0.16 | 0.00 |
| | | | | | | TPR repeat | 0.50 | 0.00 | 0.00 | 0.50 |
| | | | | | | Glycosyltransferase like family 2 | 0.50 | 0.00 | 0.50 | 0.00 |
| | | | | | | Peptidase inhibitor I9 | 0.49 | 0.49 | 0.00 | 0.00 |
| | | | | | | BTB/POZ domain | 0.48 | 0.40 | 0.00 | 0.08 |
| | | | | | | F-box-like | 0.47 | 0.00 | 0.07 | 0.40 |
| | | | | | | BED zinc finger | 0.46 | 0.00 | 0.05 | 0.41 |
| | | | | | | Disordered region downstream of MFMR | 0.46 | 0.00 | 0.46 | 0.00 |
| | | | | | | Plant protein of unknown function (DUF641) | 0.45 | 0.00 | 0.00 | 0.45 |
| | | | | | | Glyoxalase/Bleomycin resistance protein/Dioxygenase superfamily | 0.44 | 0.00 | 0.00 | 0.44 |
| | | | | | | Glycosyl transferase family group 2 | 0.44 | 0.00 | 0.44 | 0.00 |
| | | | | | | Myosin N-terminal SH3-like domain | 0.44 | 0.00 | 0.44 | 0.00 |
| | | | | | | Protein of unknown function (DUF4005) | 0.44 | 0.44 | 0.00 | 0.00 |
| | | | | | | NHL repeat | 0.43 | 0.00 | 0.43 | 0.00 |
| | | | | | | PDDEXK-like family of unknown function | 0.43 | 0.00 | 0.00 | 0.43 |
| | | | | | | Sulfotransferase domain | 0.43 | 0.43 | 0.00 | 0.00 |
| | | | | | | Bromodomain | 0.42 | 0.00 | 0.42 | 0.00 |
| | | | | | | Villin headpiece domain | 0.42 | 0.42 | 0.00 | 0.00 |
| | | | | | | DIL domain | 0.42 | 0.00 | 0.42 | 0.00 |
| | | | | | | Protein of unknown function (DUF688) | 0.42 | 0.00 | 0.00 | 0.42 |
| | | | | | | NAD dependent epimerase/dehydratase family | 0.42 | 0.30 | 0.11 | 0.00 |
| | | | | | | FBD | 0.39 | 0.00 | 0.39 | 0.00 |
| | | | | | | PLAT/LH2 domain | 0.39 | 0.39 | 0.00 | 0.00 |
| | | | | | | ShK domain-like | 0.39 | 0.39 | 0.00 | 0.00 |
| | | | | | | Sterol-sensing domain of SREBP cleavage-activation | 0.39 | 0.00 | 0.00 | 0.39 |
| | | | | | | UBX domain | 0.38 | 0.00 | 0.38 | 0.00 |
| | | | | | | Pyridine nucleotide-disulphide oxidoreductase | 0.38 | 0.38 | 0.00 | 0.00 |
| | | | | | | RNA pol II promoter Fmp27 protein domain | 0.38 | 0.00 | 0.38 | 0.00 |
| | | | | | | Microtubule binding | 0.36 | 0.00 | 0.36 | 0.00 |
| | | | | | | zinc-binding in reverse transcriptase | 0.35 | 0.00 | 0.00 | 0.35 |
| | | | | | | C2 domain of PTEN tumour-suppressor protein | 0.35 | 0.00 | 0.35 | 0.00 |

| GO | Total | Shared | Angio | Fern | | Domain | Total | Shared | Angio | Fern |
|---|---|---|---|---|---|---|---|---|---|---|
| | | | | | | Armadillo/beta-catenin-like repeat | 0.34 | 0.00 | 0.34 | 0.00 |
| | | | | | | ABC-transporter extracellular N-terminal | 0.33 | 0.33 | 0.00 | 0.00 |
| | | | | | | Domain of unknown function (DUF4283) | 0.32 | 0.00 | 0.00 | 0.32 |
| | | | | | | EF-hand domain | 0.32 | 0.08 | 0.00 | 0.24 |
| | | | | | | PA domain | 0.31 | 0.31 | 0.00 | 0.00 |
| | | | | | | double strand RNA binding domain from DEAD END PROTEIN 1 | 0.31 | 0.00 | 0.31 | 0.00 |
| | | | | | | PHD-finger | 0.31 | 0.00 | 0.31 | 0.00 |
| | | | | | | Ankyrin repeat | 0.31 | 0.00 | 0.31 | 0.00 |
| | | | | | | impB/mucB/samB family | 0.30 | 0.00 | 0.30 | 0.00 |
| | | | | | | Zinc-binding RING-finger | 0.29 | 0.00 | 0.29 | 0.00 |
| | | | | | | Granulin | 0.29 | 0.29 | 0.00 | 0.00 |
| | | | | | | Fusaric acid resistance protein family | 0.29 | 0.00 | 0.00 | 0.29 |
| | | | | | | Calponin homology (CH) domain | 0.29 | 0.00 | 0.29 | 0.00 |
| | | | | | | Plant invertase/pectin methylesterase inhibitor | 0.27 | 0.27 | 0.00 | 0.00 |
| | | | | | | LysM domain | 0.27 | 0.00 | 0.27 | 0.00 |
| | | | | | | EF hand | 0.26 | 0.12 | 0.00 | 0.15 |
| | | | | | | ARID/BRIGHT DNA binding domain | 0.25 | 0.00 | 0.25 | 0.00 |
| | | | | | | AAA domain | 0.25 | 0.00 | 0.03 | 0.23 |
| | | | | | | Mitochondrial ATPase inhibitor, IATP | 0.25 | 0.00 | 0.00 | 0.25 |
| | | | | | | impB/mucB/samB family C-terminal domain | 0.24 | 0.00 | 0.24 | 0.00 |
| | | | | | | Fibronectin type III domain | 0.24 | 0.00 | 0.24 | 0.00 |
| | | | | | | Double-stranded RNA binding motif | 0.24 | 0.00 | 0.24 | 0.00 |
| | | | | | | CCT motif | 0.24 | 0.24 | 0.00 | 0.00 |
| | | | | | | Carbohydrate binding domain | 0.24 | 0.24 | 0.00 | 0.00 |
| | | | | | | BRCT domain, a BRCA1 C-terminus domain | 0.23 | 0.00 | 0.23 | 0.00 |
| | | | | | | Conserved region of Rad21 / Rec8 like protein | 0.23 | 0.00 | 0.00 | 0.23 |
| | | | | | | GAG-pre-integrase domain | 0.22 | 0.00 | 0.00 | 0.22 |
| | | | | | | vWA found in TerF C terminus | 0.22 | 0.00 | 0.00 | 0.22 |
| | | | | | | Carbohydrate binding domain CBM49 | 0.21 | 0.21 | 0.00 | 0.00 |
| | | | | | | Staphylococcal nuclease homologue | 0.20 | 0.00 | 0.20 | 0.00 |
| | | | | | | Zinc finger, C2H2 type | 0.20 | 0.01 | 0.00 | 0.20 |
| | | | | | | Major Facilitator Superfamily | 0.20 | 0.19 | 0.01 | 0.00 |
| | | | | | | Right handed beta helix region | 0.20 | 0.00 | 0.00 | 0.20 |
| | | | | | | Type III restriction enzyme, res subunit | 0.16 | 0.00 | 0.16 | 0.00 |
| | | | | | | PLD-like domain | 0.16 | 0.16 | 0.00 | 0.00 |
| | | | | | | Dimerisation domain of Ca+-activated chloride-channel, anoctamin | 0.16 | 0.00 | 0.16 | 0.00 |
| | | | | | | SPFH domain / Band 7 family | 0.16 | 0.16 | 0.00 | 0.00 |
| | | | | | | Plant zinc cluster domain | 0.15 | 0.15 | 0.00 | 0.00 |
| | | | | | | Remorin, N-terminal region | 0.14 | 0.14 | 0.00 | 0.00 |
| | | | | | | Root cap | 0.14 | 0.00 | 0.00 | 0.14 |
| | | | | | | RING/Ubox like zinc-binding domain | 0.14 | 0.00 | 0.14 | 0.00 |

| GO | Total | Shared | Angio | Fern | | Domain | Total | Shared | Angio | Fern |
|---|---|---|---|---|---|---|---|---|---|---|
| | | | | | | Reverse transcriptase (RNA-dependent DNA polymerase) | 0.12 | 0.01 | 0.07 | 0.05 |
| | | | | | | Folate receptor family | 0.11 | 0.00 | 0.11 | 0.00 |
| | | | | | | PQQ enzyme repeat | 0.10 | 0.00 | 0.00 | 0.10 |
| | | | | | | Chlamydia polymorphic membrane protein (Chlamydia_PMP) repeat | 0.10 | 0.00 | 0.00 | 0.10 |
| | | | | | | alpha/beta hydrolase fold | 0.10 | 0.00 | 0.10 | 0.00 |
| | | | | | | Protein of unknown function (DUF 659) | 0.10 | 0.02 | 0.00 | 0.07 |
| | | | | | | Di-glucose binding within endoplasmic reticulum | 0.09 | 0.00 | 0.09 | 0.00 |
| | | | | | | Ribosomal L40e family | 0.09 | 0.09 | 0.00 | 0.00 |
| | | | | | | Carbohydrate-binding protein of the ER | 0.09 | 0.08 | 0.00 | 0.00 |
| | | | | | | Aminotransferase class I and II | 0.08 | 0.00 | 0.00 | 0.08 |
| | | | | | | Synaptobrevin | 0.08 | 0.00 | 0.00 | 0.08 |
| | | | | | | Aminotransferase class-V | 0.08 | 0.00 | 0.08 | 0.00 |
| | | | | | | Rtr1/RPAP2 family | 0.08 | 0.00 | 0.00 | 0.08 |
| | | | | | | Bacterial Ig-like domain (group 2) | 0.08 | 0.00 | 0.08 | 0.00 |
| | | | | | | Ribosomal protein S27a | 0.07 | 0.07 | 0.00 | 0.00 |
| | | | | | | IMS family HHH motif | 0.07 | 0.00 | 0.07 | 0.00 |
| | | | | | | AT hook motif | 0.07 | 0.03 | 0.00 | 0.04 |
| | | | | | | Senescence-associated protein | 0.07 | 0.00 | 0.00 | 0.07 |
| | | | | | | Der1-like family | 0.07 | 0.00 | 0.07 | 0.00 |
| | | | | | | Glyceraldehyde 3-phosphate dehydrogenase, C-terminal domain | 0.06 | 0.00 | 0.06 | 0.00 |
| | | | | | | S-locus glycoprotein domain | 0.06 | 0.06 | 0.00 | 0.00 |
| | | | | | | Glycosyl transferase family 2 | 0.06 | 0.00 | 0.06 | 0.00 |
| | | | | | | Pectate lyase superfamily protein | 0.06 | 0.06 | 0.00 | 0.00 |
| | | | | | | Zinc-finger double-stranded RNA-binding | 0.06 | 0.00 | 0.00 | 0.06 |
| | | | | | | Retroviral aspartyl protease | 0.05 | 0.00 | 0.05 | 0.00 |
| | | | | | | Sulfotransferase family | 0.05 | 0.05 | 0.00 | 0.00 |
| | | | | | | Haspin like kinase domain | 0.05 | 0.05 | 0.00 | 0.00 |
| | | | | | | RING-type zinc-finger | 0.05 | 0.00 | 0.05 | 0.00 |
| | | | | | | Ligated ion channel L-glutamate- and glycine-binding site | 0.05 | 0.05 | 0.00 | 0.00 |
| | | | | | | Wall-associated receptor kinase C-terminal | 0.05 | 0.05 | 0.00 | 0.00 |
| | | | | | | TAP-like protein | 0.05 | 0.00 | 0.05 | 0.00 |
| | | | | | | PAN-like domain | 0.05 | 0.05 | 0.00 | 0.00 |
| | | | | | | Transposase-associated domain | 0.05 | 0.00 | 0.00 | 0.05 |
| | | | | | | Endomembrane protein 70 | 0.05 | 0.00 | 0.00 | 0.05 |
| | | | | | | tify domain | 0.04 | 0.00 | 0.00 | 0.04 |
| | | | | | | Domain of unknown function (DUF3598) | 0.04 | 0.00 | 0.00 | 0.04 |
| | | | | | | gag-polypeptide of LTR copia-type | 0.04 | 0.01 | 0.00 | 0.03 |
| | | | | | | Divergent CCT motif | 0.04 | 0.00 | 0.00 | 0.04 |
| | | | | | | NAD(P)-binding Rossmann-like domain | 0.04 | 0.02 | 0.01 | 0.01 |
| | | | | | | Integrase core domain | 0.04 | 0.01 | 0.02 | 0.01 |
| | | | | | | Auxin canalisation | 0.04 | 0.00 | 0.04 | 0.00 |

| GO | Total | Shared | Angio | Fern | | Domain | Total | Shared | Angio | Fern |
|---|---|---|---|---|---|---|---|---|---|---|
| | | | | | | Chalcone and stilbene synthases, C-terminal domain | 0.04 | 0.04 | 0.00 | 0.00 |
| | | | | | | SAWADEE domain | 0.04 | 0.00 | 0.04 | 0.00 |
| | | | | | | Wall-associated receptor kinase galacturonan-binding | 0.03 | 0.03 | 0.00 | 0.00 |
| | | | | | | AAA domain, putative AbiEii toxin, Type IV TA system | 0.03 | 0.00 | 0.03 | 0.00 |
| | | | | | | Helix-turn-helix domain | 0.03 | 0.00 | 0.03 | 0.00 |
| | | | | | | Oxidoreductase NAD-binding domain | 0.03 | 0.00 | 0.03 | 0.00 |
| | | | | | | 2Fe-2S iron-sulfur cluster binding domain | 0.03 | 0.00 | 0.03 | 0.00 |
| | | | | | | Cholesterol-capturing domain | 0.03 | 0.00 | 0.03 | 0.00 |
| | | | | | | Terpene synthase family, metal binding domain | 0.03 | 0.00 | 0.00 | 0.03 |
| | | | | | | REJ domain | 0.03 | 0.00 | 0.00 | 0.03 |
| | | | | | | Lysine methyltransferase | 0.03 | 0.00 | 0.00 | 0.03 |
| | | | | | | F-box associated domain | 0.03 | 0.00 | 0.03 | 0.00 |
| | | | | | | Malic enzyme, NAD binding domain | 0.03 | 0.00 | 0.03 | 0.00 |
| | | | | | | Piwi domain | 0.03 | 0.00 | 0.00 | 0.03 |
| | | | | | | PPR repeat | 0.02 | 0.00 | 0.01 | 0.02 |
| | | | | | | Cation transport ATPase (P-type) | 0.02 | 0.02 | 0.00 | 0.00 |
| | | | | | | SAP domain | 0.02 | 0.00 | 0.02 | 0.00 |
| | | | | | | BES1/BZR1 plant transcription factor, N-terminal | 0.02 | 0.02 | 0.00 | 0.00 |
| | | | | | | ELM2 domain | 0.02 | 0.00 | 0.02 | 0.00 |
| | | | | | | RecF/RecN/SMC N terminal domain | 0.02 | 0.00 | 0.00 | 0.02 |
| | | | | | | Sigma-54 interaction domain | 0.02 | 0.02 | 0.00 | 0.00 |
| | | | | | | Fringe-like | 0.02 | 0.02 | 0.00 | 0.00 |
| | | | | | | PPR repeat family | 0.02 | 0.00 | 0.01 | 0.02 |
| | | | | | | Skp1 family, dimerisation domain | 0.02 | 0.00 | 0.02 | 0.00 |
| | | | | | | PAN domain | 0.02 | 0.02 | 0.00 | 0.00 |
| | | | | | | Acetyltransferase (GNAT) family | 0.02 | 0.02 | 0.00 | 0.00 |
| | | | | | | Core histone H2A/H2B/H3/H4 | 0.02 | 0.00 | 0.02 | 0.00 |
| | | | | | | Lateral organ boundaries (LOB) domain | 0.02 | 0.00 | 0.00 | 0.02 |
| | | | | | | ATPase family associated with various cellular activities (AAA) | 0.02 | 0.01 | 0.00 | 0.01 |
| | | | | | | Poly A polymerase head domain | 0.02 | 0.00 | 0.00 | 0.02 |
| | | | | | | Probable RNA and SrmB-binding site of polymerase A | 0.02 | 0.00 | 0.00 | 0.02 |
| | | | | | | RING-H2 zinc finger domain | 0.02 | 0.02 | 0.00 | 0.00 |
| | | | | | | Ribonucleotide reductase, small chain | 0.02 | 0.00 | 0.02 | 0.00 |
| | | | | | | Mycolic acid cyclopropane synthetase | 0.02 | 0.00 | 0.00 | 0.02 |
| | | | | | | Extensin-like region | 0.02 | 0.00 | 0.02 | 0.00 |
| | | | | | | WAX2 C-terminal domain | 0.02 | 0.00 | 0.00 | 0.01 |
| | | | | | | Enolase, N-terminal domain | 0.02 | 0.00 | 0.00 | 0.02 |
| | | | | | | F-box-like domain | 0.02 | 0.00 | 0.00 | 0.02 |
| | | | | | | PEHE domain | 0.02 | 0.00 | 0.02 | 0.00 |
| | | | | | | RING-like zinc finger | 0.01 | 0.01 | 0.00 | 0.00 |
| | | | | | | Peptidase family S64 | 0.01 | 0.01 | 0.00 | 0.00 |

| GO | Total | Shared | Angio | Fern | | Domain | Total | Shared | Angio | Fern |
|---|---|---|---|---|---|---|---|---|---|---|
| | | | | | | Alanine dehydrogenase/PNT, C-terminal domain | 0.01 | 0.01 | 0.00 | 0.00 |
| | | | | | | Concanavalin A-like lectin/glucanases superfamily | 0.01 | 0.00 | 0.00 | 0.01 |
| | | | | | | 4Fe-4S binding domain | 0.01 | 0.00 | 0.00 | 0.01 |
| | | | | | | Phage integrase family | 0.01 | 0.00 | 0.00 | 0.01 |
| | | | | | | Domain found in IF2B/IF5 | 0.01 | 0.00 | 0.01 | 0.00 |
| | | | | | | TOBE domain | 0.01 | 0.00 | 0.00 | 0.01 |
| | | | | | | Binding-protein-dependent transport system inner membrane component | 0.01 | 0.00 | 0.00 | 0.01 |
| | | | | | | Substrate binding domain of ABC-type glycine betaine transport system | 0.01 | 0.00 | 0.00 | 0.01 |
| | | | | | | Fn3-like domain | 0.01 | 0.01 | 0.00 | 0.00 |
| | | | | | | Fumarylacetoacetate (FAA) hydrolase family | 0.01 | 0.00 | 0.01 | 0.00 |
| | | | | | | Cyclic nucleotide-binding domain | 0.01 | 0.00 | 0.00 | 0.01 |
| | | | | | | Ribosomal proteins 50S-L15, 50S-L18e, 60S-L27A | 0.01 | 0.00 | 0.01 | 0.00 |
| | | | | | | Sugar (and other) transporter | 0.01 | 0.00 | 0.01 | 0.00 |
| | | | | | | JmjC domain, hydroxylase | 0.01 | 0.00 | 0.01 | 0.00 |
| | | | | | | Zinc-finger domain of monoamine-oxidase A repressor R1 | 0.01 | 0.00 | 0.01 | 0.00 |
| | | | | | | Utp25, U3 small nucleolar RNA-associated SSU processome protein 25 | 0.01 | 0.00 | 0.01 | 0.00 |
| | | | | | | Type I phosphodiesterase / nucleotide pyrophosphatase | 0.01 | 0.00 | 0.00 | 0.01 |
| | | | | | | Domain of unknown function (DUF4216) | 0.01 | 0.00 | 0.00 | 0.01 |
| | | | | | | Plant transposase (Ptta/En/Spm family) | 0.01 | 0.00 | 0.00 | 0.01 |
| | | | | | | Selenoprotein SelK_SelG | 0.01 | 0.01 | 0.00 | 0.00 |
| | | | | | | G-patch domain | 0.01 | 0.01 | 0.00 | 0.00 |
| | | | | | | Protein of unknown function (DUF1296) | 0.01 | 0.00 | 0.01 | 0.00 |
| | | | | | | Class-II DAHP synthetase family | 0.01 | 0.01 | 0.00 | 0.00 |
| | | | | | | Lamin Tail Domain | 0.01 | 0.00 | 0.01 | 0.00 |
| | | | | | | CAP-Gly domain | 0.01 | 0.01 | 0.00 | 0.00 |
| | | | | | | RNA-binding, Nab2-type zinc finger | 0.01 | 0.01 | 0.00 | 0.00 |
| | | | | | | B-block binding subunit of TFIIIC | 0.01 | 0.00 | 0.01 | 0.00 |
| | | | | | | CLASP N terminal | 0.01 | 0.00 | 0.01 | 0.00 |
| | | | | | | Bacterial regulatory protein, Fis family | 0.01 | 0.01 | 0.00 | 0.00 |
| | | | | | | PAS domain | 0.01 | 0.01 | 0.00 | 0.00 |
| | | | | | | Tubulin C-terminal domain | 0.01 | 0.01 | 0.00 | 0.00 |
| | | | | | | zinc-ribbon | 0.01 | 0.01 | 0.00 | 0.00 |
| | | | | | | Microtubule associated protein (MAP65/ASE1 family) | 0.01 | 0.00 | 0.00 | 0.01 |
| | | | | | | Prolyl oligopeptidase family | 0.01 | 0.00 | 0.01 | 0.00 |
| | | | | | | Reverse transcriptase-like | 0.01 | 0.00 | 0.01 | 0.00 |
| | | | | | | RhoGAP domain | 0.01 | 0.00 | 0.01 | 0.00 |
| | | | | | | Exocyst complex component Sec10 | 0.01 | 0.00 | 0.01 | 0.00 |
| | | | | | | ThiF family | 0.01 | 0.00 | 0.01 | 0.00 |
| | | | | | | Ubiquitin-activating enzyme active site | 0.01 | 0.00 | 0.01 | 0.00 |

| GO | Total | Shared | Angio | Fern | | Domain | Total | Shared | Angio | Fern |
|---|---|---|---|---|---|---|---|---|---|---|
| | | | | | | Putative nucleotide-binding of sugar-metabolising enzyme | 0.01 | 0.00 | 0.00 | 0.01 |
| | | | | | | Fructose-bisphosphate aldolase class-II | 0.01 | 0.00 | 0.00 | 0.01 |
| | | | | | | Protein of unknown function (DUF2974) | 0.01 | 0.01 | 0.00 | 0.00 |
| | | | | | | Eukaryotic DNA topoisomerase I, catalytic core | 0.01 | 0.01 | 0.00 | 0.00 |
| | | | | | | Zinc finger, ZZ type | 0.01 | 0.01 | 0.00 | 0.00 |
| | | | | | | Peptidase family C78 | 0.01 | 0.01 | 0.00 | 0.00 |
| | | | | | | Protein of unknown function (DUF642) | 0.01 | 0.00 | 0.00 | 0.00 |
| | | | | | | DNA methylase | 0.01 | 0.01 | 0.00 | 0.00 |
| | | | | | | ATP synthase subunit C | 0.01 | 0.01 | 0.00 | 0.00 |
| | | | | | | Subunit CCDC53 of WASH complex | 0.01 | 0.01 | 0.00 | 0.00 |
| | | | | | | LURP-one-related | 0.01 | 0.01 | 0.00 | 0.00 |
| | | | | | | Seven in absentia protein family | 0.01 | 0.01 | 0.00 | 0.00 |
| | | | | | | YMGG-like Gly-zipper | 0.01 | 0.01 | 0.00 | 0.00 |
| | | | | | | Pentatricopeptide repeat domain | 0.01 | 0.00 | 0.01 | 0.00 |
| | | | | | | EGF-like domain | 0.01 | 0.01 | 0.00 | 0.00 |
| | | | | | | Ricin-type beta-trefoil lectin domain-like | 0.01 | 0.00 | 0.00 | 0.01 |
| | | | | | | F-box associated | 0.01 | 0.00 | 0.01 | 0.00 |
| | | | | | | Scavenger mRNA decapping enzyme C-term binding | 0.01 | 0.01 | 0.00 | 0.00 |
| | | | | | | Aspartyl protease | 0.01 | 0.00 | 0.00 | 0.00 |
| | | | | | | Berberine and berberine like | 0.01 | 0.00 | 0.01 | 0.00 |

**Table S2. Details of expression of genes of interest and associated orthogroups.** Relevant orthogroups/clades were identified according to previously published family members. Colour coding represents the highest relative expression in guard cell-enriched samples compared to whole leaves for angiosperm (At, *Arabidopsis thaliana*; Hv, *Hordeum vulgare* barley) and fern (Cr, *Ceratopteris richardii*; Pv, *Polypodium vulgare*) genes within each orthogroup as follows: orange – "enriched" (significantly higher expression in guard cell than leaf samples, padj ≤ 0.01), yellow – "expressed" (mean expression in guard samples ≥ 10). White represents "negligible" expression (mean expression in guard samples < 10) or the absence of a gene from this species within this orthogroup. For *P. vulgare* samples only, whole leaf vs leaf samples without abaxial epidermis (thus guard cells) removed were also included and used to separate guard cell-enriched genes with a higher level of stringency (red; "enriched both comparisons" = expression higher in guard cells than leaves and higher in whole leaves than leaves without guard cells). Guard cell expression groups in between species comparisons are as described for Table S1. See www.stomatalevolution.org for further orthogroup and gene details. Clade subtrees (extracted from longest protein orthogroup trees available at www.stomatalevolution.org) can be viewed after copying and pasting into Microsoft Word, replacing all '^p' with nothing to remove the paragraph marks, saving as a '.txt' file and opening with Figtree (http://tree.bio.ed.ac.uk/software/figtree/).

| | Details of main orthogroup | | | | | Details of additional orthogroup | | | | | | | | | |
|---|---|---|---|---|---|---|---|---|---|---|---|---|---|---|---|
| **Gene family** | **Main orthogroup** | ***Arabidopsis thaliana*** | ***Hordeum vulgare*** | ***Polypodium vulgare*** | ***Ceratopteris richardii*** | **Additional orthogroup** | ***Arabidopsis thaliana*** | ***Hordeum vulgare*** | ***Polypodium vulgare*** | ***Ceratopteris richardii*** | **Function** | **Guard cell expression group** | **Notes** | **Related to Figure** | **Relevant clade subtree** |
| CESA | 35406 | | | | | | | | | | Cell wall | | | | |
| COBRA | 35645 | | | | | | | | | | Cell wall | | | | |
| ARCP (CSI1L) | 37233 | | | | | | | | | | Cell wall | | | | |
| KOR1L | 37843 | | | | | | | | | | Cell wall | | | | |
| EXPA | 35378 | | | | | | | | | | Cell wall | Shared | | | |
| EXPB | 35854 | | | | | | | | | | Cell wall | | | | |
| EXLA | 44550 | | | | | | | | | | Cell wall | Angio | | | |
| EXLB | 47217 | | | | | | | | | | Cell wall | | | | |
| CslA | 37631 | | | | | | | | | | Cell wall | Angio | | | |
| CslB | 35406 | | | | | | | | | | Cell wall | | | | |
| CslC | 35915 | | | | | | | | | | Cell wall | | | | |
| CslD | 35406 | | | | | | | | | | Cell wall | | | | |
| CslE | 40151 | | | | | | | | | | Cell wall | | | | |
| CslG | 40151 | | | | | | | | | | Cell wall | | | | |
| GsI | 35389 | | | | | | | | | | Cell wall | | | | |
| QUA2L | 38288 | | | | | | | | | | Cell wall | | | | |
| EXT | 37615 | | | | | | | | | | Cell wall | | | | |
| GAE | 35966 | | | | | | | | | | Cell wall | Shared | | | |
| GAUT1L/ GAUT8/ QUA1L | 35644 | | | | | | | | | | Cell wall | | | | |
| GAUT7L | 43504 | | | | | | | | | | Cell wall | | | | |
| GAUT12L | 40104 | | | | | | | | | | Cell wall | | | | |
| GAUT15L | 39278 | | | | | | | | | | Cell wall | | | | |
| PME | 35351 | | | | | | | | | | Cell wall | Shared | | | |
| CGR | 39291 | | | | | | | | | | Cell wall | | | | |
| XXT | 35605 | | | | | | | | | | Cell wall | | | | |
| LRR-RLK (FEIL) | 35961 | | | | | | | | | | Cell wall | | | | |
| RLCK | 35324 | | | | | | | | | | Cell wall | Shared | | | |
| KIN | 36869 | | | | | | | | | | Cell wall | | | | |

| | Details of main orthogroup | | | | | Details of additional orthogroup | | | | | | | | | |
|---|---|---|---|---|---|---|---|---|---|---|---|---|---|---|---|
| Gene family | Main orthogroup | *Arabidopsis thaliana* | *Hordeum vulgare* | *Polypodium vulgare* | *Ceratopteris richardii* | Additional orthogroup | *Arabidopsis thaliana* | *Hordeum vulgare* | *Polypodium vulgare* | *Ceratopteris richardii* | Function | Guard cell expression group | Notes | Related to Figure | Relevant clade subtree |
| WAK-RLK | 35376 | | | | | | | | | | Cell wall | | | | |
| MUR1L | 39626 | | | | | | | | | | Cell wall | | | | |
| FUT1L | 35868 | | | | | | | | | | Cell wall | Shared | | | |
| FUT11L | 38053 | | | | | | | | | | Cell wall | | | | |
| FUT13L | 42392 | | | | | | | | | | Cell wall | | | | |
| PGX | 35453 | | | | | | | | | | Cell wall | Shared | | | |
| FOCL1L | 45674 | | | | | | | | | | Cell wall | | | | |
| CER | 39567 | | | | | | | | | | Lipid metabolism | | | | |
| OSP1L | 35350 | | | | | | | | | | Lipid metabolism | Shared | | | |
| CDS1L | 37900 | | | | | | | | | | Lipid metabolism | | | | |
| CDS4L | 38494 | | | | | | | | | | Lipid metabolism | | | | |
| Oleosin | 36585 | | | | | | | | | | Lipid metabolism | | | | |
| MCTP | 36007 | | | | | | | | | | Lipid metabolism | Shared | | | |
| SLD | 43514 | | | | | | | | | | Lipid metabolism | | | | |
| LTP | 35515 | | | | | | | | | | Lipid metabolism | | | | |
| PAP | 35766 | | | | | | | | | | Lipid metabolism | | | | |
| ADS | 40370 | | | | | | | | | | Lipid metabolism | | | | |
| alpha/ beta-hydrolase | 39297 | | | | | | | | | | Lipid metabolism | | | | |
| SMO1-1L | 37616 | | | | | | | | | | Lipid metabolism | | | | |
| SMO2-2L | 40405 | | | | | | | | | | Lipid metabolism | | | | |
| PAT13L | 38149 | | | | | | | | | | Lipid metabolism | | | | |
| FAD3L | 36586 | | | | | | | | | | Lipid metabolism | | | | |
| FAD2L | 38879 | | | | | | | | | | Lipid metabolism | | | | |
| FAH | 37405 | | | | | | | | | | Lipid metabolism | | | | |
| KCR | 36954 | | | | | | | | | | Lipid metabolism | | | | |
| GONST | 36512 | | | | | | | | | | Lipid metabolism | | | | |
| GLOSSY1L | 35570 | | | | | | | | | | Lipid metabolism | | | | |
| PIS1L | 37398 | | | | | | | | | | Lipid metabolism | | | | |
| PGPS2L | 38491 | | | | | | | | | | Lipid metabolism | | | | |
| PLA2L | 38291 | | | | | | | | | | Lipid metabolism | | | | |
| HMGR | 36725 | | | | | | | | | | Lipid metabolism | Fern | | | |
| STE | 43939 | | | | | | | | | | Lipid metabolism | | | | |
| PI-4K | 39388 | | | | | | | | | | Lipid metabolism | | | | |
| CPT | 40194 | | | | | | | | | | Lipid metabolism | | | | |
| DES1L | 43731 | | | | | | | | | | Lipid metabolism | | | | |

| | Details of main orthogroup | | | | | Details of additional orthogroup | | | | | | | | | |
|---|---|---|---|---|---|---|---|---|---|---|---|---|---|---|---|
| Gene family | Main orthogroup | *Arabidopsis thaliana* | *Hordeum vulgare* | *Polypodium vulgare* | *Ceratopteris richardii* | Additional orthogroup | *Arabidopsis thaliana* | *Hordeum vulgare* | *Polypodium vulgare* | *Ceratopteris richardii* | Function | Guard cell expression group | Notes | Related to Figure | Relevant clade subtree |
| VDAC | 39750 | | | | | | | | | | Lipid metabolism | | | | |
| BetaHSD | 42106 | | | | | | | | | | Lipid metabolism | | | | |
| DGAT | 36624 | | | | | | | | | | Lipid metabolism | | | | |
| COX | 41308 | | | | | | | | | | Lipid metabolism | | | | |
| SETH | 43030 | | | | | | | | | | Lipid metabolism | | | | |
| LRD | 35860 | | | | | | | | | | Lipid metabolism | | | | |
| KCS | 35462 | | | | | | | | | | Lipid metabolism | Shared | | | |
| CER | 35570 | | | | | | | | | | Lipid metabolism | | | | |
| ACP | 35474 | | | | | | | | | | Lipid metabolism | | | | |
| MYB | 35331 | | | | | | | | | | Lipid metabolism | Shared | | | |
| ERF | 35329 | | | | | | | | | | Lipid metabolism | Shared | | | |
| CYP96A | 35400 | | | | | | | | | | Lipid metabolism | Shared | | | |
| CED | 41253 | | | | | | | | | | Lipid metabolism | | | | |
| PKT | 36065 | | | | | | | | | | Lipid metabolism | | | | |
| CSY | 38755 | | | | | | | | | | Lipid metabolism | | | | |
| GUP | 39099 | | | | | | | | | | Lipid metabolism | | | | |
| 4CL | 35562 | | | | | | | | | | Lipid metabolism | | | | |
| SDPs | 40993 | | | | | | | | | | Lipid metabolism | | | | |
| ACT | 35606 | | | | | | | | | | Cytoskeleton | | | | |
| PRF | 36403 | | | | | | | | | | Cytoskeleton | | | | |
| CLASP | 36736 | | | | | | | | | | Cytoskeleton | | | | |
| FRA | 36290 | | | | | | | | | | Cytoskeleton | | | | |
| KTN | 40490 | | | | | | | | | | Cytoskeleton | | | | |
| MAP20L | 38200 | | | | | | | | | | Cytoskeleton | | | | |
| MAP65L | 35611 | | | | | | | | | | Cytoskeleton | | | | |
| MAP70L | 36009 | | | | | | | | | | Cytoskeleton | | | | |
| ARP2L | 39562 | | | | | | | | | | cytoskeleton | | | | |
| ARP3L | 39044 | | | | | | | | | | cytoskeleton | | | | |
| SUS | 35658 | | | | | | | | | | Carbohydrate metabolism/ transport | | | | |
| STP | 35646 | | | | | | | | | | Carbohydrate metabolism/ transport | | | | |
| SGB1L | 35706 | | | | | | | | | | Carbohydrate metabolism/ transport | | | | |
| MFS | 35886 | | | | | | | | | | Carbohydrate metabolism/ transport | | | | |
| SUC | 36050 | | | | | | | | | | Carbohydrate metabolism/ transport | | | | |

| | Details of main orthogroup | | | | | Details of additional orthogroup | | | | | | | | | |
|---|---|---|---|---|---|---|---|---|---|---|---|---|---|---|---|
| Gene family | Main orthogroup | *Arabidopsis thaliana* | *Hordeum vulgare* | *Polypodium vulgare* | *Ceratopteris richardii* | Additional orthogroup | *Arabidopsis thaliana* | *Hordeum vulgare* | *Polypodium vulgare* | *Ceratopteris richardii* | Function | Guard cell expression group | Notes | Related to Figure | Relevant clade subtree |
| SWEET | 35480 | | | | | | | | | | Carbohydrate metabolism/ transport | Shared | | | |
| SWEET11L | 38674 | | | | | | | | | | Carbohydrate metabolism/ transport | | | | |
| SPS | 37098 | | | | | | | | | | Carbohydrate metabolism/ transport | | | | |
| UGE | 35933 | | | | | | | | | | Carbohydrate metabolism/ transport | | | | |
| cINV | 35550 | | | | | | | | | | Carbohydrate metabolism/ transport | | | | |
| cwINV | 35593 | | | | | | | | | | Carbohydrate metabolism/ transport | | | | |
| FUM | 40809 | | | | | | | | | | Carbohydrate metabolism/ transport | | | | |
| INVH | 35550 | | | | | | | | | | Carbohydrate metabolism/ transport | | | | |
| PMEI | 36248 | | | | | | | | | | Carbohydrate metabolism/ transport | | | | |
| iPGAM | 37181 | | | | | | | | | | Carbohydrate metabolism/ transport | Fern | | | |
| TPT | 37598 | | | | | | | | | | Carbohydrate metabolism/ transport | | | | |
| APL | 35775 | | | | | | | | | | Carbohydrate metabolism/ transport | | | | |
| AMY | 35721 | | | | | | | | | | Carbohydrate metabolism/ transport | | | | |
| APS | 35775 | | | | | | | | | | Carbohydrate metabolism/ transport | | | | |
| BAM | 35538 | | | | | | | | | | Carbohydrate metabolism/ transport | Shared | | | |
| BAM | 49925 | | | | | | | | | | Carbohydrate metabolism/ transport | | | | |
| GWD | 36648 | | | | | | | | | | Carbohydrate metabolism/ transport | | | | |
| PTPC | 35920 | | | | | | | | | | Carbohydrate metabolism/ transport | | | | |
| BTPC | 39397 | | | | | | | | | | Carbohydrate metabolism/ transport | | | | |
| PEPCK1L | 37136 | | | | | | | | | | Carbohydrate metabolism/ transport | | | | |
| PPCK2L | 41409 | | | | | | | | | | Carbohydrate metabolism/ transport | | | | |
| PPDK | 37041 | | | | | | | | | | Carbohydrate metabolism/ transport | | | | |
| cyNAD-MDH | 39476 | | | | | | | | | | Carbohydrate metabolism/ transport | | | | |

| | Details of main orthogroup | | | | | Details of additional orthogroup | | | | | | | | | |
|---|---|---|---|---|---|---|---|---|---|---|---|---|---|---|---|
| **Gene family** | **Main orthogroup** | ***Arabidopsis thaliana*** | ***Hordeum vulgare*** | ***Polypodium vulgare*** | ***Ceratopteris richardii*** | **Additional orthogroup** | ***Arabidopsis thaliana*** | ***Hordeum vulgare*** | ***Polypodium vulgare*** | ***Ceratopteris richardii*** | **Function** | **Guard cell expression group** | **Notes** | **Related to Figure** | **Relevant clade subtree** |
| pNAD-MDH | 36156 | | | | | | | | | | Carbohydrate metabolism/ transport | | | | |
| NADP-MDH | 39943 | | | | | | | | | | Carbohydrate metabolism/ transport | | | | |
| NADP-ME | 35828 | | | | | | | | | | Carbohydrate metabolism/ transport | | | | |
| ISA | 35835 | | | | | | | | | | Carbohydrate metabolism/ transport | | | | |
| LDA | 38520 | | | | | | | | | | Carbohydrate metabolism/ transport | | | | |
| HAD | 35510 | | | | | | | | | | Carbohydrate metabolism/ transport | Shared | | | |
| GWD | 36648 | | | | | | | | | | Carbohydrate metabolism/ transport | | | | |
| HXK | 35858 | | | | | | | | | | Carbohydrate metabolism/ transport | | | | |
| PFK | 35650 | | | | | | | | | | Carbohydrate metabolism/ transport | | | | |
| PFK5L | 36541 | | | | | | | | | | Carbohydrate metabolism/ transport | | | | |
| PFK | 36206 | | | | | | | | | | Carbohydrate metabolism/ transport | | | | |
| PFK | 38342 | | | | | | | | | | Carbohydrate metabolism/ transport | | | | |
| PGI | 37567 | | | | | | | | | | Carbohydrate metabolism/ transport | | | | |
| PGM | 37331 | | | | | | | | | | Carbohydrate metabolism/ transport | Fern | | | |
| SDH2 | 41318 | | | | | | | | | | Carbohydrate metabolism/ transport | | | | |
| SPS | 37098 | | | | | | | | | | Carbohydrate metabolism/ transport | | | | |
| tDT | 37275 | | | | | | | | | | Carbohydrate metabolism/ transport | | | | |
| PML | 39305 | | | | | | | | | | Transport | | | | |
| BOR | 35551 | | | | | | | | | | Transport | Shared | | | |
| AMT | 37154 | | | | | | | | | | Transport | | | | |
| AMT | 36938 | | | | | | | | | | Transport | Angio | | | |
| NRT1 | 35338 | | | | | | | | | | Transport | Shared | | | |
| NRT2 | 36706 | | | | | | | | | | Transport | | | | |
| NRT3.1L | 37988 | | | | | | | | | | Transport | | | | |
| NRT3.2L | 42142 | | | | | | | | | | Transport | | | | |
| ALMT | 35531 | | | | | | | | | | Transport | Shared | | Figure 2 | |
| SLAC1/ SLAH2/ SLAH3L | 35940 | | | | | | | | | | Transport | Shared | | Figure 2 | |

| | Details of main orthogroup | | | | | Details of additional orthogroup | | | | | | | | | |
|---|---|---|---|---|---|---|---|---|---|---|---|---|---|---|---|
| **Gene family** | **Main orthogroup** | ***Arabidopsis thaliana*** | ***Hordeum vulgare*** | ***Polypodium vulgare*** | ***Ceratopteris richardii*** | **Additional orthogroup** | ***Arabidopsis thaliana*** | ***Hordeum vulgare*** | ***Polypodium vulgare*** | ***Ceratopteris richardii*** | **Function** | **Guard cell expression group** | **Notes** | **Related to Figure** | **Relevant clade subtree** |
| SLAH1/4L | 48967 | | | | | | | | | | Transport | | | | |
| PHO | 35669 | | | | | | | | | | Transport | Shared | | | |
| CAT | 36141 | | | | | | | | | | Transport | Shared | | | |
| Shaker Kin (KAT/AKT) | 36099 | | | | | | | | | | Transport | | | Figure 2 | |
| Shaker Kout (ORK) | 36393 | | | | | | | | | | Transport | | | Figure 2 | |
| KUP/ HAK/ KT | 35356 | | | | | | | | | | Transport | | | | |
| TPK/KCO | 35878 | | | | | | | | | | Transport | | | | |
| HKT | 37905 | | | | | | | | | | Transport | | | | |
| NHX1-4L | 36041 | | | | | | | | | | Transport | Shared | | | |
| NHX5/6L | 35563 | | | | | | | | | | Transport | Angio | | | |
| NHX7-8L | 35954 | | | | | | | | | | Transport | | | | |
| CHX | 35623 | | | | | | | | | | Transport | Shared | | | |
| ABCB | 35374 | | | | | | | | | | Transport | Shared | | | |
| ABCC | 35379 | | | | | | | | | | Transport | | | | |
| ABCG22/ 25L | 35745 | | | | | | | | | | Transport | Shared | | | |
| ABCG40L | 35358 | | | | | | | | | | Transport | Shared | | | |
| CLC | 35800 | | | | | | | | | | Transport | | | | |
| CLCe-f | 36858 | | | | | | | | | | Transport | | | | |
| VHA Subunit A | 40326 | | | | | | | | | | Transport | | | | |
| VHA Subunit B | 38569 | | | | | | | | | | Transport | | | | |
| VHA Subunit C | 42388 | | | | | | | | | | Transport | | | | |
| VHA Subunit D | 42725 | | | | | | | | | | Transport | | | | |
| VHA Subunit E | 37885 | | | | | | | | | | Transport | | | | |
| VHA Subunit F | 42748 | | | | | | | | | | Transport | | | | |
| VHA Subunit G | 41315 | | | | | | | | | | Transport | | | | |
| VHA Subunit H | 41797 | | | | | | | | | | Transport | | | | |
| VHASubunita | 36409 | | | | | | | | | | Transport | | | | |
| VHA Subunit c | 37527 | | | | | | | | | | Transport | | | | |
| VHA Subunit c” | 41653 | | | | | | | | | | Transport | | | | |
| VHA Subunit d | 42789 | | | | | | | | | | Transport | | | | |
| VHS Subunit e | 44409 | | | | | | | | | | Transport | Angio | | | |
| Aquaporins-PIP | 35476 | | | | | | | | | | Transport | Shared | | | |
| Aquaporins-TIP | 36055 | | | | | | | | | | Transport | | | | |
| Aquaporins-NIP | 35936 | | | | | | | | | | Transport | | | | |
| ACA (ECA1-4L) | 35702 | | | | | | | | | | Transport | | | | |
| ACA | 35478 | | | | | | | | | | Transport | | | | |
| AUX | 36320 | | | | | | | | | | Transport | Shared | | | |
| CAX1-6L | 35979 | | | | | | | | | | Transport | | | | |
| CAX7-10L | 39164 | | | | | | | | | | Transport | | | | |
| CAX11L | 44884 | | | | | | | | | | Transport | | | | |
| CNGC1 | 35616 | | | | | | | | | | Transport | | | | |

| | Details of main orthogroup | | | | | Details of additional orthogroup | | | | | | | | | |
|---|---|---|---|---|---|---|---|---|---|---|---|---|---|---|---|
| **Gene family** | **Main orthogroup** | ***Arabidopsis thaliana*** | ***Hordeum vulgare*** | ***Polypodium vulgare*** | ***Ceratopteris richardii*** | **Additional orthogroup** | ***Arabidopsis thaliana*** | ***Hordeum vulgare*** | ***Polypodium vulgare*** | ***Ceratopteris richardii*** | **Function** | **Guard cell expression group** | **Notes** | **Related to Figure** | **Relevant clade subtree** |
| CNGC2 | 39492 | | | | | | | | | | Transport | | | | |
| GLR | 35362 | | | | | | | | | | Transport | Shared | | | |
| GPT1L | 37995 | | | | | | | | | | Transport | | | | |
| AVP1L | 37097 | | | | | | | | | | Transport | | | | |
| AVP2L | 39324 | | | | | | | | | | Transport | | | | |
| KEA1-2L | 37476 | | | | | | | | | | Transport | | | | |
| KEA3L | 37303 | | | | | | | | | | Transport | | | | |
| MATE (RHC1/ DTX50L) | 36017 | | | | | | | | | | Transport | Shared | | | |
| MSL1L | 40096 | | | | | | | | | | Transport | | | | |
| MSL2/3L | 37037 | | | | | | | | | | Transport | | | | |
| MSL4-10L | 36163 | | | | | | | | | | Transport | | | | |
| OSCA1/2L | 35523 | | | | | | | | | | Transport | | | | |
| OSCA2.1L | 35350 | | | | | | | | | | Transport | Shared | | | |
| OSCA3L | 37027 | | | | | | | | | | Transport | | | | |
| OSCA4L | 43875 | | | | | | | | | | Transport | | | | |
| PIL | 37762 | | | | | | | | | | Transport | | | | |
| PIN | 35513 | | | | | | | | | | Transport | | | | |
| PLT | 36134 | | | | | | | | | | Transport | | | | |
| TPC1 | 36664 | | | | | | | | | | Transport | | | | |
| HOS | 40790 | | | | | | | | | | Signalling | | | | |
| HAD | 36347 | | | | | | | | | | Signalling | | | | |
| ABF | 35542 | | | | | | | | | | Signalling | | | | |
| Alix | 39206 | | | | | | | | | | Signalling | | | | |
| AlphaCA | 35675 | | | | | | | | | | Signalling | | | | |
| AHA | 35426 | | | | | | | | | | Signalling | Shared | | | |
| HK | 35659 | | | | | | | | | | Signalling | | | | |
| Alpha Beta Hydrolase | 36162 | | | | | | | | | | Signalling | | | | |
| ARF | 36896 | | | | | | | | | | Signalling | | | | |
| ARF | 35529 | | | | | | | | | | Signalling | | | | |
| ASK | 35504 | | | | | | | | | | Signalling | | | | |
| AXRL | 38379 | | | | | | | | | | Signalling | | | | |
| BCA | 35806 | | | | | | | | | | Signalling | | | | |
| bHLH(PIF) | 35354 | | | | | | | | | | Signalling | Shared | | | |
| bHLH (MYC/ JAML) | 35372 | | | | | | | | | | Signalling | Shared | | | |
| bHLH (BIM2) | 36401 | | | | | | | | | | Signalling | | | | |
| Bonzai | 36174 | | | | | | | | | | Signalling | | | | |
| BSK | 36076 | | | | | | | | | | Signalling | | | | |
| BSL | 35874 | | | | | | | | | | Signalling | | | | |
| bZIP (TGA1L) | 35541 | | | | | | | | | | Signalling | | | | |
| bZIP (VIP1L) | 35576 | | | | | | | | | | Signalling | | | | |
| BZL | 36124 | | | | | | | | | | Signalling | | | | |

| | Details of main orthogroup | | | | | Details of additional orthogroup | | | | | | | | | |
|---|---|---|---|---|---|---|---|---|---|---|---|---|---|---|---|
| **Gene family** | **Main orthogroup** | ***Arabidopsis thaliana*** | ***Hordeum vulgare*** | ***Polypodium vulgare*** | ***Ceratopteris richardii*** | **Additional orthogroup** | ***Arabidopsis thaliana*** | ***Hordeum vulgare*** | ***Polypodium vulgare*** | ***Ceratopteris richardii*** | **Function** | **Guard cell expression group** | **Notes** | **Related to Figure** | **Relevant clade subtree** |
| CAR | 35617 | | | | | | | | | | Signalling | | | | |
| CaS | 39653 | | | | | | | | | | Signalling | | | | |
| CBL | 35473 | | | | | | | | | | Signalling | | | | |
| CDK | 37952 | | | | | | | | | | Signalling | | | | |
| CIPK | 35388 | | | | | | | | | | Signalling | | & AT5G01820 CIPK14 and AT2G38490 CIPK22 (missing from orthogroup) | Figure 2 | |
| CKL | 35545 | | | | | | | | | | Signalling | | | | |
| CLE25L | 44375 | | | | | | | | | | Signalling | | | | |
| CLE9L | 43070 | | | | | | | | | | Signalling | Shared | | | |
| cMCU | 36674 | | | | | | | | | | Signalling | | | | |
| COP1L | 35797 | | | | | | | | | | Signalling | | | | |
| SnRK2 | 35613 | | | | | | | | | | Signalling | | OST1/SnRK 2-2/SnRK2-3 subclade shown in Figure 2 | Figure 2 | (('Ma_po_Mapoly0011s0096.1.p/228816':0.31932692576420263,'Ma_po_Mapoly0061s0075.1.p/235162':0.06464164443529063):0.061312308292645445,((('Ph_pa_Pp3c6_11090V3.1.p/265406':0.05546258106756108,'Ph_pa_Pp3c5_21160V3.1.p/256924':0.026584389146407705):0.009182994805270628,('Ph_pa_Pp3c5_17150V3.1.p/257575':0.12952923723293486,'Ph_pa_Pp3c6_16600V3.1.p/265847':0.011974772231543529):0.020032656534016535):0.06917538979998783,((('Gi_bi_Gb_18878/126574':1.1530081976331668,'Pi_ab_MA_10430788g0010/273596':0.0707880971271515):0.017444848062922347,('Gi_bi_Gb_32504/120822':1.0000004999621837E-6,'Gi_bi_Gb_15703/146974':0.20615092921586586):0.021185175186343885):0.024077193432233468,(((('Po_vu_TRINITY_DN74059_c8_g2__TRINITY_DN74059_c8_g2_i4__g.50298__m.50298/374218':0.09523960773512652,('Ce_ri_TRINITY_DN42080_c5_g1__TRINITY_DN42080_c5_g1_i13__g.4609__m.4609/74095':0.21489954031671665,'Sa_cu_Sacu_v1.1_s0258.g026859/405766':0.4487469815645444):0.017857302765118988):0.05793030638093999,('Ce_ri_TRINITY_DN42080_c5_g2__TRINITY_DN42080_c5_g2_i12__g.4610__m.4610/74096':0.1082438771150469,('Po_vu_TRINITY_DN65559_c1_g2__TRINITY_DN65559_c1_g2_i5__g.165286__m.165286/359914':0.13633031148652552,'Sa_cu_Sacu_v1.1_s0061.g015371/397344':0.18294286602184195):0.030909244159779137):0.03643976446100594):0.16964021745423286,('Ch_br_g22934/90921':0.43888038560038245,('Ce_ri_TRINITY_DN42080_c4_g1__TRINITY_DN42080_c4_g1_i1__g.4584__m.4584/74094':1.0000004999621837E-6,'Kl_ni_kfl00097_0360_v1.1/204793':0.011152375836305461):0.28321170271042106):0.007427184496213712):0.05473579944084728,(('Pi_ab_MA_490939g0010/316886':0.6860929134183975,'Gi_bi_Gb_02396/139075':0.13926538361754348):0.05160457117590056,(('Pi_ab_MA_10231938g0010/297287':1.0000005001842283E-6,'Pi_ab_MA_178212g0010/309414':1.0000005001842283E-6):0.33797823896867585,(('Pi_ab_MA_10431614g0020/273991':0.18066999913710347,('Pi_ab_MA_7213602g0010/321083':0.161961944460063 5,'Pi_ab_MA_8718813g0010/324351':0.20655170154745206):0.07892747482035256):0.19432883151300318,('Am_tr_evm_27.model.AmTr_v1.0_scaffold00033.261/12619':0.04096428304103483,(('Ce_ri_TRINITY_DN26242_c0_g1__TRINITY_DN26242_c0_g1_i1__g.179728__m.179728/56822':0.11332510677628349,'Ar_th_AT4G33950.1/46899':0.06188966877521817 6):0.05327448154095693,(('Ho_vu_HORVU5Hr1G097630.1/168843':0.10651067708648587,('Ho_vu_HORVU4Hr1G013540.5/180029':0.047562636099467204,('Ho_vu_HORVU0Hr1G011570.1/165149':0.0018681538311962065,'Ho_vu_HORVU5Hr1G018340.5/183146':0.19997960958764072):0.20006011196860563):0.04736682682254245):0.02936866956989758 7,('Ar_th_AT3G50500.2/47659':0.10206600061043059,'Ar_th_AT5G66880.2/31849':0.04690834572566271 5):0.1761471320176553):0.0480994983420453):0.014279853770194695):0.09255862123998382):0.05946762682476936):0.0041272796136906376):0.026158261500249358):0.03212166359881219):0.088169895359973):0.02771168609721375):0.006229851401317552; |
| SnRK1 | 35946 | | | | | | | | | | Signalling | | | | |
| CPK | 35391 | | | | | 37344 | | | | | Signalling | | only CPK subclade included from 35391 (PEPKR1 genes excluded) | Figure 2 | |
| CRY | 36219 | | | | | | | | | | Signalling | | | | |
| CTR | 35421 | | | | | | | | | | Signalling | | | | |
| Cullin | 36813 | | | | | | | | | | Signalling | | | | |
| CYP707 | 35386 | | | | | | | | | | Signalling | Shared | | | |
| CYP21-1L | 35703 | | | | | | | | | | Signalling | | | | |
| CYP21-3L | 39907 | | | | | | | | | | Signalling | Fern | | | |

| | Details of main orthogroup | | | | | Details of additional orthogroup | | | | | | | | | |
|---|---|---|---|---|---|---|---|---|---|---|---|---|---|---|
| Gene family | Main orthogroup | *Arabidopsis thaliana* | *Hordeum vulgare* | *Polypodium vulgare* | *Ceratopteris richardii* | Additional orthogroup | *Arabidopsis thaliana* | *Hordeum vulgare* | *Polypodium vulgare* | *Ceratopteris richardii* | Function | Guard cell expression group | Notes | Related to Figure | Relevant clade subtree |
| CYP74 | 36996 | | | | | | | | | | Signalling | | | | |
| CYP79 | 46381 | | | | | | | | | | Signalling | | | | |
| CYP80 | 46381 | | | | | | | | | | Signalling | | | | |
| CYP90 | 35386 | | | | | | | | | | Signalling | Shared | | | |
| CYP97 | 36619 | | | | | | | | | | Signalling | | | | |
| CYP701 | 38775 | | | | | | | | | | Hormone metabolism (GA) | Fern | | | |
| CYP711 (MAX1) | 41753 | | | | | | | | | | Hormone metabolism (SL) | Angio | | | |
| PEPR | 35335 | | | | | | | | | | Signalling | Shared | | | |
| Dehydrin | 39942 | | | | | | | | | | Signalling | | | | |
| ERD | 38121 | | | | | | | | | | Signalling | | | | |
| LEA | 42300 | | | | | | | | | | Signalling | | | | |
| DELLA | 36108 | | | | | | | | | | Signalling | | | | |
| E3Ubi | 41950 | | | | | | | | | | Signalling | | | | |
| EAR | 43487 | | | | | | | | | | Signalling | | | | |
| EIN3L | 36189 | | | | | | | | | | Signalling | | | | |
| EIN2L | 38987 | | | | | | | | | | Signalling | | | | |
| FOX | 35742 | | | | | | | | | | Signalling | | | | |
| TIR1L | 35460 | | | | | | | | | | Signalling | | | | |
| EBF1L | 37333 | | | | | | | | | | Signalling | | | | |
| MAX2L | 40311 | | | | | | | | | | Signalling | | | | |
| FREE1L | 41067 | | | | | | | | | | Signalling | | | | |
| GAD | 35838 | | | | | | | | | | Signalling | | | | |
| PP2C (Group A) | 35624 | | | | | | | | | | Signalling | | | Figure 2 | |
| GCR | 37620 | | | | | | | | | | Signalling | | | | |
| GH3 | 35471 | | | | | | | | | | Signalling | | | | |
| GC1L | 36323 | | | | | | | | | | Signalling | | | | |
| GID | 35385 | | | | | | | | | | Signalling | | | | |
| GPA1L | 40428 | | | | | | | | | | Signalling | | | | |
| GB1L | 40464 | | | | | | | | | | Signalling | | | | |
| AGG1L | 41271 | | | | | | | | | | Signalling | | | | |
| GPX | 36196 | | | | | | | | | | Signalling | | | | |
| GTG | 39760 | | | | | | | | | | Signalling | | | | |
| HVA22 | 36093 | | | | | | | | | | Signalling | Shared | | | |
| IAA | 35444 | | | | | | | | | | Signalling | Shared | | | |
| JMJ | 36879 | | | | | | | | | | Signalling | | | | |
| KAPP | 38472 | | | | | | | | | | Signalling | | | | |
| LOX | 35363 | | | | | | | | | | Signalling | Shared | | | |
| LOT1 | 43042 | | | | | | | | | | Signalling | | | | |
| LRX | 35826 | | | | | | | | | | Signalling | Shared | | | |
| MATH-BTB | 36426 | | | | | | | | | | Signalling | | | | |

| | Details of main orthogroup | | | | | Details of additional orthogroup | | | | | | | | | |
|---|---|---|---|---|---|---|---|---|---|---|---|---|---|---|---|
| Gene family | Main orthogroup | *Arabidopsis thaliana* | *Hordeum vulgare* | *Polypodium vulgare* | *Ceratopteris richardii* | Additional orthogroup | *Arabidopsis thaliana* | *Hordeum vulgare* | *Polypodium vulgare* | *Ceratopteris richardii* | Function | Guard cell expression group | Notes | Related to Figure | Relevant clade subtree |
| MOCA | 37824 | | | | | | | | | | Signalling | | | | |
| MAPK (MPK4/ 12L) | 35689 | | | | | | | | | | Signalling | Shared | | | |
| MAPK (MPK9L) | 35459 | | | | | | | | | | Signalling | | | | |
| MAPKK (MKK1/ 2L) | 37632 | | | | | | | | | | Signalling | | | | |
| MAPKK (MKK3L) | 39023 | | | | | | | | | | Signalling | | | | |
| MAPKK (MKK4/ 5L) | 39382 | | | | | | | | | | Signalling | | | | |
| MAPKKK (BHP1L) | 36075 | | | | | | | | | | Signalling | | | | |
| MAPKKK (VIKL) | 37363 | | | | | | | | | | Signalling | | | | |
| MAPKKK (HT1L) | 35904 | | | | | | | | | | Signalling | Shared | | | |
| MAPKKK (CBCL) | 37214 | | | | | | | | | | Signalling | | | | |
| MAPKKK (RAF13L) | 38013 | | | | | | | | | | Signalling | | | | |
| MAPKKKK (BLUS1L) | 35711 | | | | | | | | | | Signalling | | | | |
| MAPKKKK (MAP4K ALPHA 1L) | 37842 | | | | | | | | | | Signalling | | | | |
| MAPKKKK (SIK1L) | 40345 | | | | | | | | | | Signalling | | | | |
| MED | 39772 | | | | | | | | | | Signalling | | | | |
| PUB | 35371 | | | | | | | | | | Signalling | | | | |
| MgChe | 39128 | | | | | | | | | | Signalling | | | | |
| MUN | 35973 | | | | | | | | | | Signalling | | | | |
| NINJA | 36987 | | | | | | | | | | Signalling | | | | |
| NPR | 37248 | | | | | | | | | | Signalling | | | | |
| NRGA | 38495 | | | | | | | | | | Signalling | | | | |
| PHOT | 36528 | | | | | | | | | | Signalling | | | | |
| PHY | 36292 | | | | | | | | | | Signalling | | | | |
| ETR | 35963 | | | | | | | | | | Signalling | | | | |
| PLC | 35676 | | | | | | | | | | Signalling | | | | |
| PLD | 35496 | | | | | | | | | | Signalling | Shared | | | |
| PP1 | 36241 | | | | | | | | | | Signalling | | | | |
| PRMT | 39216 | | | | | | | | | | Signalling | | | | |
| PRSL | 36095 | | | | | | | | | | Signalling | Shared | | | |
| PRX | 35328 | | | | | | | | | | Signalling | Shared | | | |
| PYR/ PYL/ RCAR | 36014 | | | | | | | | | | Signalling | | & AT5G45860 PYL11 and AT4G18620 PYL13 (missing from orthogroup) | Figure 2 | |
| RALF | 36888 | | | | | | | | | | Signalling | | | | |
| RAN GTPase | 37306 | | | | | | | | | | Signalling | | | | |
| RAV | 35720 | | | | | | | | | | Signalling | | | | |
| RBOH | 35411 | | | | | | | | | | Signalling | Shared | | | |
| RBX | 41424 | | | | | | | | | | Signalling | | | | |

| | Details of main orthogroup | | | | | Details of additional orthogroup | | | | | | | | | |
|---|---|---|---|---|---|---|---|---|---|---|---|---|---|---|---|
| **Gene family** | **Main orthogroup** | ***Arabidopsis thaliana*** | ***Hordeum vulgare*** | ***Polypodium vulgare*** | ***Ceratopteris richardii*** | **Additional orthogroup** | ***Arabidopsis thaliana*** | ***Hordeum vulgare*** | ***Polypodium vulgare*** | ***Ceratopteris richardii*** | **Function** | **Guard cell expression group** | **Notes** | **Related to Figure** | **Relevant clade subtree** |
| RING-H2 | 35336 | | | | | | | | | | Signalling | Shared | | | |
| RLCK | 35759 | | | | | | | | | | Signalling | | | | |
| LRR-RLK (GHR1L) | 37702 | | | | | | | | | | Signalling | | | | |
| LRR-RLK (FLS2/ EFR/ PEPR/ BAML) | 35335 | | | | | | | | | | Signalling | Shared | | | |
| LRR-RLK (KIN7L) | 35483 | | | | | | | | | | Signalling | Shared | | | |
| LRR-RLK (BAK1L) | 35337 | | | | | | | | | | Signalling | Shared | | | |
| LRR-RLK (BRI1L) | 39875 | | | | | | | | | | Signalling | | | | |
| LRR-RLK (HPCA1L) | 35564 | | | | | | | | | | Signalling | | | | |
| RLCK | 35324 | | | | | | | | | | Signalling | Shared | | | |
| LysM-RLK (CERK1L) | 37000 | | | | | | | | | | Signalling | | | | |
| LysM-RLK (LYK5L) | 35530 | | | | | | | | | | Signalling | Angio | | | |
| WAK-RLK | 35376 | | | | | | | | | | Signalling | | | | |
| ROP-GTPase | 35788 | | | | | | | | | | Signalling | | | | |
| SAUR | 35369 | | | | | | | | | | Signalling | Shared | | | |
| SBPase | 43329 | | | | | | | | | | Signalling | | | | |
| SCS | 38940 | | | | | | | | | | Signalling | | | | |
| SCD | 38479 | | | | | | | | | | Signalling | | | | |
| SKPL | 36523 | | | | | | | | | | Signalling | | | | |
| SLYL | 36639 | | | | | | | | | | Signalling | | | | |
| SOD | 37705 | | | | | | | | | | Signalling | | | | |
| SPA | 35797 | | | | | | | | | | Signalling | | | | |
| TPR | 35865 | | | | | | | | | | Signalling | | | | |
| PI3K | 37386 | | | | | | | | | | Signalling | | | | |
| VSR | 35859 | | | | | | | | | | Signalling | | | | |
| WRKY | 35346 | | | | | | | | | | Signalling | Shared | | | |
| YAK | 36779 | | | | | | | | | | Signalling | | | | |
| ZIM | 35590 | | | | | | | | | | Signalling | | | | |
| MCTP | 36007 | | | | | | | | | | Signalling | Shared | | | |
| GSTF | 35554 | | | | | | | | | | Signalling | Shared | | | |
| GSTU | 35402 | | | | | | | | | | Signalling | | | | |
| GSTT | 37402 | | | | | | | | | | Signalling | Angio | | | |
| GSTZ | 37634 | | | | | | | | | | Signalling | | | | |
| DHAR | 37989 | | | | | | | | | | Signalling | | | | |
| GST (TCHQD1L) | 41145 | | | | | | | | | | Signalling | | | | |
| PER | 35328 | | | | | | | | | | Signalling | Shared | | | |
| MLO | 35446 | | | | | | | | | | Signalling | Angio | | | |
| SAI-LLP | 35353 | | | | | | | | | | Signalling | Shared | | | |
| RPS | 35330 | | | | | | | | | | Signalling | | | | |
| JASSYL | 39702 | | | | | | | | | | Signalling | | | | |

| | Details of main orthogroup | | | | | Details of additional orthogroup | | | | | | | | | |
|---|---|---|---|---|---|---|---|---|---|---|---|---|---|---|---|
| Gene family | Main orthogroup | *Arabidopsis thaliana* | *Hordeum vulgare* | *Polypodium vulgare* | *Ceratopteris richardii* | Additional orthogroup | *Arabidopsis thaliana* | *Hordeum vulgare* | *Polypodium vulgare* | *Ceratopteris richardii* | Function | Guard cell expression group | Notes | Related to Figure | Relevant clade subtree |
| 14-3-3s | 35818 | | | | | | | | | | Signalling | | | | |
| SDR110C | 35501 | | | | | | | | | | Hormone metabolism (ABA) | Shared | whole orthogroup shown in Figure S1 with ABA2 subclade highlighted | Figure S1 | (Am_tr_evm_27.model.AmTr_v1.0_scaffold00053.71/5857:0.81092272935430598224,(Ar_th_AT1G52340.1/29433:0.75632144633914455412,(Ho_vu_HORVU5Hr1G111190.1/173328:0.00000100000050002909,Ho_vu_HORVU3Hr1G046320.1/198779:0.00000100000050002909)100:0.58229176861302223145)78:0.27737758916065641257)81:0.40243735158871463131; |
| PSY | 37068 | | | | | | | | | | Hormone metabolism (ABA) | | | | |
| PKT | 36065 | | | | | | | | | | Hormone metabolism (JA) | | | | |
| PDS | 40812 | | | | | | | | | | Hormone metabolism (ABA) | | | | |
| JMT | 35409 | | | | | | | | | | Hormone metabolism (JA) | | | | |
| GA20ox3L | 39481 | | | | | | | | | | Hormone metabolism (GA) | | | | |
| GA3ox1L | 36160 | | | | | | | | | | Hormone metabolism (GA) | | | | |
| ABA4 | 39542 | | | | | | | | | | Hormone metabolism (ABA) | | | Figure S1 | |
| AMI | 35999 | | | | | | | | | | Hormone metabolism (IAA) | | | | |
| ACC oxidase | 49788 | | | | | | | | | | Hormone metabolism (C2H4) | | | | |
| ACC synthase | 37495 | | | | | | | | | | Hormone metabolism (C2H4) | | | | |
| ACX | 36593 | | | | | | | | | | Hormone metabolism (JA) | | | | |
| AIM | 36633 | | | | | | | | | | Hormone metabolism (JA) | | | | |
| AO | 36044 | | | | | | | | | | Hormone metabolism (ABA+IAA) | | | Figure S1 | |
| 3Oxo5A | 45345 | | | | | | | | | | Hormone metabolism (SL) | | | | |
| AOC | 38703 | | | | | | | | | | Hormone metabolism (JA) | Fern | | | |
| BCH | 36580 | | | | | | | | | | Hormone metabolism (ABA) | | | Figure S1 | |

| | Details of main orthogroup | | | | | Details of additional orthogroup | | | | | | | | | |
|---|---|---|---|---|---|---|---|---|---|---|---|---|---|---|---|
| Gene family | Main orthogroup | *Arabidopsis thaliana* | *Hordeum vulgare* | *Polypodium vulgare* | *Ceratopteris richardii* | Additional orthogroup | *Arabidopsis thaliana* | *Hordeum vulgare* | *Polypodium vulgare* | *Ceratopteris richardii* | Function | Guard cell expression group | Notes | Related to Figure | Relevant clade subtree |
| NCED+ CCD4 | 36464 | | | | | | | | | | Hormone metabolism (ABA) | | only NCED subclade shown in Figure S1 | Figure S1 | ('Ma_po_Mapoly0149s0036.1.p/221703':1.1427605554842906,((('Ph_pa_Pp3c25_4816V3.1.p/266742':1.0000005001842283E-6,'Ph_pa_Pp3c25_4810V3.1.p/267023':1.0000005001842283E-6):0.031174786437564883,'Ph_pa_Pp3c16_17210V3.1.p/269154':0.16033337780895618):0.7591101873161596,(('Ma_po_Mapoly0015s0066.1.p/230036':0.49624596651155395,'Se_mo_233638/407294':0.6587038894098876):0.11268535441004168,(((('Pi_ab_MA_10428505g0020/272579':0.22391161795467163,'Pi_ab_MA_10428505g0010/272578':0.09679645095739131):0.18262605273987065,('Gi_bi_Gb_07367/142582':0.3095161344752744,('Pi_ab_MA_10434448g0010/275231':0.21031965138266795,'Gi_bi_Gb_19071/146161':0.16094748825833705):0.1011456919323348):0.0536077642578805):0.0569699478676122,('Pi_ab_MA_10174788g0010/270145':0.5915630699925951,('Gi_bi_Gb_14132/159949':0.889677226554408,'Gi_bi_Gb_14131/159948':0.03262560098282474):0.28522247040442306):0.2522995074043335):0.08200170013466979,((('Am_tr_evm_27.model.AmTr_v1.0_scaffold00092.158/2755':0.45428743279571604,(('Ho_vu_HORVU5Hr1G008050.1/179049':0.17776512711673798,('Ho_vu_HORVU5Hr1G044510.1/178309':0.0016505006268943134,('Ho_vu_HORVU5Hr1G092850.2/167160':1.0000005001842283E-6,'Ho_vu_HORVU5Hr1G054970.1/167906':1.0000005001842283E-6):0.0535595357315648):0.23440679662935437):0.31818506383534517,(('Ar_th_AT4G18350.1/28289':0.33638952257786014,'Ar_th_AT1G30100.1/42047':0.34061432057286645):0.194074793493765,('Ar_th_AT1G78390.1/46862':0.3110460517710756,'Ar_th_AT3G14440.1/43429':0.2233865035437121 4):0.09048380645446574):0.07951816788576771):0.1463279073921986):0.1610637719536534,('Am_tr_evm_27.model.AmTr_v1.0_scaffold00039.158/15918':0.6113488227740649,'Ar_th_AT3G24220.1/43720':0.7834022881841483):0.19010227069522534):0.12649130210871729,((('Ce_ri_TRINITY_DN43034_c1_g1__TRINITY_DN43034_c1_g1_i1__g.86231__m.86231/76381':0.538080362336141,'Po_vu_TRINITY_DN76298_c1_g1__TRINITY_DN76298_c1_g1_i5__g.129132__m.129132/378134':0.2323895589762528):0.18476030658549325,(('Sa_cu_Sacu_v1.1_s0149.g023278/403178':0.6550500912411952,('Ce_ri_TRINITY_DN38658_c0_g1__TRINITY_DN38658_c0_g1_i4__g.140841__m.140841/66037':0.650146506509012,'Sa_cu_Sacu_v1.1_s0002.g001124/387813':0.42746108677419326):0.066127197760526):0.1149411055093914,('Sa_cu_Sacu_v1.1_s0158.g023803/403534':0.548803441035921,(('Po_vu_TRINITY_DN72995_c3_g1__TRINITY_DN72995_c3_g1_i4__g.4669__m.4669/372471':0.20077266147347617,'Ce_ri_TRINITY_DN28724_c0_g1__TRINITY_DN28724_c0_g1_i1__g.3415__m.3415/57304':0.37378309008634325):0.04383739612608606,(('Ce_ri_TRINITY_DN31580_c0_g1__TRINITY_DN31580_c0_g1_i4__g.138823__m.138823/58642':0.5176910948134004,'Po_vu_TRINITY_DN79474_c1_g1__TRINITY_DN79474_c1_g1_i4__g.247760__m.247760/383790':0.12893167849846643):0.0593199337950611,'Po_vu_TRINITY_DN72995_c2_g1__TRINITY_DN72995_c2_g1_i3__g.4661__m.4661/372470':0.3449060554997385):0.01695253875121061):0.05288939714513008):0.13204553157336907):0.09900263309861579):0.0736840117113422):0.04628600054668697):0.04597048127956338):0.10613357353680497):0.3107739474492375):0.1029489757392348; |
| CCD7 | 45705 | | | | | | | | | | Hormone metabolism (SL) | | | | |
| CCD8 | 45692 | | | | | | | | | | Hormone metabolism (SL) | | | | |
| CKX | 35681 | | | | | | | | | | Hormone metabolism (CK) | Angio | | | |
| D27L | 38300 | | | | | | | | | | Hormone metabolism (SL) | | | | |
| DAD1L | 35507 | | | | | | | | | | Hormone metabolism (JA) | | | | |
| EDS | 35811 | | | | | | | | | | Hormone metabolism (SA) | | | | |
| MoCo | 37513 | | | | | | | | | | Hormone metabolism (ABA) | | | Figure S1 | |
| OPCL | 35562 | | | | | | | | | | Hormone metabolism (JA) | | | | |
| OPR | 35662 | | | | | | | | | | Hormone metabolism (JA) | | | | |
| TAR | 36082 | | | | | | | | | | Hormone metabolism (IAA) | | | | |
| TPS | 35512 | | | | | | | | | | Hormone metabolism (GA) | | | | |
| YUC | 35647 | | | | | | | | | | Hormone metabolism (IAA) | Shared | | | |
| ZEP (ABA1L) | 39379 | | | | | | | | | | Hormone metabolism (ABA) | | | Figure S1 | |

| | | Details of main orthogroup | | | | Details of additional orthogroup | | | | | | | | | |
|---|---|---|---|---|---|---|---|---|---|---|---|---|---|---|---|
| **Gene family** | **Main orthogroup** | ***Arabidopsis thaliana*** | ***Hordeum vulgare*** | ***Polypodium vulgare*** | ***Ceratopteris richardii*** | **Additional orthogroup** | ***Arabidopsis thaliana*** | ***Hordeum vulgare*** | ***Polypodium vulgare*** | ***Ceratopteris richardii*** | **Function** | **Guard cell expression group** | **Notes** | **Related to Figure** | **Relevant clade subtree** |
| EPF1/2L | 42166 | | | | | | | | | | Stomatal patterning | | | | |
| EPFL9L | 42878 | | | | | | | | | | Stomatal patterning | | | | |
| EPFL1L | 35910 | | | | | | | | | | Stomatal patterning | | | | |
| ERECTA | 35631 | | | | | | | | | | Stomatal patterning | Angio | | | |
| KIN | 35946 | | | | | | | | | | Stomatal patterning | | | | |
| SCREAM | 36591 | | | | | | | | | | Stomatal patterning | | | | |
| SPCH/ MUTE/ FAMA | 35891 | | | | | | | | | | Stomatal patterning | Shared | only FAMA clade shown in Figure 2 | Figure 2 | (((Se_mo_91359/420354:0.77606641378039509238,((Po_vu_TRINITY_DN61025_c0_g1__TRINITY_DN61025_c0_g1_i4__g.156171__m.156171/352742:0.86100825949012649296,Ce_ri_TRINITY_DN43364_c1_g1__TRINITY_DN43364_c1_g1_i8__g.23332__m.23332/77156:0.73942904806745146651)92:0.48268526183051069234,Sa_cu_Sacu_v1.1_s0039.g012172/395014:1.05650164742948327046)79:0.45209206987570221825)37:0.17789630045627985444,((Se_mo_430716/427709:0.00000100000050002909,Se_mo_131087/427255:0.26768808877712868544)100:0.87791626505501063615,((Pi_ab_MA_57244g0010/288632:0.26545899693193864000,Gi_bi_Gb_32351/151975:0.10061289000624701329)96:0.21195620266765433093,(Am_tr_evm_27.model.AmTr_v1.0_scaffold00089.12/321:0.60300698477784742124,(Ho_vu_HORVU1Hr1G071330.3/195217:0.72700714927027576540,Ar_th_AT3G24140.1/35947:0.34550215357992614118)61:0.21075791739833071858)98:0.28082269591962621957)66:0.22219151563719971576)27:0.24510443573167792208)29:0.15090865523501095780,(Ph_pa_Pp3c19_19910V3.2.p/251416:0.41084662981231501178,Ph_pa_Pp3c22_14220V3.3.p/238567:0.23691691701137432280)100:1.24252227745082022281)31:0.07860462632993660270; |
| TMM | 38141 | | | | | | | | | | Stomatal patterning | | | | |

**Table S3. Gene sequence details.** Genes were sequenced during this study or identified either in the literature or by performing BLASTp searches using Arabidopsis protein sequences against the relevant genome or transcriptome assembly at Phytozome (v12; https://phytozome.jgi.doe.gov/; [S1]), ConGenIE (http://congenie.org/; [S2]), FernBase (https://fernbase.org; [S3]), TreeGenes (https://treegenesdb.org/; [S4, 5]), 1KP (https://db.cngb.org/onekp/; [S6]), or OrcAE (http://bioinformatics.psb.ugent.be/orcae/; [S7]), as indicated, and confirmed with reciprocal BLASTp searches back against Arabidopsis and preliminary phylogenetic analyses. Common name and/or family (for gymnosperms) is indicated in parentheses next to the species name.

| Plant group | Species | Gene name/ clade | Sequence ID | Sequence Source | Reference |
|---|---|---|---|---|---|
| Angiosperms | *Amborella trichopoda* | AmtrSLAC1 | AmTr_v1.0_scaffold00069.215 | v1.0, Phytozome, [S8] | [S9] |
| | | AmtrSLAH1 | AmTr_v1.0_scaffold00015.30 | | |
| | | AmtrSLAH2a | AmTr_v1.0_scaffold00075.12 | | |
| | | AmtrSLAH2b | XM_011624533.2 (corresponds to AmTr_v1.0_scaffold00075.14) | GenBank | |
| | | AmtrOST1 | AmTr_v1.0_scaffold00033.261 | v1.0, Phytozome, [S8] | [S10] |
| | | SnRK2 | AmTr_v1.0_scaffold00073.30 | | |
| | | | AmTr_v1.0_scaffold00017.69 | | |
| | | | AmTr_v1.0_scaffold00002.150 | | |
| | *Arabidopsis thaliana* | AtSLAC1 | AT1G12480 | TAIR10, Phytozome [S11] | [S12] |
| | | AtSLAH1 | AT1G62280 | | |
| | | AtSLAH2 | AT4G27970 | | |
| | | AtSLAH3 | AT5G24030 | | |
| | | AtSLAH4 | AT1G62262 | | |
| | | AtOST1/ AtSnRK2.6 | AT4G33950 | | [S13] |
| | | AtSnRK2.1 | AT5G08590 | | |
| | | AtSnRK2.2 | AT3G50500 | | |
| | | AtSnRK2.3 | AT5G66880 | | |
| | | AtSnRK2.4 | AT1G10940 | | |
| | | AtSnRK2.5 | AT5G63650 | | |
| | | AtSnRK2.7 | AT4G40010 | | |
| | | AtSnRK2.8 | AT1G78290 | | |
| | | AtSnRK2.9 | AT2G23030 | | |
| | | AtSnRK2.10 | AT1G60940 | | |
| | *Hordeum vulgare* (barley) | HvSLAC1 | AWG47875 | GenBank | [S14] |
| | | SLAH1/4 | HORVU3Hr1G033800 | r1, Phytozome [S15] | - |
| | | | HORVU2Hr1G118150 | | |
| | | | HORVU7Hr1G076220 | | |
| | | | HORVU5Hr1G066120 | | |
| | | | HORVU3Hr1G098700 | | |
| | | | HORVU4Hr1G076510 | | |
| | | SLAH2/3 | HORVU1Hr1G092640 | | [S14] |
| | | | HORVU3Hr1G030840 | | - |
| | | | HORVU5Hr1G055480 | | |
| | | | HORVU3Hr1G055740 | | [S16] |
| | | | HORVU1Hr1G030060 | | [S14] |
| | | | HORVU3Hr1G030850 | | [S14] |
| | | | HORVU3Hr1G055740 | | - |
| | | | HORVU3Hr1G005540 | | |
| | | | HORVU1Hr1G092740 | | |
| | | HvOST1.1 | HORVU4Hr1G013540 | | [S14] |
| | | HvOST1.2 | HORVU5Hr1G018340 | | |
| | | HvOST1.3 | HORVU5Hr1G097630 | | |
| | | HvOST1.4 | HORVU7Hr1G118150 | | |
| | | HvOST1.5 | HORVU2Hr1G029900 | | |
| | | SnRK2 | HORVU0Hr1G011570 | | - |
| | | | HORVU2Hr1G125950 | | |
| | | | HORVU4Hr1G087060 | | |
| | | | HORVU2Hr1G075470 | | |
| | | | HORVU1Hr1G055340 | | |
| | | | HORVU3Hr1G082690 | | |
| | | | HORVU1Hr1G074670 | | |
| | | | HORVU2Hr1G110230 | | |
| | | | HORVU2Hr1G029900 | | |
| Gymnosperms | *Amentotaxus argotaenia* (Taxaceae) | AmarSLAC1a | IAJW_scaffold_2007984 | 1KP [S6] | This study |
| | *Austrocedrus chilensis* (Cupressaceae) | AuchSLAC1a | YYPE_scaffold_2011657 | 1KP [S6] | This study |

| Plant group | Species | Gene name/ clade | Sequence ID | Sequence Source | Reference |
|---|---|---|---|---|---|
| | *Cephalotaxus harringtonia* (Taxaceae) | CehaSLAC1a | GJTI_scaffold_2060989 | 1KP [S6] | This study |
| | *Falcatifolium taxoides* (Podocarpaceae) | FataSLAC1a | ROWR_scaffold_2062445 | 1KP [S6] | This study |
| | *Ginkgo biloba* (Ginkgoaceae) | GbSLAC1a | MZ265387 | GenBank | This study |
| | | GbSLAC1b | Gb_39068 | http://gigadb.org/data set/100613 [S17] | |
| | | GbSLAC1c | Gb_19767 | | |
| | | GbSLAH2 | Gb_20865 | | |
| | | GbOST1 | Gb_02396 | | |
| | | SnRK2 | Gb_18878 | | |
| | | | Gb_11442 | | |
| | | | Gb_04315 | | |
| | | | Gb_15703 | | |
| | | | Gb_32504 | | |
| | *Gnetum montanum* (Gnetaceae) | GnmoSLAC1a | TnS000149361t06 | https://datadryad.org/ stash/dataset/doi:10. 5061/dryad.0vm37 [S18] | This study |
| | | GnmoSLAC1b | TnS000149361t07 | | |
| | | GnmoSLAC1c | TnS000149361t08 | | |
| | | GnmoSLAH1 | TnS000511471t06 | | |
| | | GnmoSLAH2 | TnS000686847t04 | | |
| | | | TnS000156791t21 | | |
| | | | TnS000142227t17 | | |
| | | | TnS000370521t01 | | |
| | | | TnS000949855t01 | | |
| | | | TnS000593659t01 | | |
| | | | TnS000380285t01 | | |
| | | SnRK2 | TnS000874347t07 | | |
| | | | TnS000086753t01 | | |
| | | | TnS000086753t03 | | |
| | | | TnS000088173t04 | | |
| | | | TnS000006441t06 | | |
| | | | TnS000419735t01 | | |
| | *Papuacedrus papuana* (Cupressaceae) | PapaSLAC1 | OVIJ_scaffold_2013792 | 1KP [S6] | This study |
| | *Picea abies* (Pinaceae) | PaSLAC1a | MZ265388 | GenBank | [S9] |
| | | PaSLAC1b | MZ265389 | | |
| | | PaSLAH1 | MZ265390 | | |
| | | PaSLAH2 | MZ265391 | | |
| | | PaOST1a | MA_10431614g0020 | v1.0, High Confidence Genes, ConGenIE [S2] | [S10] |
| | | PaOST1b | MA_10430788g0010 | | |
| | | SnRK2 | MA_10430272g0020 | | |
| | | | MA_19316g0010 | | |
| | | | MA_62786g0010 | | |
| | *Pilgerodendron uviferum* (Cupressaceae) | PiuvSLAC1a | ETCJ_scaffold_2011268 | 1KP [S6] | This study |
| | *Pinus taeda* (Pinaceae) | PitaSLAC1a | PITA_000008851 | v1.0, high quality whole genes, ConGenIE [S2, 19] | This study |
| | | PitaSLAC1b1 | PITA_000050611 | | |
| | | PitaSLAC1b2 | PITA_000013420 | | |
| | | PitaSLAC1c | PITA_000090181 | | |
| | | PitaSLAC1d | PITA_000093249 | | |
| | | PitaSLAH1a | PITA_000041418 | | |
| | | PitaSLAH1b | PITA_000041418 | | |
| | | PitaSLAH2a | PITA_000038302 | | |
| | | PitaSLAH2b | PITA_000051348 | | |
| | | PitaSLAH2c | PITA_000044861 | | |
| | | SnRK2 | PITA_000005625 | | |
| | | | PITA_000054419 | | |
| | | | PITA_000080600 | | |
| | | | PITA_000038518 | | |
| | | | PITA_000050071 | | |
| | | | PITA_000022758 | | |
| | *Pinus lambertiana* (Pinaceae) | PilaSLAC1b1 | PILA_00095 | V1.5, TreeGenes [S20] | This study |
| | | PilaSLAC1b2 | PILA_00097 | | |
| | | PilaSLAH1a | PILA_07555 | | |
| | | PilaSLAH1b | PILA_29112 | | |
| | | PilaSLAH2 | PILA_17561 | | |
| | | SnRK2 | PILA_01316 | | |
| | | | PILA_21884 | | |
| | | | PILA_20148 | | |
| | | | PILA_05349 | | |
| | | | PILA_08243 | | |
| | | | PILA_13153 | | |
| | *Pseudotsuga menziesii* (Pinaceae) | PsmeSLAC1a | PSME_17196 | v1.0, TreeGenes [S21] | This study |
| | | PsmeSLAC1b | PSME_24836 | | |
| | | PsmeSLAH2a | PSME_32370 | | |

| Plant group | Species | Gene name/ clade | Sequence ID | Sequence Source | Reference |
|---|---|---|---|---|---|
| | | PsmeSLAH2b | PSME_45884 | | |
| | | SnRK2 | PSME_29493 | | |
| | | | PSME_44831 | | |
| | | | PSME_12301 | | |
| | | | PSME_31121 | | |
| | | | PSME_33643 | | |
| | | | PSME_44283 | | |
| | | | PSME_22440 | | |
| | *Retrophyllum minus* (Podocarpaceae) | RemiSLAC1a | VGSX_scaffold_2072197 | 1KP [S6] | This study |
| | *Sequoiadendron giganteum* (Cupressaceae ) | SegiSLAC1a | SEGI_21902 | V2.0, TreeGenes [S22] | This study |
| | | SegiSLAC1b | SEGI_03395 | | |
| | | SegiSLAH1 | SEGI_37572 | | |
| | | SegiSLAH2 | SEGI_10075 | | |
| | | SegiOST1 | SEGI_02689 | | |
| | | SnRK2 | SEGI_03622 | | |
| | | | SEGI_37214 | | |
| | | | SEGI_12444 | | |
| | | | SEGI_03199 | | |
| | *Wollemia nobilis* (Araucariaceae) | WonoSLAC1a | RSCE_scaffold_2058598 | 1KP [S6] | This study |
| Ferns | *Azolla filiculoides* | AfSLAC1a | Azfi_s0064.g035597 | v1.2, FernBase; Li, et al. [S3] | [S23] |
| | | AfSLAC1b1 | Azfi_s0107.g045132 | | |
| | | AfSLAC1b2 | Azfi_s0107.g045136 | | |
| | | AfSLAC1b3 | Azfi_s0107.g045144 | | |
| | | AfSLAC1b4 | Azfi_s0107.g045141 | | |
| | | AfSLAC1c | Azfi_s0035.g025471 | | |
| | | AfSLAC1d1 | Azfi_s0019.g015208 | | |
| | | AfSLAC1d2 | Azfi_s1105.g097655 | | |
| | | AfSLAC1d3 | Azfi_s2247.g109987 | | |
| | | AfSLAC1e | Azfi_s0099.g044047 | | |
| | | AfSLAC1f | Azfi_s0099.g044050 | | |
| | | AfSLAC1g | Azfi_s0447.g070685 | | |
| | | SnRK2 | Azfi_s1324.g101449 | | [S10] |
| | | | Azfi_s0585.g078524 | | |
| | | | Azfi_s0538.g076300 | | |
| | | | Azfi_s0241.g059729 | | |
| | | | Azfi_s0022.g016063 | | |
| | | | Azfi_s0121.g046917 | | |
| | | | Azfi_s0137.g051038 | | |
| | | | Azfi_s0241.g059726 | | |
| | | | Azfi_s0059.g034604 | | |
| | *Ceratopteris richardii* | CrCPK | MZ265397 | GenBank | This study |
| | | CrSLAC1a | ANV22160 | | [S9] |
| | | CrSLAC1b | ANV22161 | | |
| | | CrSLAC1c | MZ265382 | | This study |
| | | CrSLAC1d | MZ265383 | | |
| | | CrSLAC1e | MZ265384 | | |
| | | CrSLAC1f | MZ265385 | | |
| | | CrSnRK2-1 | available in McAdam, et al. [S9] | | [S9] |
| | | CrSnRK2-2 | | | |
| | | CrSnRK2-3 | | | |
| | | CrSnRK2-4 | | | |
| | | CrSnRK2-5 | | | |
| | | CrSnRK2-6 | | | |
| | *Polypodium vulgare* | PvSLAC1a | MZ265379 | GenBank | This study |
| | | PvSLAC1b | MZ265380 | | |
| | | PvSLAC1c | TRINITY_DN76189_c2_g1 | www.stomatalevolution.org | |
| | | PvSLAC1d | TRINITY_DN75593_c0_g2 | | |
| | | PvSLAC1e | TRINITY_DN48986_c0_g1 | | |
| | | PvGAIA1 | MZ265381 | GenBank | |
| | | SnRK2 | TRINITY_DN74059_c9_g1 | www.stomatalevolution.org | |
| | | | TRINITY_DN69169_c0_g1 | | |
| | | | TRINITY_DN69990_c1_g1 | | |
| | | | TRINITY_DN58876_c0_g1 | | |
| | | | TRINITY_DN66878_c4_g1 | | |
| | | | TRINITY_DN66878_c3_g1 | | |
| | | | TRINITY_DN55509_c1_g1 | | |
| | | | TRINITY_DN80376_c2_g1 | | |
| | | | TRINITY_DN73955_c0_g1 | | |
| | *Salvinia cucullata* | ScSLAC1a | Sacu_v1.1_s0003.g001833 | v1.2, FernBase; Li, et al. [S3] | [S23] |
| | | ScSLAC1b | Sacu_v1.1_s0031.g010544 | | |
| | | ScSLAC1c | Sacu_v1.1_s0002.g000891 | | |
| | | ScSLAC1d | Sacu_v1.1_s0079.g017763 | | |
| | | ScSLAC1e | Sacu_v1.1_s0139.g022623 | | |
| | | ScSLAC1f | Sacu_v1.1_s0204.g025672 | | |
| | | ScSLAC1g | Sacu_v1.1_s0004.g002304 | | |

| Plant group | Species | Gene name/ clade | Sequence ID | Sequence Source | Reference |
|---|---|---|---|---|---|
| | | SnRK2 | Sacu_v1.1_s0061.g015371 | | [S10] |
| | | | Sacu_v1.1_s0258.g026859 | | |
| | | | Sacu_v1.1_s0024.g008972 | | |
| | | | Sacu_v1.1_s0183.g024944 | | |
| | | | Sacu_v1.1_s0075.g017417 | | |
| | | | Sacu_v1.1_s0004.g002142 | | |
| Lycophytes | *Selaginella moellendorffii* | SmSLAC1a1 | APA28903 | GenBank | [S9] |
| | | SmSLAC1a2 | APA28904 | | |
| | | SmSLAC1b | APA28905 | | |
| | | SmSLAC1c | APA28906 | | |
| | | SmSLAC1d | APA28907 | | |
| | | SmOST1a | 158991 | v1.0, Phytozome [S24] | [S9] |
| | | SmOST1b | 164978 | | |
| | | SmOST1c | 171183 | | |
| Mosses | *Sphagnum fallax* | SfSLAC1a | MZ265392 | GenBank | This study |
| | | SfSLAC1b | MZ265393 | | |
| | | SfSLAC2a | Sphfalx0007s0105 | v0.5, Phytozome, DOE-JGI (These sequence data were produced by the US Department of Energy Joint Genome Institute) | |
| | | SfSLAC2b | Sphfalx0159s0015 | | |
| | | SfSLAC2c | Sphfalx0004s0195 | | |
| | | SfSLAH | Sphfalx0269s0010 | | |
| | | SfOST1.1 | MZ265394 | GenBank | |
| | | SfOST1.2 | MZ265395 | | |
| | | SfOST1.3 | MZ265396 | | |
| | *Physcomitrella patens* | PpSLAC1 | Pp3c1_33890V3.1 | v3.3, Phytozome [S25, 26] | [S27] |
| | | PpSLAC2 | Pp3c2_26550V3.1 | | |
| | | PpSLAH1 | Pp3c14_15640V3.1 | | |
| | | PpSLAH2 | Pp3c9_17220V3.1 | | |
| | | PpOST1.1/ PpSnRK2a | Pp3c5_21160V3.1 | | [S27-30] |
| | | PpOST1.2/ PpSnRK2b | Pp3c6_16600V3.1 | | |
| | | PpOST1.3/ PpSnRK2c | Pp3c6_11090V3.1 | | |
| | | PpOST1.4/ PpSnRK2d | Pp3c5_17150V3.1 | | |
| Liverworts | *Marchantia polymorpha* | MpSLAC1 | Mapoly0073s0030 | v3.1, Phytozome [S27] | [S27] |
| | | MpSLAC2 | Mapoly0073s0098 | | [S23] |
| | | MpOST1.1/ MpSNRK2A | Mapoly0061s0075 | | [S27] |
| | | MpOST1.2/ MpSNRK2B | Mapoly0011s0096 | | |
| Hornworts | *Anthoceros agrestis* | AaSLAC1 | MZ265386 | GenBank | [S31]; This study |
| | | AaOST1 | AagrBONN_evm.model.Sc2ySwM_228.3243.1 | https://figshare.com/articles/Genome_assemblies_and_annotations_of_the_three_Anthoceros_accessions_as_well_as_alignment_matrices_and_tree_files_used_for_reconstructing_the_land_plant_phylogeny_/9974999 [S31] | |
| | *Anthoceros angustus* | AangSLAC1 | AANG000829 | https://doi.org/10.5061/dryad.msbcc2ftv [S32] | This study |
| | | AangOST1 | AANG006294 | | |
| | *Anthoceros punctatus* | ApunSLAC1 | Apun_evm.TU.utg000148l.40 | https://www.hornworts.uzh.ch/en.html [S31] | This study |
| | | ApunOST1 | Apun_evm.model.utg000043l.1191.1 | | |
| Charophytic algae | *Chara braunii* | | No BLASTp or tBLASTn hits for AtSLAC1 | Chbra.pep.20180417/chara_genome [S33], OrcAE | [S33]; This study |
| | | SnRK2 | CbG22934 | | |
| | *Chlorokybus atmophyticus* | | No BLASTp or tBLASTn hits for AtSLAC1 | https://db.cngb.org/search/assembly/CNA0002353/ [S34] | [S34];This study |
| | | SnRK2 | Chrsp90S08126 | | |
| | | | Chrsp32S08952 | | |
| | *Klebsormidium nitens* | KnSLAC1 | CEO16428.1 | GenBank | [S27] |
| | *Mesostigma viride* | MeviSLAC1 | Mesvi144S02876 | https://db.cngb.org/search/assembly/CNA0002352/ [S34] | [S34]; This study |
| | | SnRK2 | Mesvi2102S04184 | | |
| | | | Mesvi1159S00983 | | |
| | *Mesotaenium endlicherianum* | | No BLASTp or tBLASTn hits for AtSLAC1 | https://figshare.com/articles/Genomes_of_subaerial_Zygnematophyceae_provide_insights_into_land_plant | [S35]; This study |
| | | SnRK2 | ME000293S04937 | | |

| Plant group | Species | Gene name/ clade | Sequence ID | Sequence Source | Reference |
|---|---|---|---|---|---|
| | | | | _evolution/9911876/1 [S35] | |
| | *Penium margaritaceum* | | No BLASTp or tBLASTn hits for AtSLAC1 | http://bioinfo.bti.cornell.edu/cgi-bin/Penium/download.cgi [S36] | [S36]; This study |
| | | SnRK2 | pm004004.t1 | | |
| | | | pm018762g0010 | | |
| | | | pm003108g0050 | | |
| | *Prasinoderma coloniale* | | No BLASTp or tBLASTn hits for AtSLAC1 | https://db.cngb.org/search/project/CNP0000924/ [S37] | [S37]; This study |
| | | SnRK2 | PRCOL_00007272-RA | | |
| | *Spirogloea muscicola* | | No BLASTp or tBLASTn hits for AtSLAC1 | https://figshare.com/articles/Genomes_of_subaerial_Zygnematophyceae_provide_insights_into_land_plant_evolution/9911876/1 [S35] | [S35]; This study |
| | | SnRK2 | SM000086S23047 | | |
| | | | SM000002S05533 | | |
| | | | SM000015S01198 | | |
| | | | SM000035S13104 | | |
| | | | SM000008S22152 | | |
| | | | SM000086S23041 | | |
| | | | SM000002S05527 | | |
| | | | SM000015S01193 | | |

**Table S4. Overview of transcriptomes and differential gene expression analysis.** Significance cutoff for guard cell genes: padj <= 0.01 and logfold < 0.

| Species | Unigenes | Isoforms | Protein coding (Uni)genes | Guard Cell protein coding genes |
|---|---|---|---|---|
| *P. vulgare* | 293847 | 635910 | 50833 | 8333 |
| *C. richardii* | 165371 | 347757 | 30260 | 1900 |
| *A. thaliana* | | | 27628 | 1826 |
| *H. vulgare* | | | 37673 | 6653 |

**Table S5. Genomic data used for gene expression analysis and evolutionary reconstruction.**

| Species | Data sets | References |
|---|---|---|
| *Amborella trichopoda* | Atrichopoda_291_v1.0.protein.fa.gz | [S8] |
| *Arabidopsis thaliana* | Genome: Arabidopsis_thaliana.TAIR10.dna.toplevel.fa.gz<br>Proteome: Arabidopsis_thaliana.TAIR10.pep.all.fa | [S11] |
| *Chara braunii* | mRNA_Chbra_active_pep_20170414.tfa.xz | [S34] |
| *Ginkgo biloba* | Gb.pep.fa | [S38] |
| *Hordeum vulgare* | Genome: Hordeum_vulgare.IBSC_v2.dna.toplevel.fa.gz<br>Proteome: Hordeum_vulgare.Hv_IBSC_PGSB_v2.pep.all.fa | [S39] |
| *Klebsormidium nitens* | 160614_klebsormidium_v1.1_AA.fasta | [S40] |
| *Marchantia polymorpha* | Mpolymorpha_320_v3.1.protein.fa.gz | [S41] |
| *Picea abies* | Pabies1.0-all-pep.faa.gz | [S2] |
| *Physcomitrella patens* | Ppatens_318_v3.3.protein.fa.gz | [S26] |
| *Salvinia cucullata* | Salvinia_cucullata.protein.highconfidence_v1.2.fasta | [S3] |
| *Selaginella moellendorffii* | Smoellendorffii_91_v1.0.protein.fa.gz | [S24] |

**Table S6. Primer details.**

| Gene | Purpose | Primer sequences (5' to 3') |
|---|---|---|
| *AaSLAC1* | Cloning full length CDS without stop codon | F:GGCTTAAUATGAGCAGTAATAGGCCGGCGGGGTC |
| | | R1:GGTTTAAUCCTACGCCGTTTGAAGTGACG |
| | Cloning full length CDS with stop codon | F:GGCTTAAUATGAGCAGTAATAGGCCGGCGGGGTC |
| | | R2:GGTTTAAUTTATACGCCGTTTGAAGTGA |
| *AmtrOST1* | Cloning full length CDS without stop codon | F:GGCTTAAUATGGATCGGACTGCTCTCAC |
| | | R1:GGTTTAAUCCCATTGCATAGACGATCTCACCAC |
| | Cloning full length CDS with stop codon | F:GGCTTAAUATGGATCGGACTGCTCTCAC |
| | | R2:GGTTTAAUTTACATTGCATAGACGATCTCACC |
| *AmtrSLAC1* | Cloning full length CDS without stop codon | F1:GGCTTAAUATGAAGCCAGATAAATATTGCAG |
| | | R1:GGTTTAAUCCGACCTTTCCTCCACTTGCAT |
| | Cloning full length CDS with stop codon | F1:GGCTTAAUATGAAGCCAGATAAATATTGCAG |
| | | R2:GGTTTAAUTCAGACCTTTCCTCCACTTG |
| | Introducing Y481L mutation | F2:ACTTAGCCAUTGCAATAACAAGACACA +R1/R2, |
| | | F1+R3:ATGGCTAAGUCATTTGGGAACAATGAGCC |
| *CrCPK* | Cloning full length CDS without stop codon | F:GGCTTAAUATGGGTAACTGCTGCAGCAA |
| | | R1:GGTTTAAUCCCTTCTGTAGTGAGCCATCTTTC |
| | Cloning full length CDS with stop codon | F:GGCTTAAUATGGGTAACTGCTGCAGCAA |
| | | R2:GGTTTAAUCTACTTCTGTAGTGAGCCATCTT |
| *CrSLAC1c* | Cloning full length CDS without stop codon | F:GGCTTAAUATGGCAAGAGAAGATTGCAAAG |
| | | R1:GGTTTAAUCCGCCTTGGGTCCATGCGAGCTG |

| Gene | Purpose | Primer sequences (5' to 3') |
|---|---|---|
| | Cloning full length CDS with stop codon | F:GGCTTAAUATGGCAAGAGAAGATTGCAAAG |
| | | R2:GGTTTAAUTTAGCCTTGGGTCCATGC |
| *CrSLAC1d* | Cloning full length CDS without stop codon | F:GGCTTAAUATGCCTTCATCAGCGCGTTC |
| | | R1:GGTTTAAUCCTACTGGTAGGTCATTTTCCT |
| | Cloning full length CDS with stop codon | F:GGCTTAAUATGCCTTCATCAGCGCGTTC |
| | | R2:GGTTTAAUTCATACTGGTAGGTCATTTTCCT |
| *CrSLAC1e* | Cloning full length CDS without stop codon | F:GGCTTAAUATGGGATATGTATCCAATATTGAAG |
| | | R1:GGTTTAAUCCGATATAACATGGTAGAAAATCTTT |
| | Cloning full length CDS with stop codon | F:GGCTTAAUATGGGATATGTATCCAATATTGAAG |
| | | R2:GGTTTAAUTTAGATATAACATGGTAGAAAATC |
| *CrSLAC1f* | Cloning full length CDS without stop codon | F:GGCTTAAUATGTATCCATACATGAAGGA |
| | | R1:GGTTTAAUCCTTTTGCGCCCCTCTGTGA |
| | Cloning full length CDS with stop codon | F:GGCTTAAUATGTATCCATACATGAAGGA |
| | | R2:GGTTTAAUCTATTTTGCGCCCCTCTG |
| *GbSLAC1a* | Cloning full length CDS without stop codon | F:GGCTTAAUATGGACACCAAAATCGAAAAA |
| | | R1:GGTTTAAUCCTCTTGTTTGTTTCGTAAAGC |
| | Cloning full length CDS with stop codon | F:GGCTTAAUATGGACACCAAAATCGAAAAA |
| | | R2:GGTTTAAUTTATCTTGTTTGTTTCGTAAAGC |
| *PaSLAC1a* | Cloning full length CDS without stop codon | F1:GGCTTAAUATGAATCCCATAAGACCTGCAA |
| | | R1:GGTTTAAUCCTGGCAATTTATGGTTGAAGAGA |
| | Cloning full length CDS with stop codon | F1:GGCTTAAUATGAATCCCATAAGACCTGCAA |
| | | R2:GGTTTAAUTCATGGCAATTTATGGTTGAA |
| | Introducing GbS120 mutation | F2:CTTGACAAACAGAAuTCcCTGCTGCAAGCCCG+R1/R2, |
| | | F1+R3:GGAATTCTGTTTGuCAAgACCAGACTTGGTCCCGAA |
| | Introducing F512A mutation | F3:ACAgcCCCAAuGACTGCAGCAGCAGTGGC+R1/R2, |
| | | F1+R4:ATTGGGgcTGuATATGCCCACCAAGCT |
| *PaSLAC1b* | Cloning full length CDS without stop codon | F:GGCTTAAUATGGAAAACCAGAACTCTTTCA |
| | | R1:GGTTTAAUCCATCTAGGGGACCATGACCAG |
| | Cloning full length CDS with stop codon | F:GGCTTAAUATGGAAAACCAGAACTCTTTCA |
| | | R2:GGTTTAAUTTAATCTAGGGGACCATGACCA |
| *PaSLAH1* | Cloning full length CDS without stop codon | F:GGCTTAAUATGTCGCAGATAAGCATGAGC |
| | | R1:GGTTTAAUCCCTGAAAATTAGCTGGTGGGTTT |
| | Cloning full length CDS with stop codon | F:GGCTTAAUATGTCGCAGATAAGCATGAGC |
| | | R2:GGTTTAAUTCACTGAAAATTAGCTGGTG |
| *PaSLAH2* | Cloning full length CDS without stop codon | F:GGCTTAAUATGGAGTCAATTGAAATCACCA |
| | | R1:GGTTTAAUCCGAGAGATAATTCAGAGAGTAAAGT |
| | Cloning full length CDS with stop codon | F:GGCTTAAUATGGAGTCAATTGAAATCACCA |
| | | R2:GGTTTAAUTTAGAGAGATAATTCAGAGAGTAAA |
| *PvGAIA1* | Cloning full length CDS without stop codon | F:GGCTTAAUATGGATCGCGTTGTTGCGGGTGCT |
| | | R1:GGTTTAAUCCAATAGCACTCACATACTCTCCA |
| | Cloning full length CDS with stop codon | F:GGCTTAAUATGGATCGCGTTGTTGCGGGTGCT |
| | | R2:GGTTTAAUTCAAATAGCACTCACATACTCTC |
| *PvSLAC1a* | Cloning full length CDS without stop codon | F1:GGCTTAAUATGGCAACATATGGTGCGC |
| | | R1:GGTTTAAUCCTGCATCTCTGTCGATAGAATTTTCC |
| | Cloning full length CDS with stop codon | F1:GGCTTAAUATGGCAACATATGGTGCGC |
| | | R2:GGCTTAAUTCATGCATCTCTGTCGATAGAATTTT |
| | Introducing V663L mutation | F2:ATcTCGCCAUTGCCATAACAAGGAAGAAGCA+R1/R2, |
| | | F1+R3:ATGGCGAgAUCATTCGGAAACAAACTGCCC |
| *PvSLAC1b* | Cloning full length CDS without stop codon | F:GGCTTAAUATGCGAAACAACGGTGCA |
| | | R1:GGTTTAAUCCTGCTACACCCTCTTGAAGC |
| | Cloning full length CDS with stop codon | F:GGCTTAAUATGCGAAACAACGGTGCA |
| | | R2:GGTTTAAUTCATGCTACACCCTCTTGAA |
| *SfOST1-1* | Cloning full length CDS without stop codon | F:GGCTTAAUATGGACCCCTTGGAGATCA |
| | | R1:GGTTTAAUCCTATAGCGCACACAAACTCCCCACT |
| | Cloning full length CDS with stop codon | F:GGCTTAAUATGGACCCCTTGGAGATCA |
| | | R2:GGTTTAAUTTATATAGCGCACACAAACTCC |
| *SfOST1-2* | Cloning full length CDS without stop codon | F:GGCTTAAUATGGACTTTTCGAGTATACAAGATGT |
| | | R1:GGTTTAAUCCTATGGCACACACAAATTCCCCACT |
| | Cloning full length CDS with stop codon | F:GGCTTAAUATGGACTTTTCGAGTATACAAGATGT |
| | | R2:GGTTTAAUTTATATGGCACACACAAATTCC |
| *SfOST1-3* | Cloning full length CDS without stop codon | F:GGCTTAAUATGGATTTGTTTTCTGGGATA |
| | | R1:GGTTTAAUCCTATAGCACACACAAATTCCCCACT |
| | Cloning full length CDS with stop codon | F:GGCTTAAUATGGATTTGTTTTCTGGGATA |
| | | R2:GGTTTAAUTTATATAGCACACACAAATTCCCCACT |
| *SfSLAC1a* | Cloning full length CDS without stop codon | F:GGCTTAAUATGGACTCTGCTGTTGCTGTT |
| | | R1:GGTTTAAUCCAGCTTCAAGAGGCTGATCAGGAAG |
| | Cloning full length CDS with stop codon | F:GGCTTAAUATGGACTCTGCTGTTGCTGTT |
| | | R2:GGTTTAAUTCAAGCTTCAAGAGGCTGAT |
| *SfSLAC1b* | Cloning full length CDS without stop codon | F:GGCTTAAUATGGCAGCAGAGTTGCGACGGGAGGT |
| | | R1:GGTTTAAUCCACCTCTAGAAGAGGTCACAC |
| | Cloning full length CDS with stop codon | F:GGCTTAAUATGGCAGCAGAGTTGCGACGGGAGGT |
| | | R2:GGTTTAAUTATGCGCTGCTGACAACTTT |

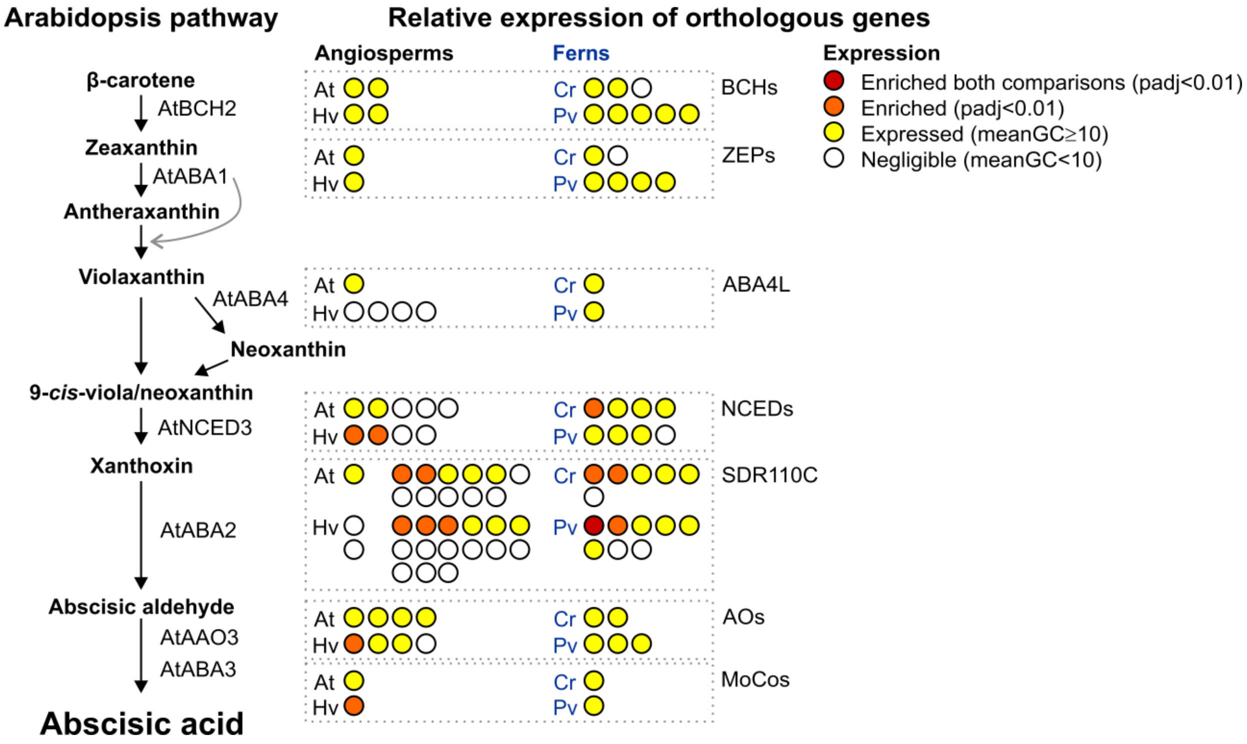


**Figure S1. Relative expression of orthogroups containing abscisic acid (ABA) biosynthesis pathway components.** The Arabidopsis biosynthesis pathway is shown on the left with ABA precursors shown interconnected by arrows, and the proteins that catalyse each step indicated. Orthogroups/subclades containing the genes that encode these proteins are shown together in boxes on the right with each circle representing a different gene and colour coding representing relative expression in guard cell-enriched samples compared to whole leaves is shown for angiosperm (At, *Arabidopsis thaliana*; Hv, *Hordeum vulgare* barley) and fern (Cr, *Ceratopteris richardii*; Pv, *Polypodium vulgare*) models as indicated. For *P. vulgare* samples only, whole leaf vs 'leaf samples without abaxial epidermis' (thus guard cells) removed were also included and used to separate guard cell-enriched genes with a higher level of stringency (red; "enriched both comparisons" = expression higher in guard cells than leaves, and higher in whole leaves than leaves without guard cells). Please note that the ABA2 clade of SDRs is only found in angiosperms [S42, 43], but based on the leaky nature of *aba2* mutants [S44, 45], other related SDRs (e.g. within the same SDR110C clade) are likely to be able to catalyse this step of ABA biosynthesis; Arabidopsis and barley ABA2 orthologs (left; determined phylogenetically) are shown separately to other SDR110C clade members for clarity. See Table S2 for gene details.

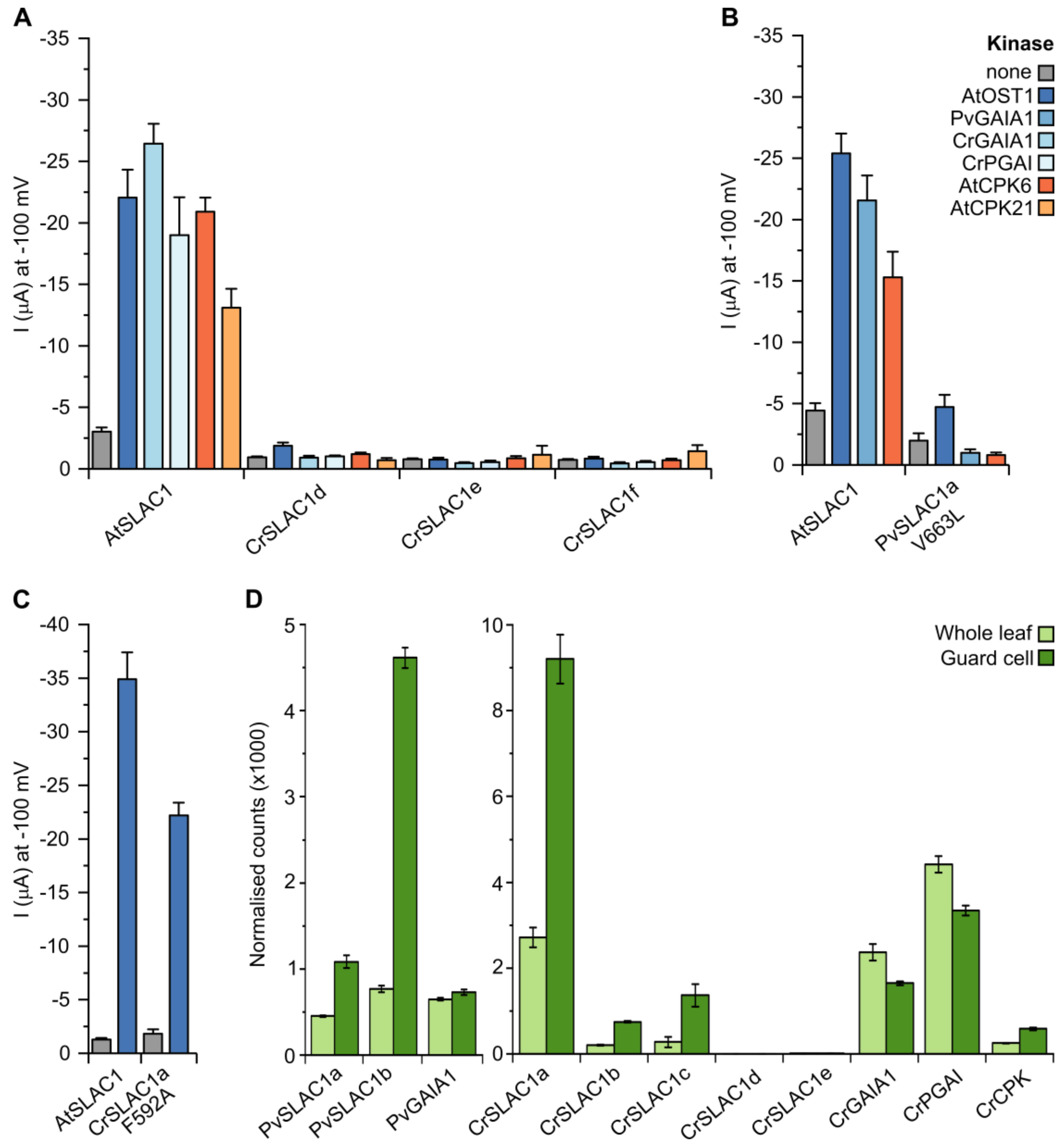


**Figure S2. Additional data for fern SLAC homolog activity and expression**. **(A-B)** Mean whole-oocyte current measurements at −100 mV in nitrate-based standard medium of **(A)** wild-type SLACs from the angiosperm *A. thaliana* and the fern *C. richardii*, **(B)** a mutant of PvSLAC1a in which the C-terminal motif is altered to match Arabidopsis, and (C) a gate mutant of CrSLAC1a, co-expressed with or without kinases in *Xenopus* oocytes (mean ± SEM, n ≥ 3). **(D)** Expression of *P. vulgare* and *C. richardii* genes of interest in whole leaf and guard cell-enriched samples. Please note that CrSLAC1f was not present in our transcriptome. Counts were normalised by sample-specific size factors determined by median ratio of gene counts relative to geometric mean per gene.

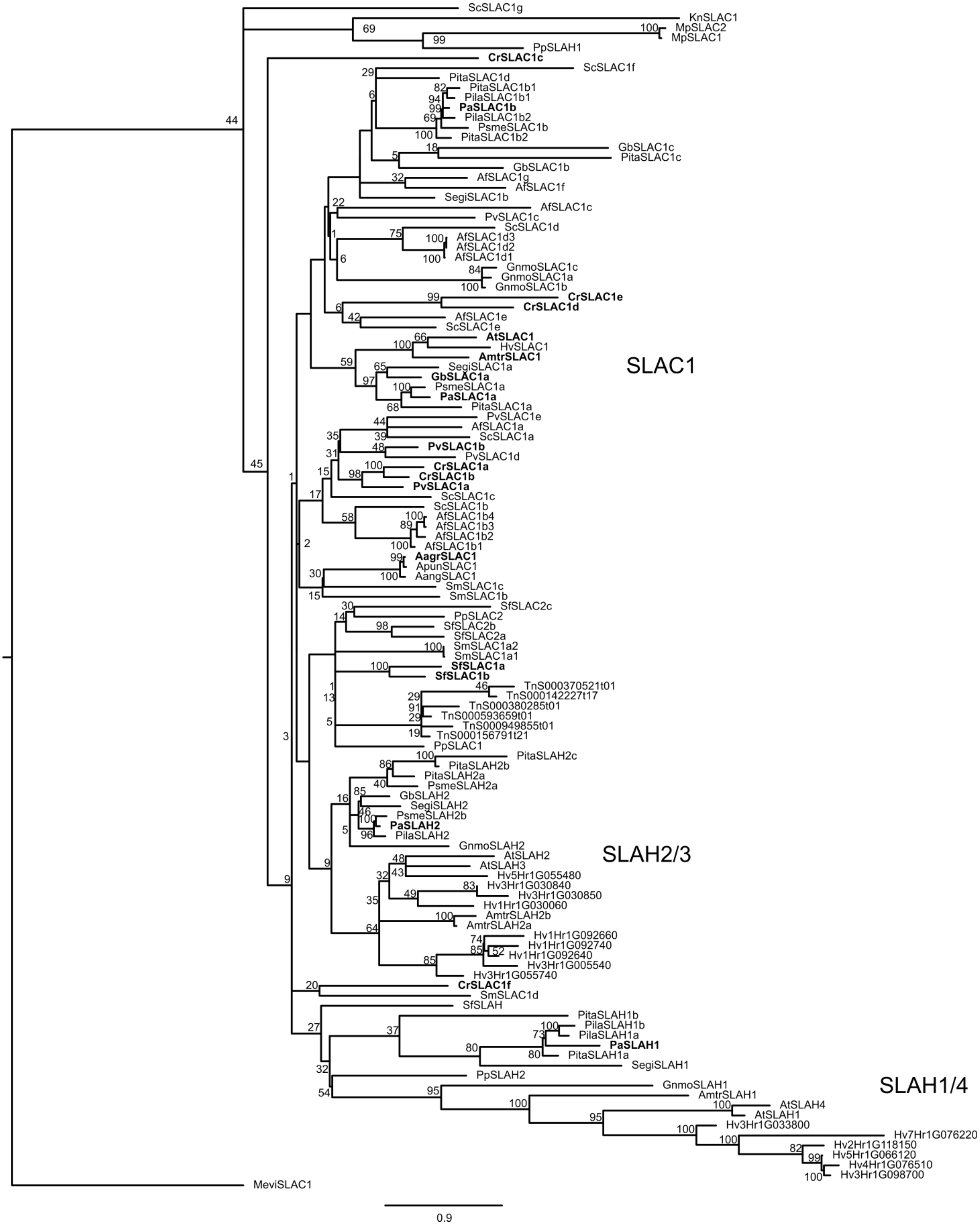


**Figure S3. Phylogeny of the SLAC/SLAH family in streptophytes**. SLAC/SLAH channels measured in *Xenopus* oocytes in this study are shown in bold. Bootstrap values from 1000 replicates are shown as percentages on nodes of the maximum likelihood phylogenetic tree. Full sequence and species details are given in Table S3.

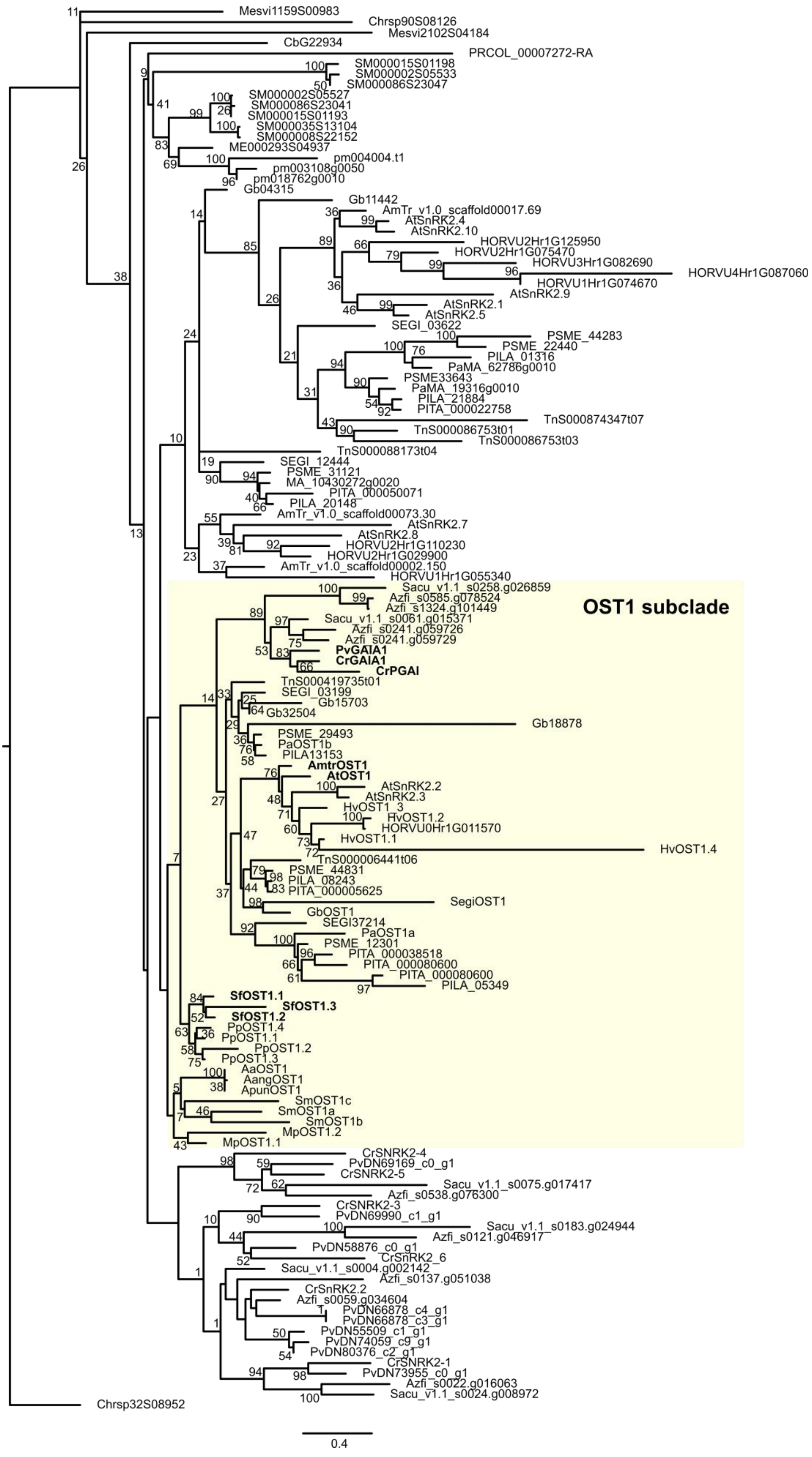


**Figure S4. Phylogeny of the SnRK2 family in streptophytes.** Kinases from the OST1 subclade measured in *Xenopus* oocytes in this study are shown in bold. Bootstrap values from 1000 replicates are shown as percentages on nodes of the maximum likelihood phylogenetic tree. Full sequence and species details are given in Table S3.

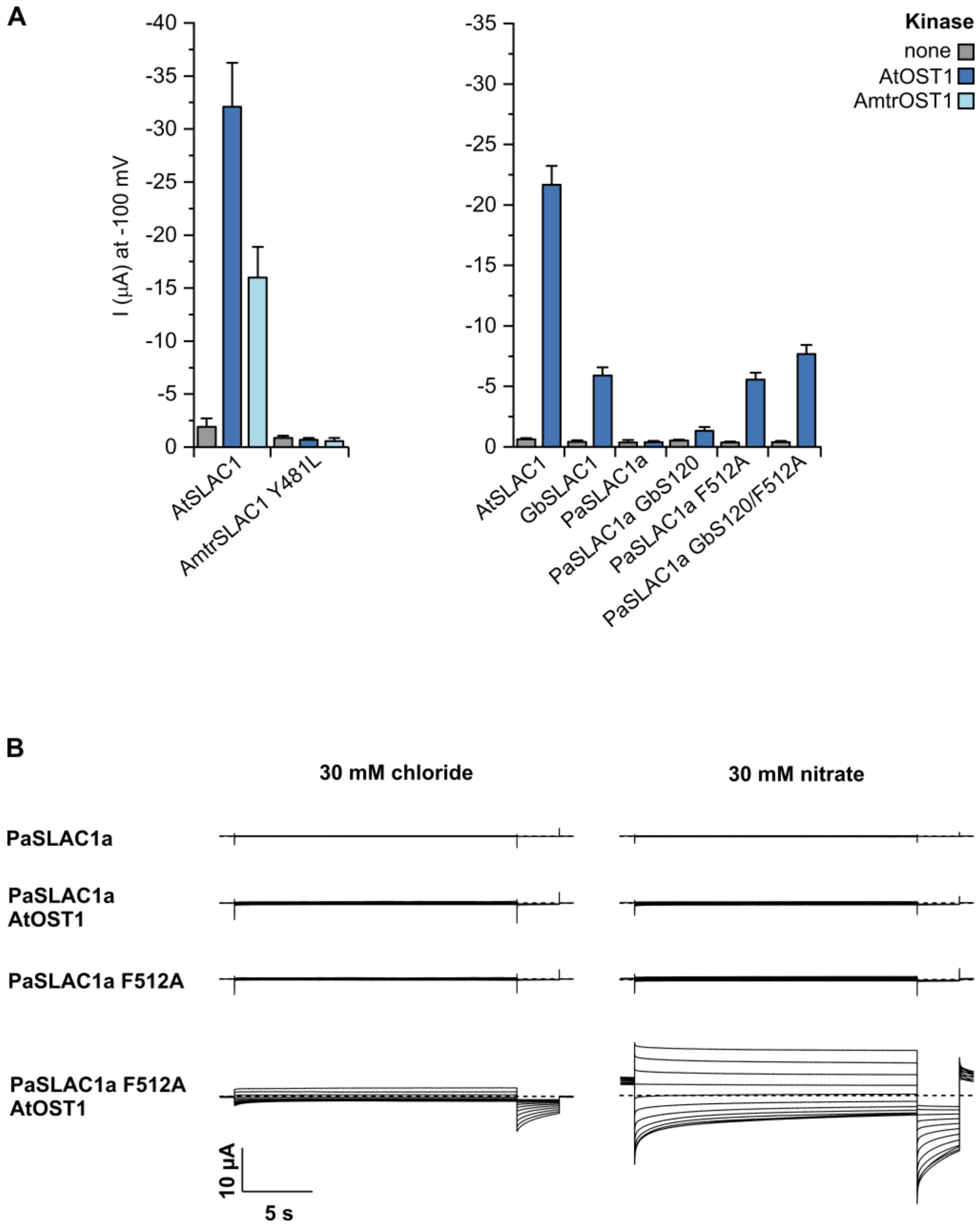


**Figure S5. Additional data for seed plant SLAC1 orthologs. (A)** Mean whole-oocyte current measurements at −100 mV in nitrate-based standard medium with SLAC/SLAH channels from the angiosperms *A. thaliana* and *A. trichopoda*, and the gymnosperms *P. abies* and the *G. biloba*, including the Y481L mutant created to match the AtSLAC1 C-terminal motif (see **Fig. 5**), the gate mutant F512A and the GbS120 mutant created to match the GbSLAC1 S120 region (see **Fig. 5**), co-expressed with or without kinases in *Xenopus* oocytes (mean ± SEM, n ≥ 4). **(B)** Example whole-oocyte currents of PaSLAC1a wild-type and F512A gate mutant, either expressed alone or with AtOST1, recorded in chloride and nitrate bath solutions. Voltage pulses lasting 20 s ranging from +40 to −180 mV in 20 mV decrements were applied (holding potential $V_H$ was 0 mV).
